# Uncertainties of cell number estimation in cyanobacterial colonies and the potential use of sphere packing

**DOI:** 10.1101/2022.02.13.480236

**Authors:** Enikő T-Krasznai, Verona Lerf, István Tóth, Tibor Kisantal, Gábor Várbíró, Gábor Vasas, Viktória B-Béres, Judit Görgényi, Áron Lukács, Zsuzsanna Kókai, Gábor Borics

## Abstract

Cyanobacteria are notorious bloom formers causing various water quality concerns, such as toxin production, extreme diurnal variation of oxygen, or pH, etc., therefore, their monitoring is essential to protect the ecological status of aquatic systems. Cyanobacterial cell counts and biovolumes are currently being used in water management and water quality alert systems. In this study, we investigated the accuracy of colonial biovolume and cell count estimation approaches used in the everyday practice. Using real like 3-dimensional images of cyanobacterial colonies, we demonstrated that their shape cannot be approximated by ellipsoids. We also showed that despite the significant relationship between overall colony volume and cell biovolumes because of the considerable scatter of cell count data the regressions give biased estimates for cyanobacterial cell counts. We proposed a novel approach to estimate cell counts in colonies that was based on the random close sphere packing method. This method provided good results only in those cases when overall colony volumes could be accurately measured. The visual investigation of colonies done by skilled experts has given precise but lower estimates for cell counts. Estimation results of several experts were surprisingly good which suggest that this capability can be improved, and estimation bias can be reduced to the level acceptable for water quality estimations.

**Highlights:**

Cyanobacterial colony – cell biovolume relationships provide biased estimates for cellbiovolumes.
Sphere packing approach provides good cellcount estimates if colony volumes are accurately measured.
Considering cyanobacterial colonies as ellipsoids gives inaccurate volume estimates.
Skilled experts slightly underestimate the cellcounts but dispersion of their estimates is low.

## Introduction

Despite the efforts that have been made to reduce the nutrient loads to surface waters, eutrophication remains a major source of concerns for water management (Smith & Schindler, 2009). The process coincides with drastic increase in phytoplankton biomass and undesirable compositional changes, reducing the stability of aquatic ecosystems and threatening the services they provide to society. Monitoring the quality of surface waters therefore is essential to avoid the negative consequences of eutrophication.

Water blooms, which can be caused by cyanobacteria and several eukaryotic algae (Reynolds & Walsby, 1975) are the most serious problems in eutrophic waters worldwide. While some of the eukaryotic algae are capable to form freshwater red tides (e.g. *Uroglenopsis americana*; Ochrophyta) (Ishikawa et al., 2005), others produce surface scums (e.g. *Botryococcus braunii*; Chlorophyta) (Paerl et al., 2001). The formed blooms can be skin irritating (*Gonyostomum semen*; Ochrophyta) (Hongve et al., 1988), or toxic (e.g. *Prymnesium parvum*; Haptophyta) (Vasas et al., 2012). Cyanobacteria as one of the most notorious groups of phytoplankton, are responsible for the vast majority of harmful algal blooms throughout the world (Pearl et al., 2001). Cyanobacterial blooms, apart from being aesthetically displeasing, can be toxic, and can decrease water quality causing health problems both for animals and humans, and thus, reduce water use (Carvalho et al., 2013). Because of the globally increasing occurrence of cyanobacterial harmful algal blooms (CyanoHABs), several methods have been developed for estimating the biomass of cyanobacteria (e.g. qPCR, chlorophyll-a concentration, concentration of microcystin, ESP; Alcántara et al., 2018; Seltenrich, 2014; Wang et al., 2015; *in vivo* phycocyanin fluorescence, McQuaid et al,, 2011). However, giving the accepted thresholds for cell counts (WHO, 1999, 2003) requires the use of traditional microscopic analyses of samples with identification of taxa and estimation of cell counts and taxon-specific biovolumes (Carvallho et al., 2013).

Biovolume estimation of phytoplankton is based on the microscopy measurements of linear dimensions of the units that are compared to geometric shapes (Hillebrand et al., 1999). Multiplying these values with volumetric cell counts (cells mL^-1^) provides species-biovolume data, which serves as a basis for both scientific and management purposes (CEN 16695, 2015), This approach provides reasonable estimates for specific biovolumes in the case of simple-shaped unicellular taxa and multicellular filaments but for complex shapes it might give uncertain results (Borics et al., 2021).

Based on their morphological appearances, cyanobacteria can be distinguished into three major groups: i) unicellular, ii) filamentous, and iii) colony forming. While measuring the cell sizes and numbers of unicells and filaments (considering them as cylinders) can be easily accomplished, estimation of cell numbers in colonial forms has an unknown uncertainty of accuracy, especially in the case of coenobial (*Aphanocapsa, Coelomoron, Microcystis* or *Pannus*) and utricular (*Coelosphaerium, Snowella* or *Woronichinia*) forms. However, the problem is not new, and during the last decades, some alternative approximations and procedures were proposed to estimate the cell number per colony. In the case of most methods, disintegration of colonies to single cells is always the first step, which can be carried out by sonication, heating, boiling or using alkaline hydrolysis (Box, 1981). Second step is to count the cell numbers per sample, which can be carried out by hemocytometer (Joung et al., 2006), or FlowCAM (Wang et al., 2015). These methods however are proposed to use in the case of blooms with the dominance of *Microcystis* spp. Reynolds and Jaworski (1978) estimated the concentration of cells per colony using a regression equation fitted to data derived from natural populations. Joung et al (2006) applied a similar approach; after disintegration of *Microcystis* colonies by boiling, they counted the cell numbers and calculated their concentrations in the colonies, which had been measured previously and were considered as spheres. The authors found a clear relationship between colony size and *Microcystis* cell numbers.

Alcántara et al (2018) estimated the colony volume of *Microcystis aeruginosa* complex’ using a geometrical approximation. Three theoretical morphologies of the colonies (sphere, prolate spheroid and ellipsoid) were compared, and the authors propose to apply ellipsoid forms to estimate colonial biovolume of *Microcystis* spp. This approach can be useful to estimate colony size in routine monitoring, but unfortunately does not provide any help in measuring cellular biovolumes.

Recently an increasing demand for monitoring of cyanobacterial blooms is emerging, because the climate change and the increasing anthropogenic pressures trigger the bloom development both in eutrophic (Huisman et al., 2018) and oligotrophic (Reinl et al., 2021) environments.

In this study we aim to investigate the accuracy of the existing approaches and of a new one that merges the advantages of 3-D imagery of microalgae (Borics et al., 2021) and the practical application of the age-old sphere packing problem (Hifi & M’hallah, 2009).

We hypothesised that colony volume and cell count estimation together give a better prediction for colonial cyanobacterial biovolume than the aforementioned regression approaches.

## Material and methods

### Sampling

To assess the accuracy of the above mentioned approaches we collected phytoplankton samples from various Hungarian standing waters in 2020. Samples were collected from the photic zone of the waters and preserved by Lugol’s solution in the field. Samples were stored at 4 °C until the analyses.

### Sample processing and giving the reference cellcounts

We investigated the colonies by Leica DMRB light microscope at 100× and 400× magnifications. Photos were taken by Canon 4000D camera and the colonies were measured by QuickPHOTO CAMERA 3.2 program.

The studied colonies were randomly selected in the droplets. In order to know the real shape of the colonies we preferred those that could be rotated under the cover glass. We took several photos of these colonies in different layers, and these were used later to measure the colony size (length, width and depth), the cell size and then the distances of cells within the colonies. To estimate the mean distances among the cells a minimum of 20 measurements were taken. The mucilage of the colony was not measured, only the space filled by cells. In order to know the exact number of cells within the colony, the water was carefully desiccated from under the cover glass, resulting in flattening of the colonies until the cells lied in one layer (Fig. 1.). Finally, we took a photo of the flattened colonies and counted the cells in the photographs (shown in electronic supplementary material, Fig. S1). Counted cells were marked by colour dots using Microsoft Paint. These cell numbers were considered as references during comparison of the three cell count estimation approaches shown below. Based on the measurements on the cells, we counted cell volumes and overall cell biovolumes.

**Fig. 1.**
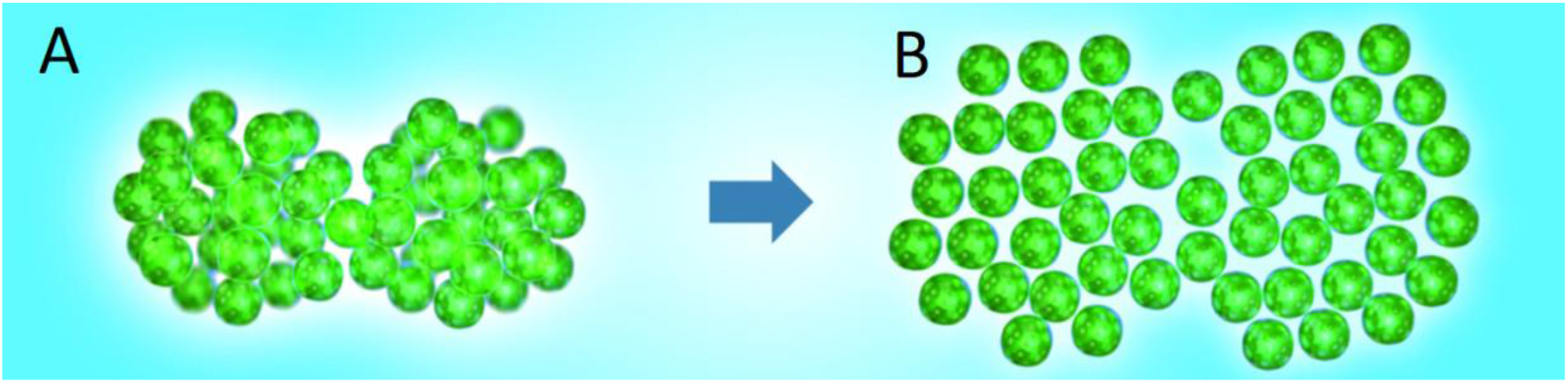
Estimation of colonial cell numbers by flattening the colonies; (D) original colony before desiccation, (E) flattened colony.

### Cell count estimations

We compared the accuracy of three cell count estimations:

1. traditional cell count estimation carried out by experts (A1)
2. geometric approach, based on sphere packing and cell size and cell distance measurements (A2)
3. regression approach based on the colony volume and cell number relationships (A3)

#### Traditional approach (A1)

During traditional microscopic analysis of phytoplankton samples, the analysers have to give an estimation on the number of cells in the colonies appearing in the field of the microscope. We asked 15 experts to give an estimation for the cell numbers in the case of 100 colonies (Fig.2). We have shown at least three photographs of each colony taken in different layers to the experts who had 40 seconds for the visual investigations. Estimated cell counts were registered in paper, and these were tabulated electronically. The ratio between experts’ estimations and reference cell numbers were used to illustrate estimation bias of each expert. Results of the experts are illustrated as boxplots.

**Fig. 2.**
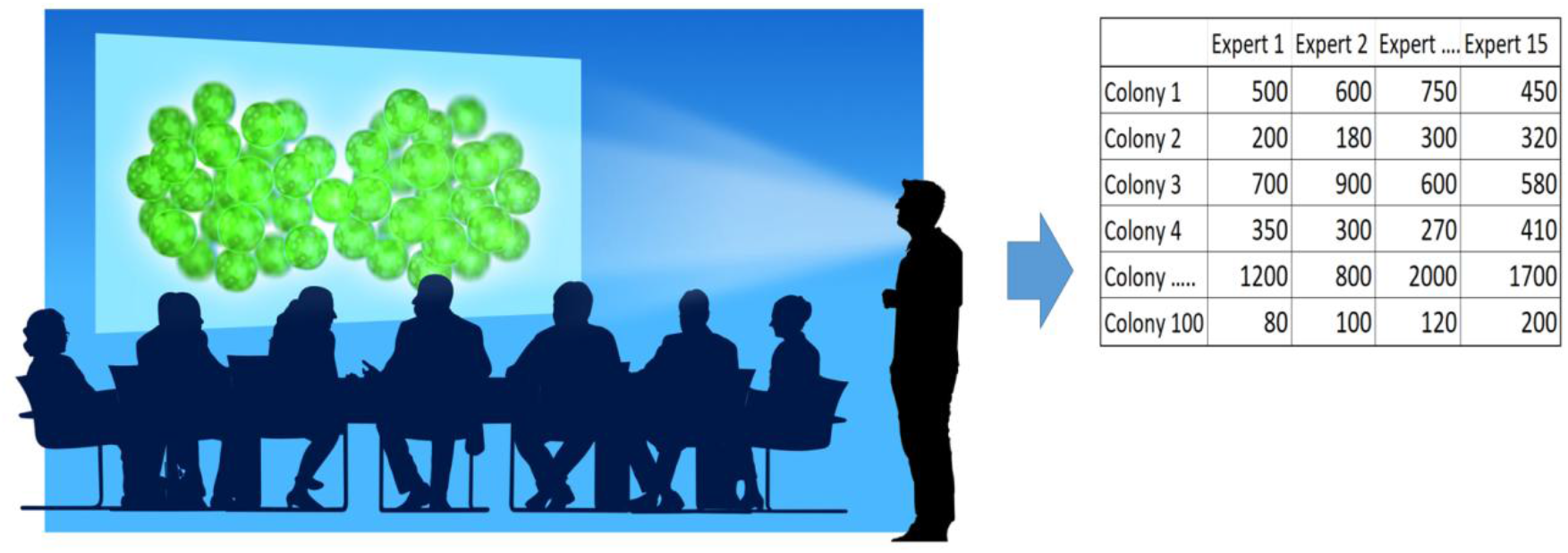
Visual estimation of colonial cell numbers. Data are tabulated and computerized.

#### Geometric approach (A2_3D_ and A2_ell_)

In our geometric approach, we considered colonial cyanobacteria as distantly packed spheres in a 3-dimensional space (Fig. 3). Arrangement of non-overlapping spheres in a Euclidean space and the calculation of packing density is an age-old problem in geometry (Kepler, 1611). Several models have been developed to estimate packing density of differently arranged spheres and results range between 0.5236 (cubic arrangement) to 0.7405 (π/3×√2, i.e. Kepler’s conjecture). Although cyanobacterial cells are small and do not, or rarely touch each other, theoretically they can be enlarged to the size at which they densely fill the available space. Since packing density of loosely packed spheres is approximately 0.55 (Zamponi, 2008), overall cell volume of the enlarged cells will be the 55% of the colony volume. Volume of the enlarged cells can also be calculated, because their radius equals with the half of the distance between cyanobacterial cells (supposing that distances of cell centres have been measured; Fig. 3). The colonial volume/enlarged cell volume ratio gives the cell numbers, and thus using the volume of the original cyanobacterial cells their overall volume can be calculated.

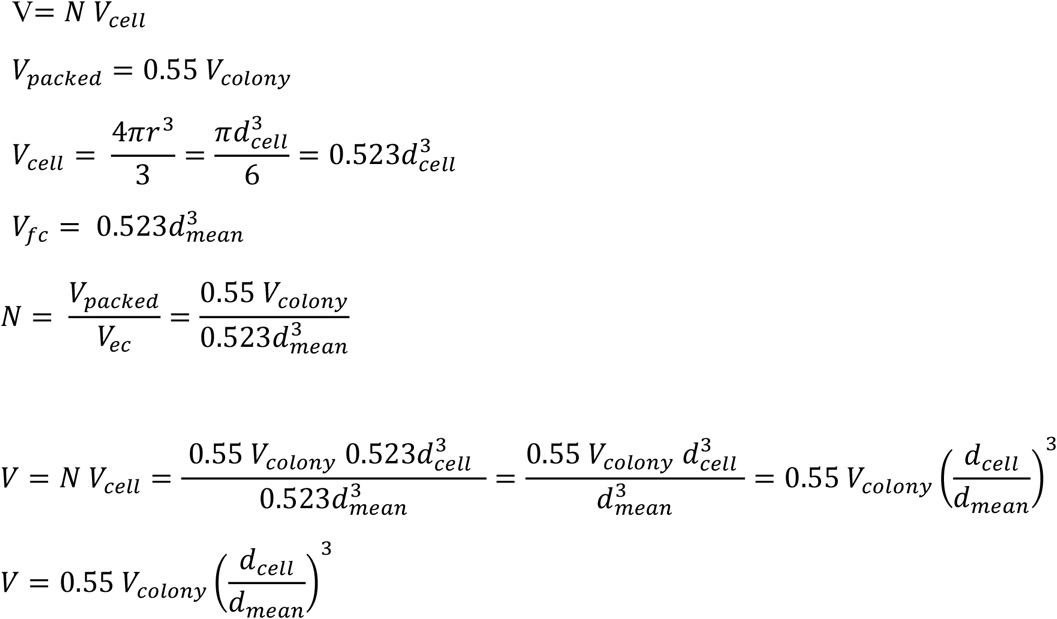

V: overall volume of the cells in the colony
V_cell_: median volume of the cells in the colony
N: number of cells in the colony
V_packed_: volume of the colony filled with loosely packed enlarged spherical cells
V_colony_: overall volume of the colony (cells and mucilage)
V_ec_: mean volume of a virtually enlarged cell
d_cell_: mean diameter of the cyanobacterial cells in the colony
d_mean_: mean diameter of the fattened cells, which is equal with the mean distances measured between the cells’ centers.
r: radius of the sphere (here the cells)

**Fig. 3.**
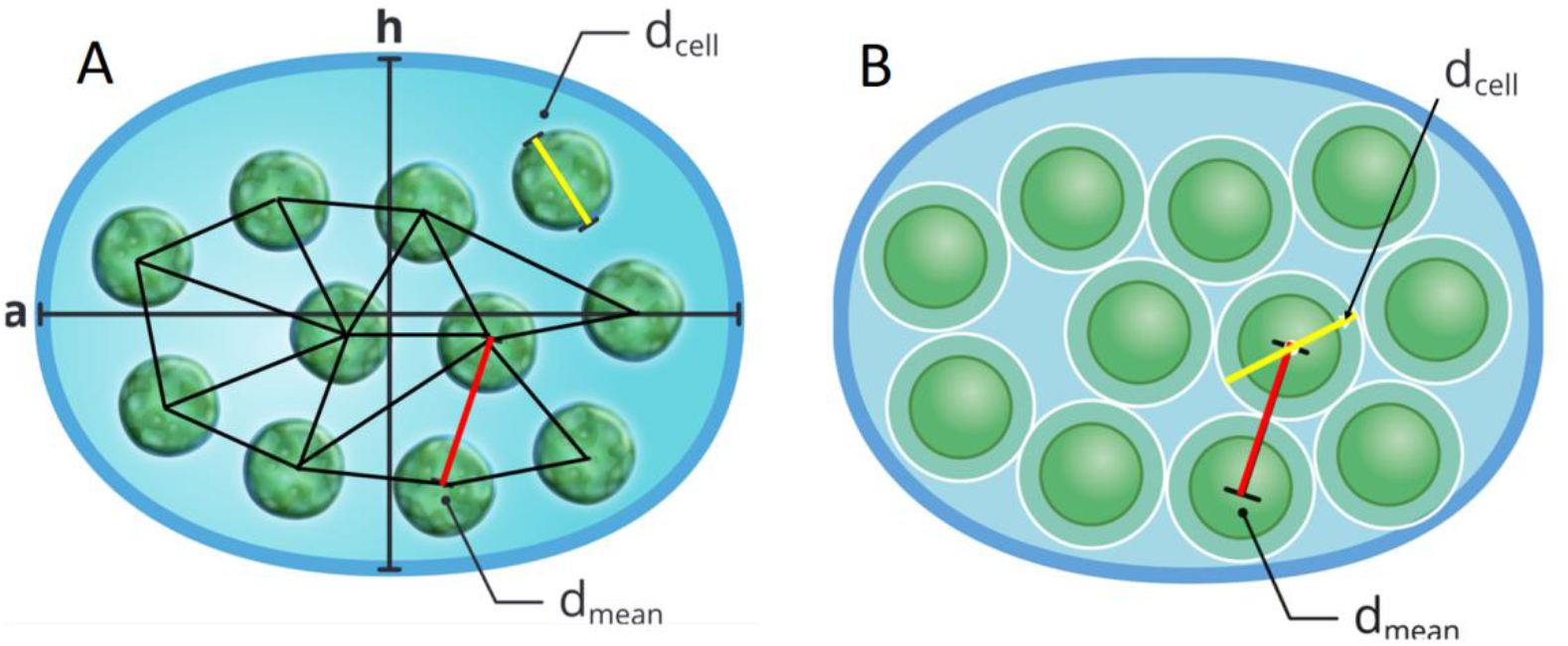
Cellnumber estimation by close sphere packing. A: loosely arranged sells in the colony; black lines are cell distances; a: length, h: width of the colony; dcell: average diameter of a cells, dmean: average distance between the cell centres. B: colony with enlarged cells. Cell diameters are equal with average cell distances measured between the centre of cells.

In this study, two overall colony volume (V_colony_) values were used: A2_3D_ measured on the 3D models and A2_ell_ estimated by the experts using linear measurements on the colonies (Alcántara et al., 2018) (Fig. 4). We made the geometric cell count estimations using the results of both approaches separately and the estimation biases were expressed as estimated/reference ratios.

**Fig. 4.**
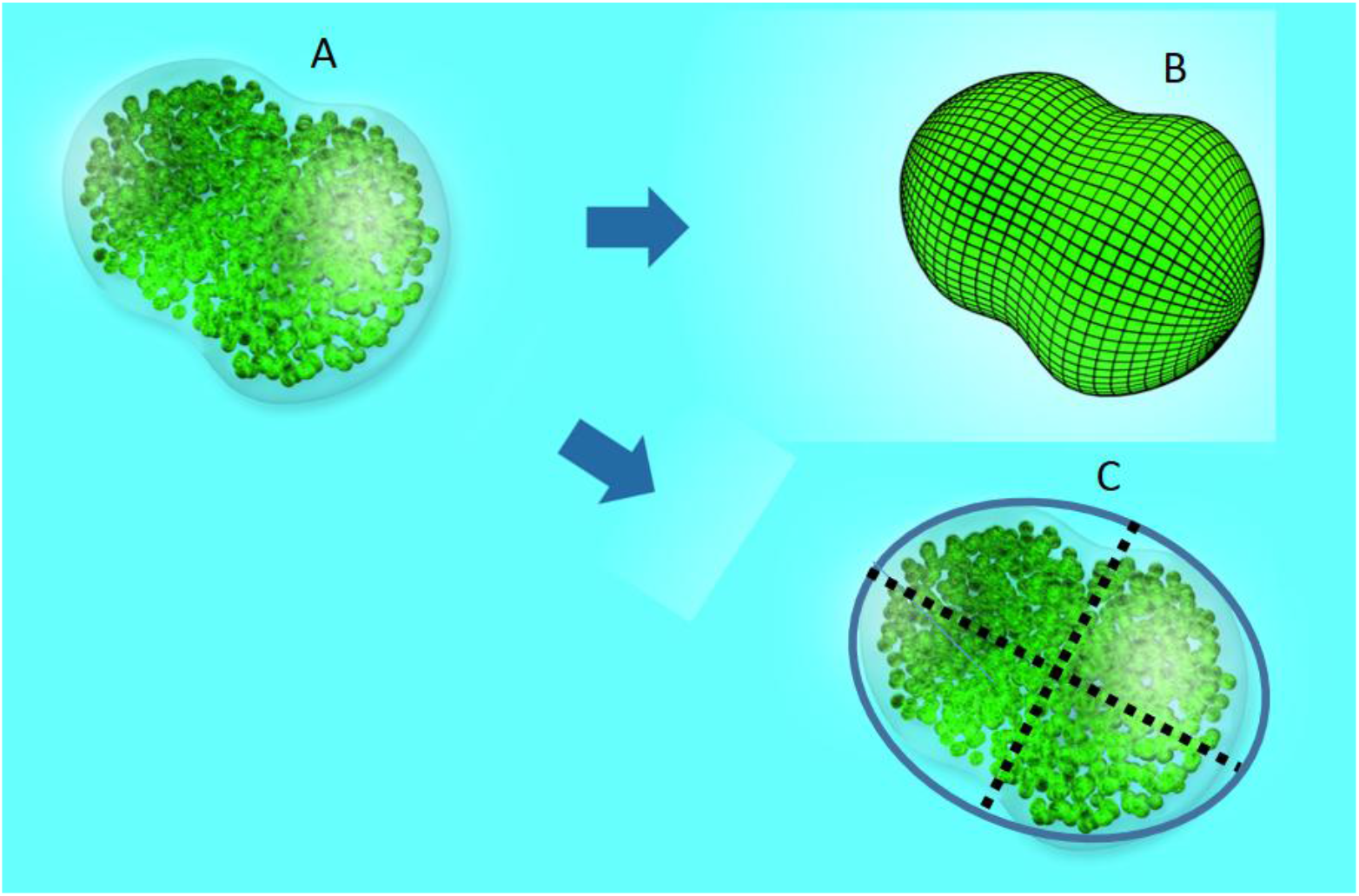
Estimation of the volume of cyanobacterial colony (A), using real like3-D modelling (B), and approximated by ellipsoid (C).

#### Regression approach

To study the accuracy of the regression approach to questions have to be answered: what is the accuracy of colony volume estimations, and what is the accurate relationship between the colony volumes and overall cell biovolumes within the colonies.

To estimate the accuracy of colony volume estimations we created the real-like 3-D models of each (100) studied colony and these served as references (Fig. 4). For the real-like 3-D visualization, we used the free open source 3-D graphics application Blender 2.79 (Blender, 2020). The models were based on the photographs that were taken at different layers on the colonies. Volume of the models were calculated using the NeuroMorf software toolset (Jorstad et al., 2006). Since diameters of the colonies are known from microscopic measurements, colonial volumes provided by NeuroMorf were transformed to *μm^3^*-scale.

We estimated the colony volumes using the formulae proposed by Alcántara et al (2018):

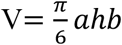

where, V: volume of an ellipsoid, a: length, h: width and b: depth of the colonies. We estimated the depth of colonies by the following formula:

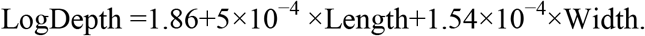

This estimation requires measuring the length and width of the colonies. Using the photographs that were previously used for cell count estimation of the colonies the experts (15) were asked to measure the’ linear dimensions of cells and colonies, moreover the mean distances of cells on 30 colonies (Fig. 4.). Cell diameter measurements were done on randomly selected five cells, while during cell distances measurements, depending on the structure of the colonies, ten to twenty-five cell distances were measured.

The photographs were imported to LibreCad software (an open-source 2-D CAD application) and all measurements where done using the ruler function of the program. Since cell diameters are known from microscopic measurements, cell distance values provided by LibreCad could be transformed to μm-scale.

These estimated colony volumes were compared to that of the reference given by the 3-D approach. Estimation bias was expressed as the ratio between the estimated volumes (Alcántara et al., 2018) and volumes given by 3-D modelling of the colonies.

To study the second question i.e. “what is the relationship between the colony volumes and overall cell biovolumes within the colonies?” the sum of the reference cell biovolumes were plotted against the reference colony volumes, and the relationship between variables was studied using OLS (Ordinary Least Square) analysis.

## Results

We involved 100 cyanobacterium colonies into the analyses, which based on their morphology belonged to the *Microcystis* and *Chroococcus* genera. Real-like 3-D modelling, cell countings, cell size and distance measurements were carried out in the case of each colony as described in the Methods section. Cell distance, cell and colony size measurements were performed on a smaller (30) set of colonies by experts.

In the first step we compared the results of the traditional approach (A1) estimating the cell counts by experts; and the geometric (sphere packing) approaches (A2) estimating the cell numbers by using their average cell sizes and distances in colonies of which volumes were determined by 3-D modelling (A2_3D_); and by treating them as ellipsoids (A2_ell_) (Alcántara et al., 2018). We created ratios of these values and the reference ones, i.e. those given by direct counting after colony disintegrations. Multiplying these values by 100 we get percentages. Subtracting 100 from these values expresses the results as percentage differences. We used the number of estimations in the percentage difference categories as basis for comparison (Fig.5 A-C).

**Fig. 5.**
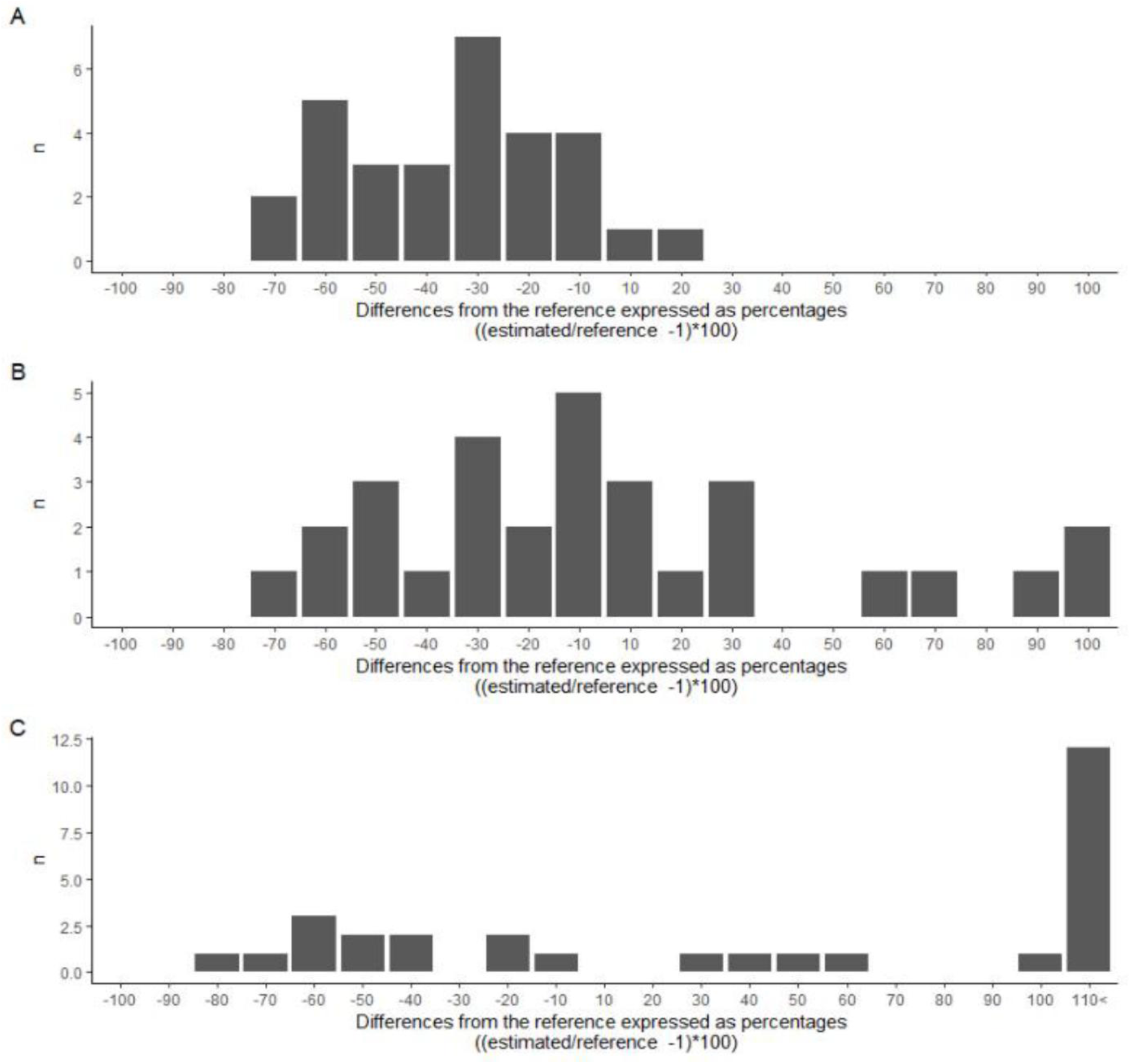
Distribution of estimation bias for the three (A1, A2_3D_, A2_ell_) cell count estimation approaches; a: visual estimation of cell count by experts (A1); b: close sphere packing using colony volumes derived from 3-D models of colonies (A2_3D_); c: close sphere packing using colony volumes derived from the ellipsoid approximation approach (A2_ell_). Values on the x axis are differences from the reference expressed as percentages ((estimated/reference - 1)×100). The columns indicate the number of observations in the given range.

The above mentioned approaches (A1, A2_3D_, A2_ell_) showed considerable differences both in terms of data dispersion and location of the mode. The first approach (Fig. 5A) was characterised by low data dispersion (−70 to 20%). Location of the mode indicated −30% estimation error. In the case of the A2_3D_ and A2_ell_ approaches, which are based on the overall colony volume estimation and cell distances, dispersion of data was larger. When colony volumes were estimated by using 3-D models (A2_3D_) (Fig. 5B) data dispersed from −70 to 100%. The mode however located at −10 %. The A2ell approach (when colonies were considered as ellipsoids) provided the less accurate predictions (Fig. 5C). Estimated volumes dispersed from −80 to >120% percentage range and the mode fell outside the >110% value.

To study the differences among experts who performed the counting, estimated values were divided by the reference values and these were presented as boxplots (Fig. 6). Results revealed large individual differences in cell number estimations. Median values of estimated cell counts fell in the range of 0.5-1.5 in the case of 75% of experts. However, most experts underestimated the real cell numbers, and only three experts gave overestimations. Colonial cell numbers had a pronounced impact on the estimation results. In the case of colonies in which the number of cells was higher than approximately 300, overestimations only occasionally occurred (Fig. 7).

**Fig. 6.**
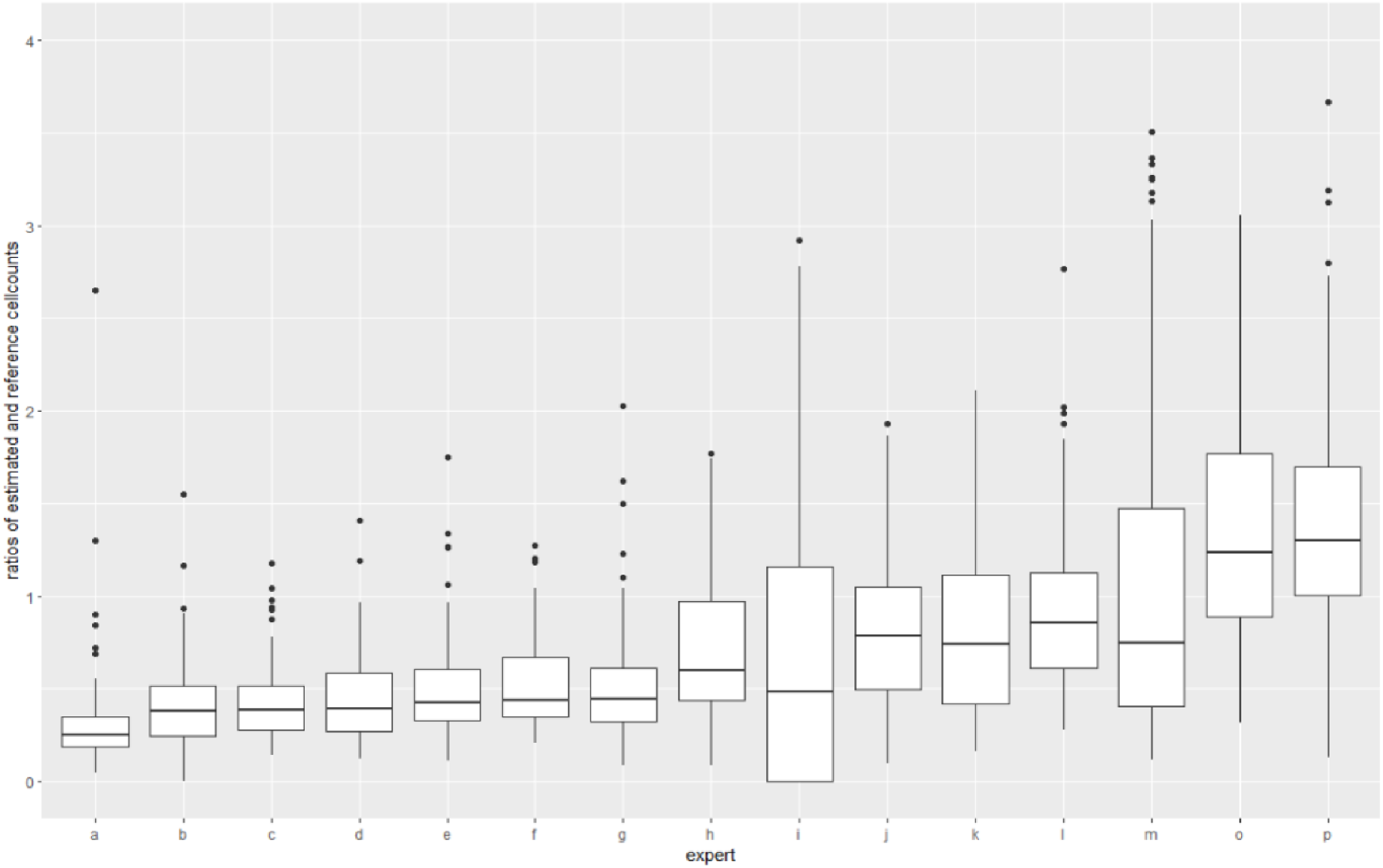
Distribution of estimated cell counts provided by the 15 experts. Values on the y axis are the ratios of estimated and reference cell counts.

**Fig. 7.**
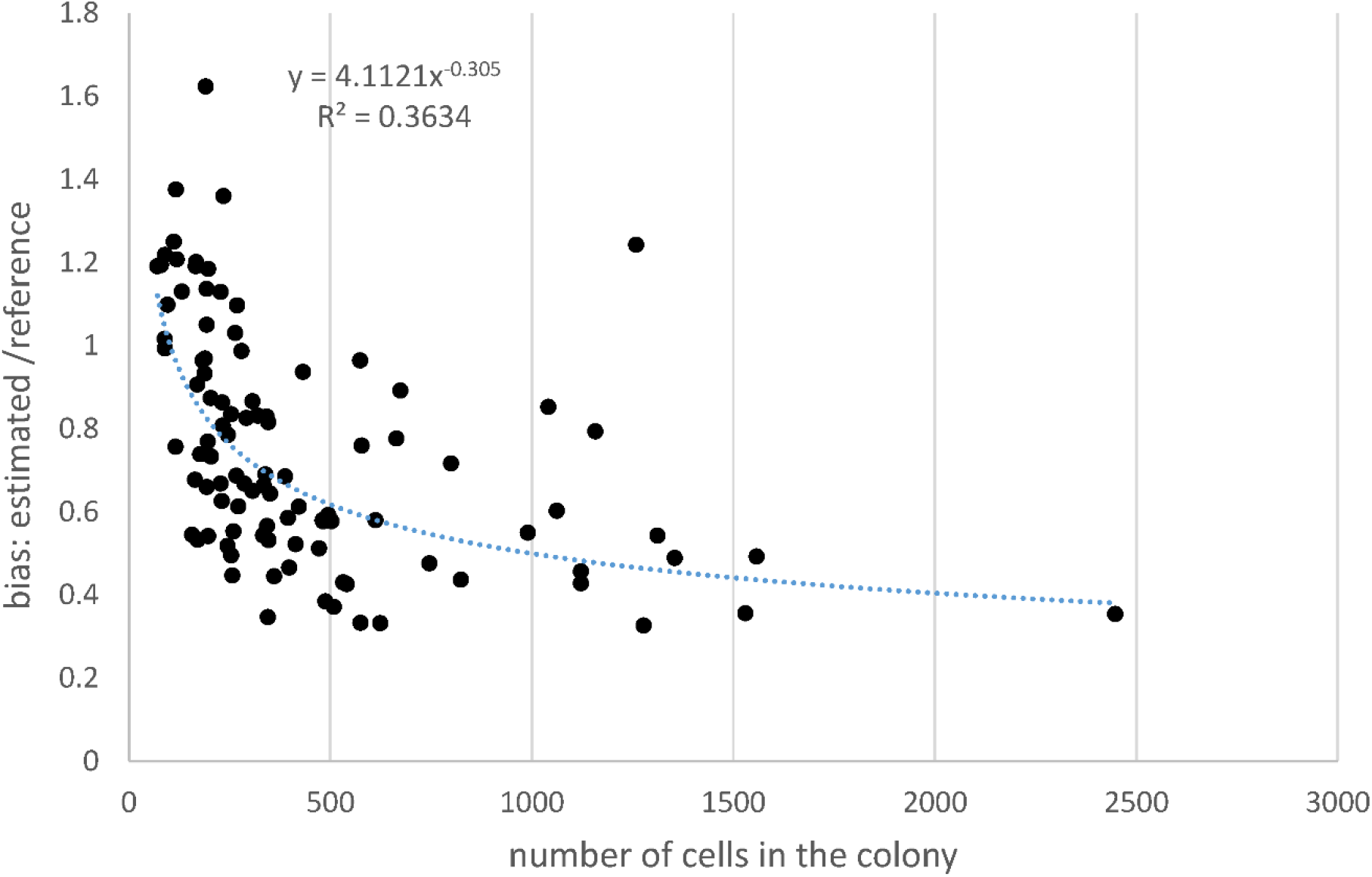
Relationship between the number of cells in the colony and the estimation bias expressed as: estimated /reference. Cell numbers have been notoriously underestimated when colonial cell numbers exceeded 500.

As was shown in the methods, the regression approach (A3) is based on the overall colony volume/cell number relationships. This approach requires accurate colony volume estimation and strong relationship between colony size and cell numbers. Our results revealed that considering cyanobacterial colonies as ellipsoids leads to severely biased estimates for their volumes (Fig. 8). Volumes of the ellipsoids were calculated on the basis of measured length and width of the colonies and the calculated depth values. Means of the experts’ volume calculations were divided with that of the reference (i.e. calculated by the Neuromorph on the 3D models). We plotted these values against the volume of the colonies (Fig. 8). The results showed large uncertainties in colony biovolume estimates. In the majority of cases considering the colonies as ellipsoids resulted in overestimation of the colony volumes. These uncertainties however did not depend on the volumes. Large overestimations occurred along the whole scale of colony volumes.

**Fig. 8.**
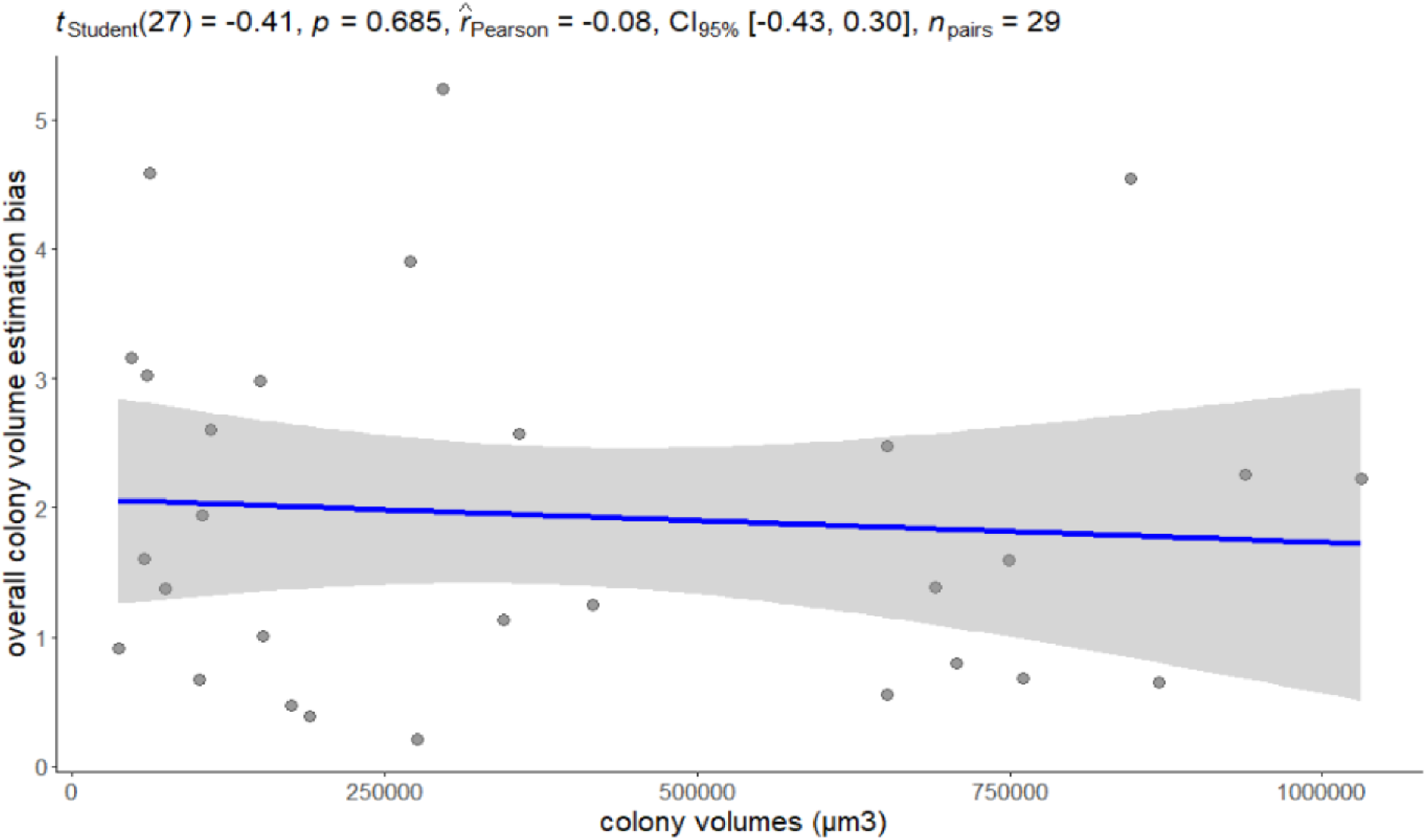
Relationship between the overall colony volume estimation bias and colony volumes β (μm^3^). Estimation bias is given as the ratio between estimated and reference colony volumes. Estimated volumes were calculated using the ellipsoid approximation approach (Alcántara, et al., 2018), while volumes based on computer calculations on real like 3-D images of colonies, were considered as reference.

We found strong linear relationship between colony volumes (based on the 3D models) and cell volumes (using reference cell counts of disintegrated colonies) (Fig.9). The relationship was characterised by high R^2^ value (R^2^ =0.54) and even scatter of residuals around the regression line (homoscedastic). However, the residuals cover a range of approx. one order of magnitude, which indicates low precision.

**Fig. 9.**
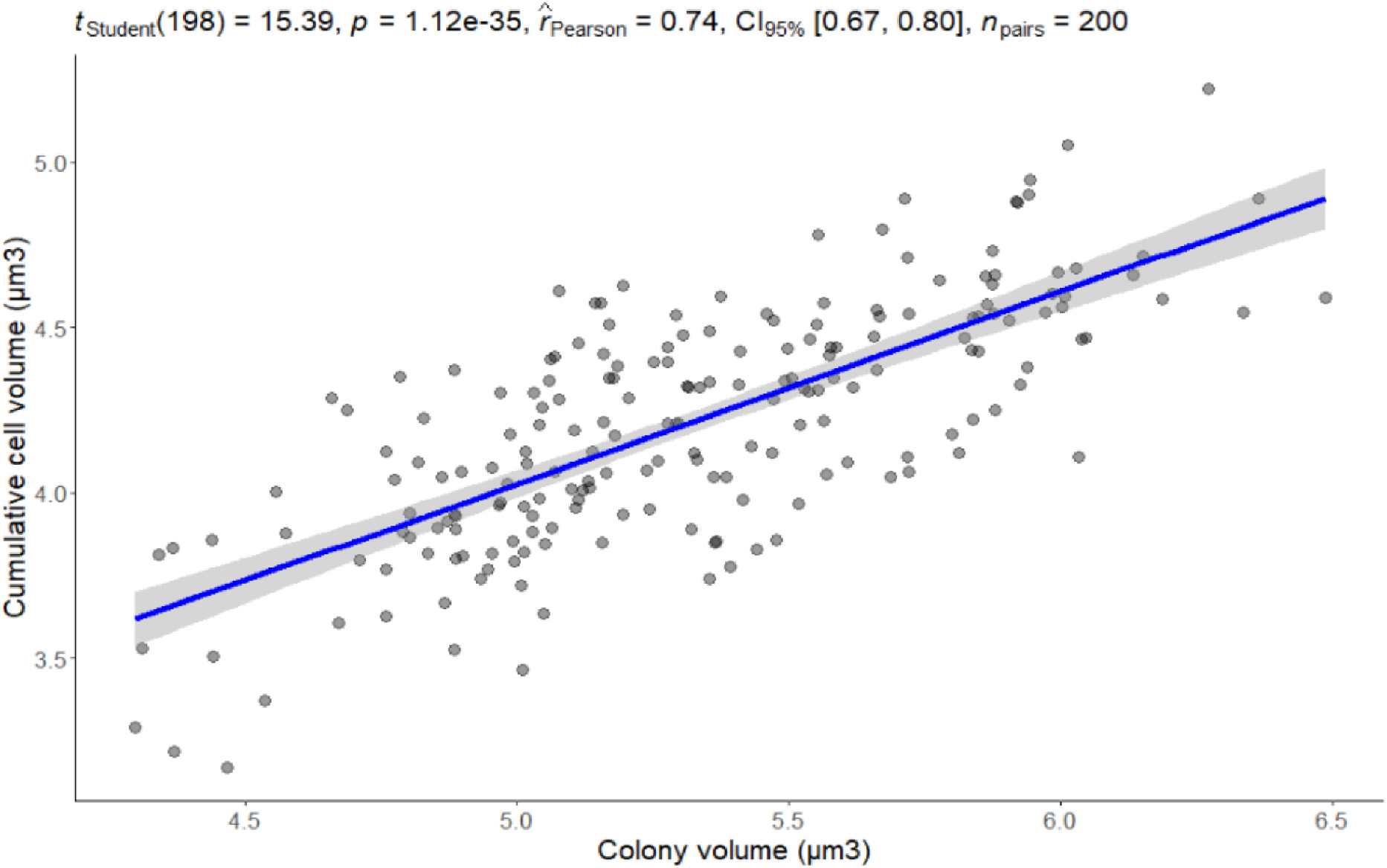
Colony volume (μm^3^) and cumulative cell volume (μm^3^) relationship for cyanobacterial colonies. Colony volumes were determined using 3D images of colonies, while cell numbers were determined after disintegration of colonies. Cells were considered as spheres. Cell diameters were determined during microscopic measurements.

## Discussion

Cyanobacteria are the most notorious bloom forming planktic photosynthesizers in freshwaters their biovolumes are frequently used bioindicators in lake quality assessment (Carvalho et al., 2013). While volume of single celled and filamentous forms can be measured accurately, in the case of colonial cyanobacteria the estimation of cell numbers has unknown bias, moreover it is also questionable whether ignoring the volume of mucilage is acceptable or not.

In the case of many cyanobacterial species, the gelatinous envelope covering the living cell, filament and colony has a prominent appearance in both microscopic and macroscopic examinations (De Philippis et al., 2001). Of the many names used, exopolysaccharide (EPS) is perhaps the most appropriate one for the gelatinous forms (Chen et al., 2016). In addition to being an important taxonomic marker for many species, it has many physiological, ecophysiological, and biogeochemical functions. Without wishing to be exhaustive, the protective function, metal binding capacity, its effect on the nutrient-cycles, and the tolerance of the organism to frost and temperature should be emphasized. As well as in many cases it serves as a substrate and habitat for microorganisms (heterotrophs and autotrophs too) and it has effects on the organization of the own cells of the cyanobacterial species (colony forming) (Liu et al., 2018; De Philippis & Vincenzini, 1998).

Considering the amount and significance of EPS, it seems evident to calculate with this product when calculating the biovolume of cyanobacterial species (Liu et al., 2018). However, the heterogeneity, visibility, unique species-, and environment-specific appearances of EPS complicate the issue. Overall, based on the EPS chemistry, it is a hygroscopic acidic heteropolysaccharide natural polymer, that also contains proteins (not to mention many other extracellular organic compounds produced by Cyanobacteria) (Chen et al., 2016). In addition, it can be classified into soluble and insoluble fractions, making it difficult to define exactly how to calculate with this material.

Overall, it is difficult to make a decision whether to consider the cyanobacterial EPS content in studies or not. Although in water quality monitoring the applied metrics calculate exclusively with the cell volumes, there can be several questions (concerning nutrient cycling, grazing, sinking properties) in the case of which investigation of EPS content cannot be set aside.

The study of colony volume / cell volume relationships gave us controversial results. Although the relationship was strong, the large amount of scatter in data implies that this approach is also prone to considerable error. The large scatter partly can be explained by that we involved different colonial species into the analyses. However, colony compactness (sum of cell volumes/overall colony volume ratio) may change even within a species, depending on the actual constraints of the environment (Reynolds & Jaworski, 1978; Wu et al., 2020). The experienced one order of magnitude differences in the residuals enables only rough estimation of cyanobacterial cell biovolumes.

The multi-layer photographs taken on the colonies and the subsequent 3D modelling of them revealed high diversity in colony shapes. Spherical, ellipsoid, tube-like or irregular reticulate forms also occurred. Therefore, not surprisingly the approach according to which the colonies should be treated as ellipsoids (Alácantra et al., 2018) resulted in considerable bias in volume estimation. Colony shape is a species-specific property (Komárek & Komárková, 2002), but because of the high phenotypic plasticity of colonial cyanobacteria, size and morphology of colonies can change responding to various biotic and abiotic factors (Xiao et al., 2018). Infochemicals released from some grazing flagellates induce *Microcystis* production of mucilage and colony formation (Burkert et al., 2001; Yang & Kong, 2012). Among abiotic constraints low temperature and shortage of light (Li et al., 2013; Xu et al., 2016), high concentration of lead (Bi et al., 2013) or calcium (Wang et al., 2011) also induce the formation of mucilaginous colonies.

The proposed geometric approach, which is based on close sphere packing (A2), requires exact colony volume estimates. However, this can be done only by using the tedious 3-D modelling of the colonies. Other colony volume estimations have considerable bias causing large mistakes in cell count estimations, despite cell size and cell distance measurements are performed accurately.

Comparing to the geometric (sphere packing) approach (A2_3D_, A2_ell_), mean values of experts’ cell number estimations (A1) showed slightly lower accuracy but higher precision. These surprisingly good results can be accounted for by an innate ability of human mind and learned skills. We humans and some other species are capable of subitizing (Kaufman & Lord, 1949), i.e. to recognize a small group of objects without counting. This evolutionarily primary numerical ability is a fundamental skill in the development of number sense. Subitizing, however has got its limitations, i.e. it cannot perform well on multiple sets, or on high number of items (Liu et al., 2020). Cell number estimation however is analogous to the problem of crowd counting, which is an active research topic in computer vision (Ranjan et al., 2018). The key idea behind crowd counting is simple: area times density. Enumeration of the items in regular crowds can be done by counting them in selected rows and columns and multiple the values to obtain the final count. This approach is self-evident in the case of regular colonies characteristic for *Merismopedia* or *Eucapsis* genera. In the case of irregular forms, however, this approach cannot be applied. In this study, the experts used the same strategy: they focused on one section of the colony, tried to count it and extrapolated to other sections. This approach was especially successful in the case of colonies with uniform density.

Although in terms of mean values the traditional cell count estimations of the experts provided good results, considerable differences were observed at individual level. The fact that 80% of the observers notoriously underestimated the number of cells in the colonies can be accounted for by the difficulty to make estimation for the spatial extension of a three dimensional object from two-dimensional images provided by the microscope. Some of the experts provided good results both in terms of accuracy (position of the median) and precision (data dispersion) and this indicates that cell count estimation is a learnable skill.

We investigated the accuracy of various cyanobacterial biovolume and cell count estimation approaches. We demonstrated that overall colony volume estimation from linear dimensions have considerable bias. Although the colony volume - cell number relationship was significant because of the large residual variability for any given value on the x axis the regression model provides only a rough estimate for cell counts. The proposed geometric model (based on close sphere packing; A2) can be applied only in those cases when overall colony volume can be safely estimated (A2_3D_). Cell counts were underestimated by the involved experts (A1), but precision (variance) of these estimations was low, and individual differences indicates that estimation competence can be developed.

Although all cyanobacterium biovolume estimation approaches aim to estimate total cell biovolumes, the volume of mucilage in which cells are embedded can also be important.

## Acknowledgement

We would like to thank you the contributing experts.

## Supplementary information

Supplementary Fig. S1. Photographs of original colonies, flattened colonies, the real-like 3-dimensional models and cellnumbers of each measured colony.

## Author contributions

GB conceived the original idea, ETK and GB wrote the manuscript with input from, GV, VBB, JG, ÁL, ZsK. ETK carried out the experiments, VL performed the calculations, IT, TK carried out visualisation, VG performed the computations. All authors discussed the results and contributed to the final manuscript.

