## Supplementary figures and images for "Uncertainties of cell number estimation in cyanobacterial colonies and the potential use of sphere packing"

### Supplementary Figure S1

| Number of colony | Original colony | Flattaned colony | 3D colony | Number of cell |
|---|---|---|---|---|
| 1. | 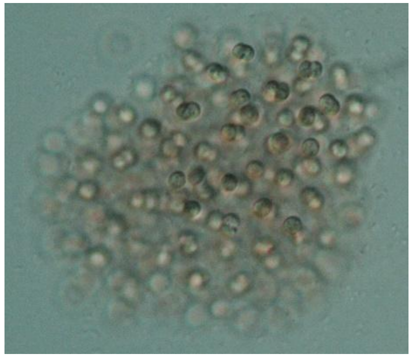 | 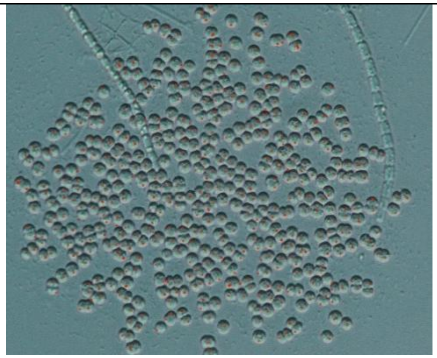 | 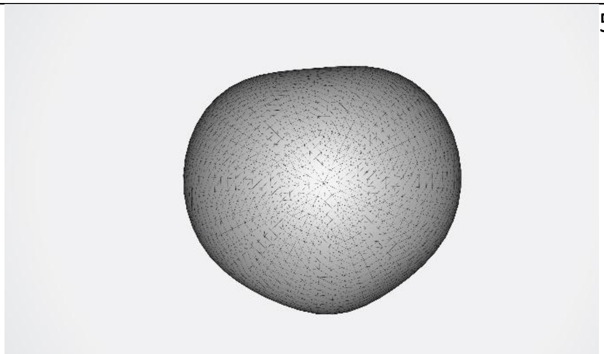 | 509 |
| 2. | 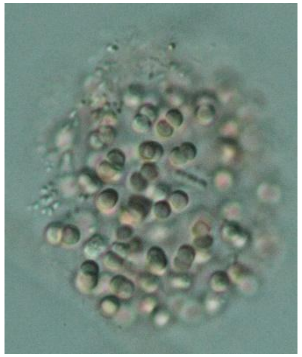 | 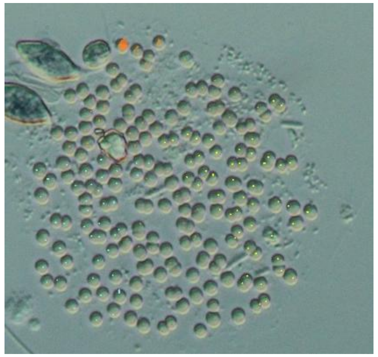 | 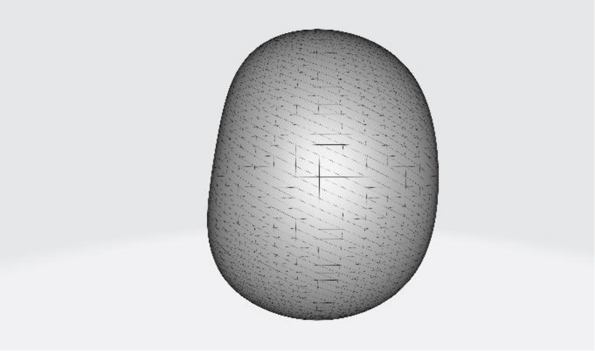 | 197 |
| 3. | 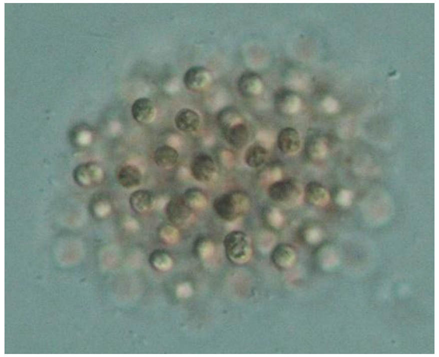 | 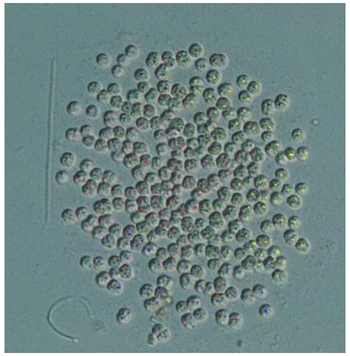 | 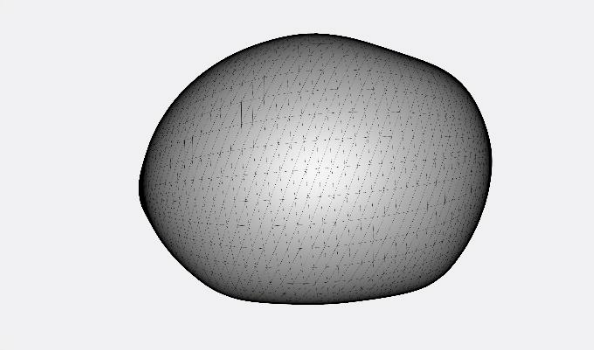 | 272 |

4.

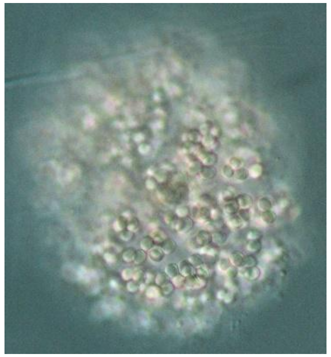

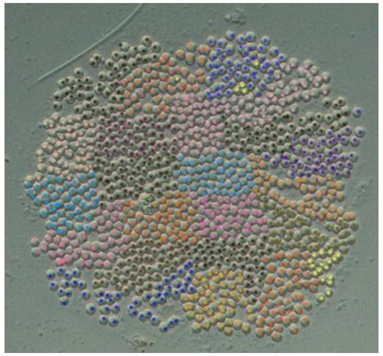

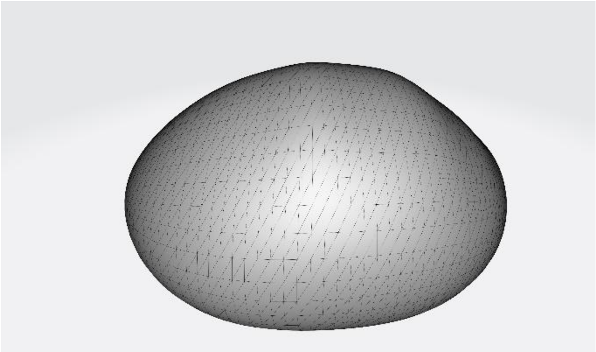

1122

5.

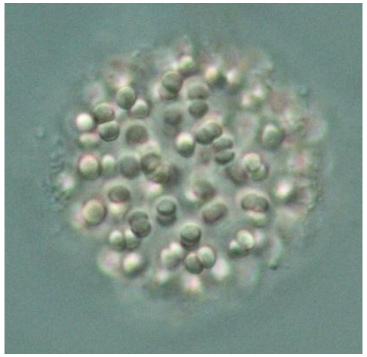

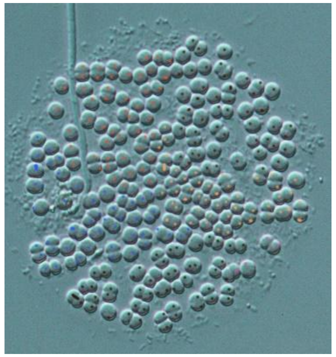

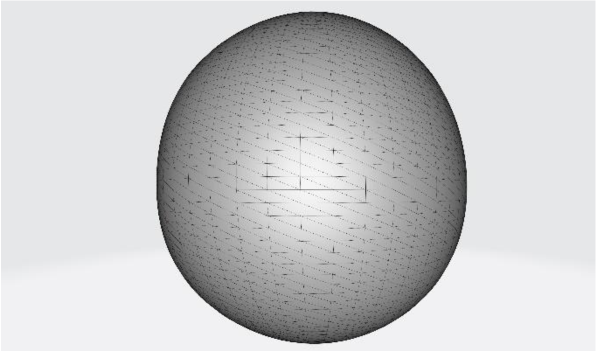

254

6.

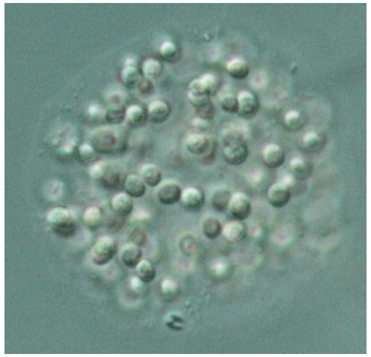

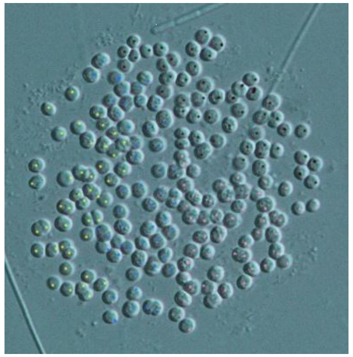

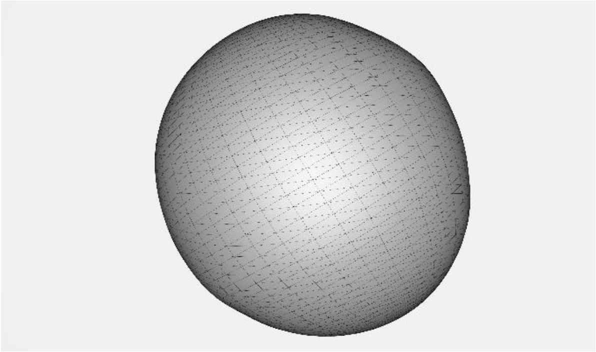

193

7.

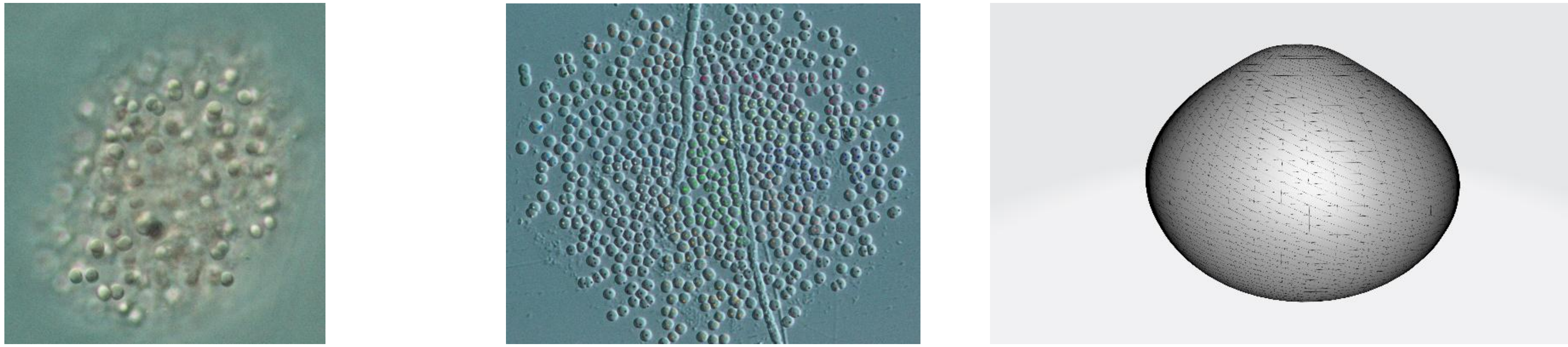

624

8.

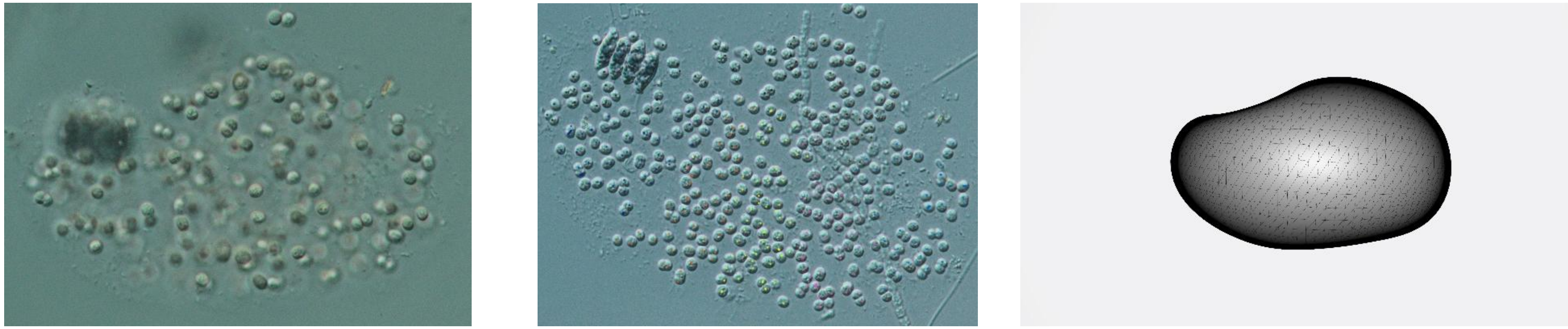

346

9.

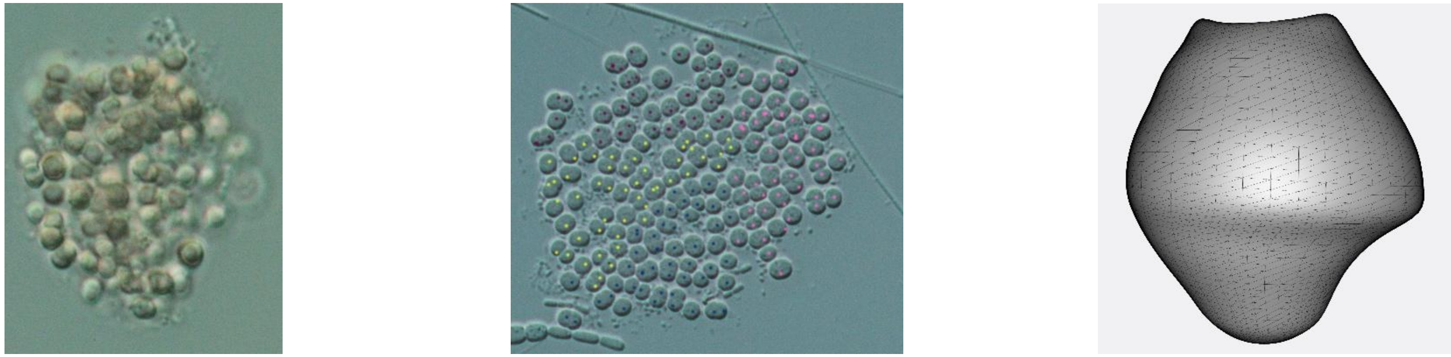

188

10.

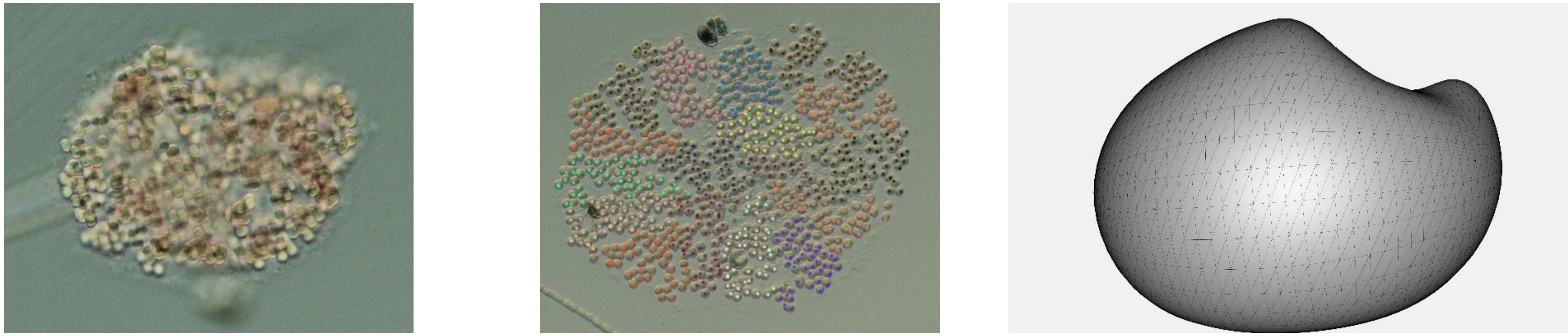

824

11.

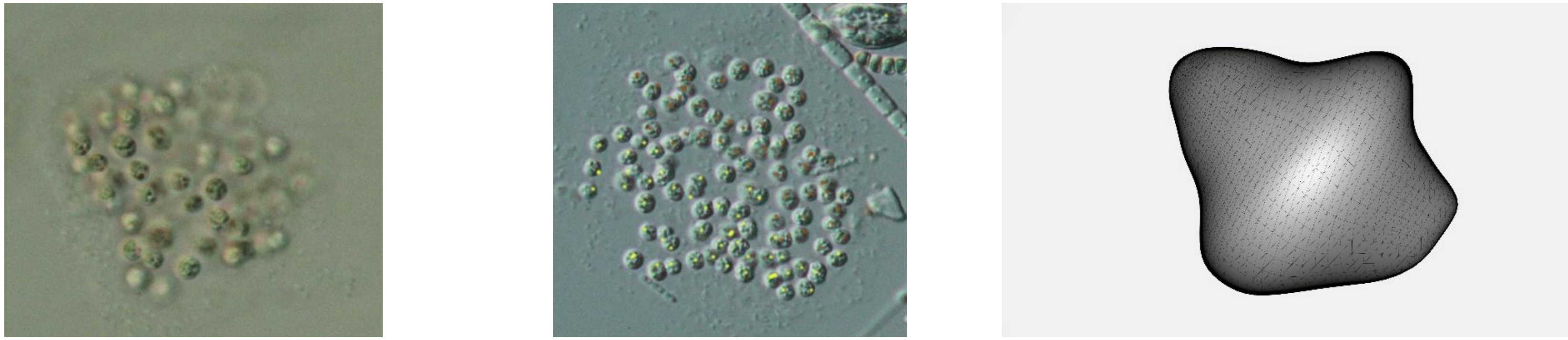

96

12.

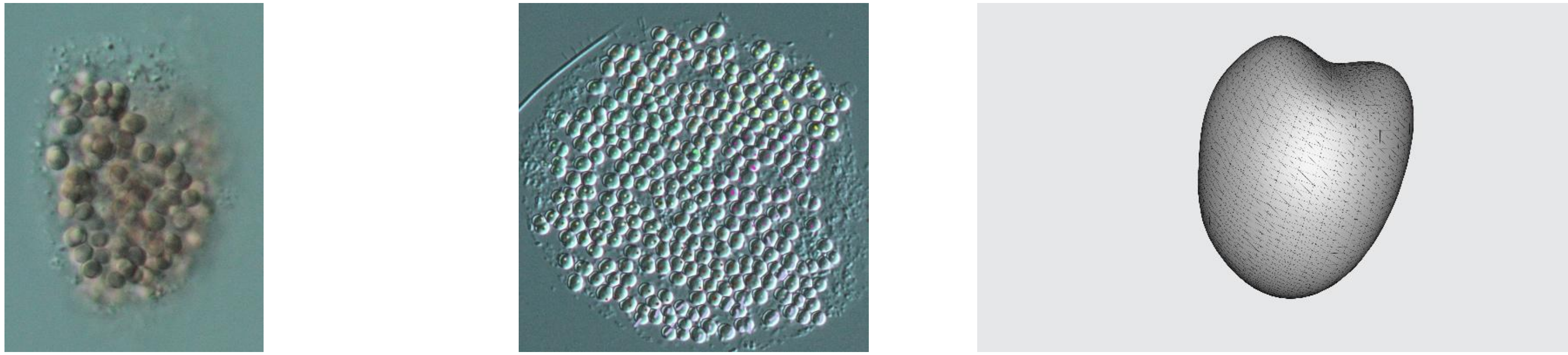

345

13.

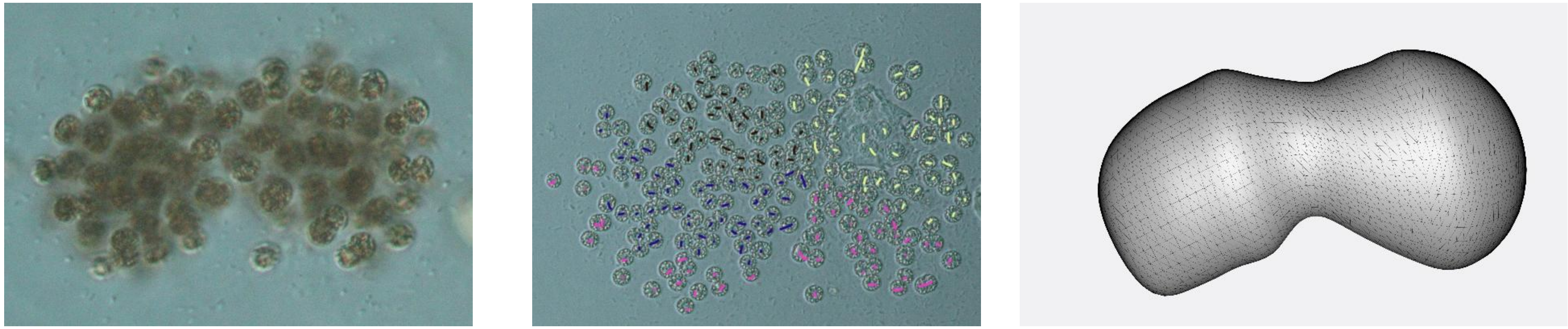

204

14.

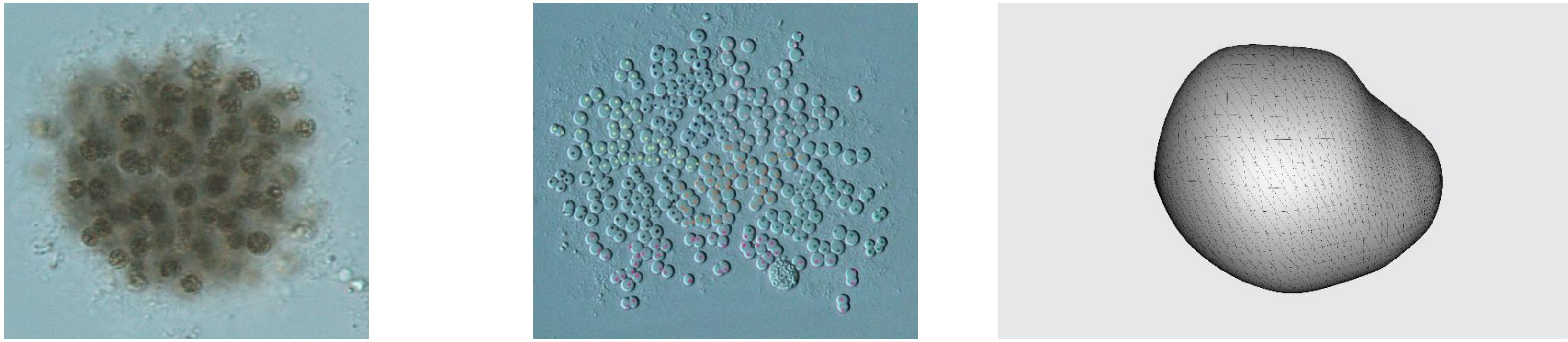

347

15.

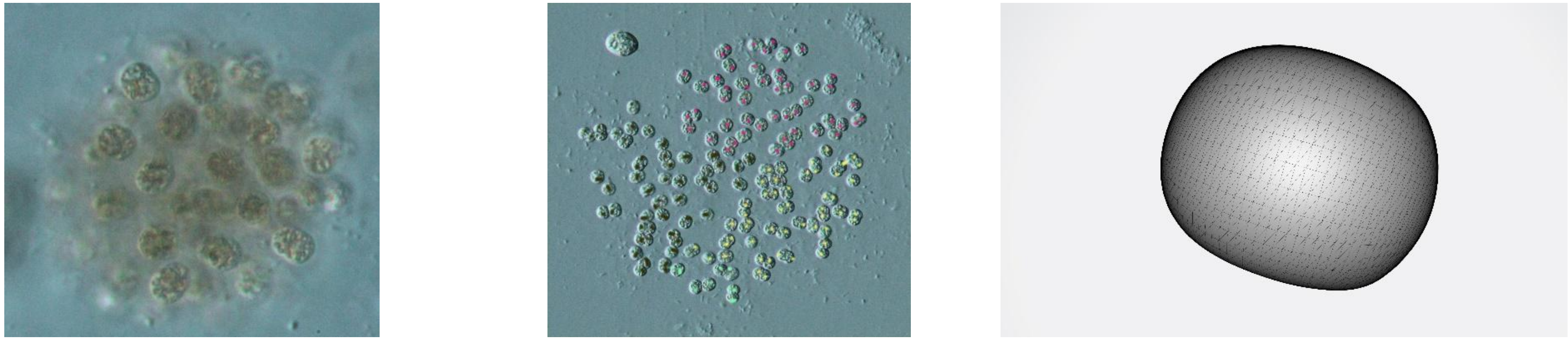

157

16.

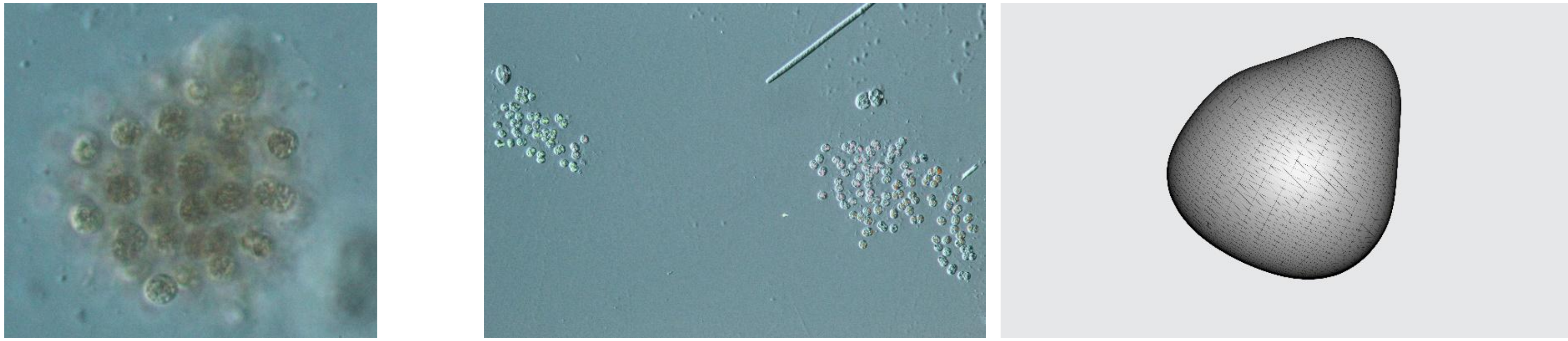

171

17.

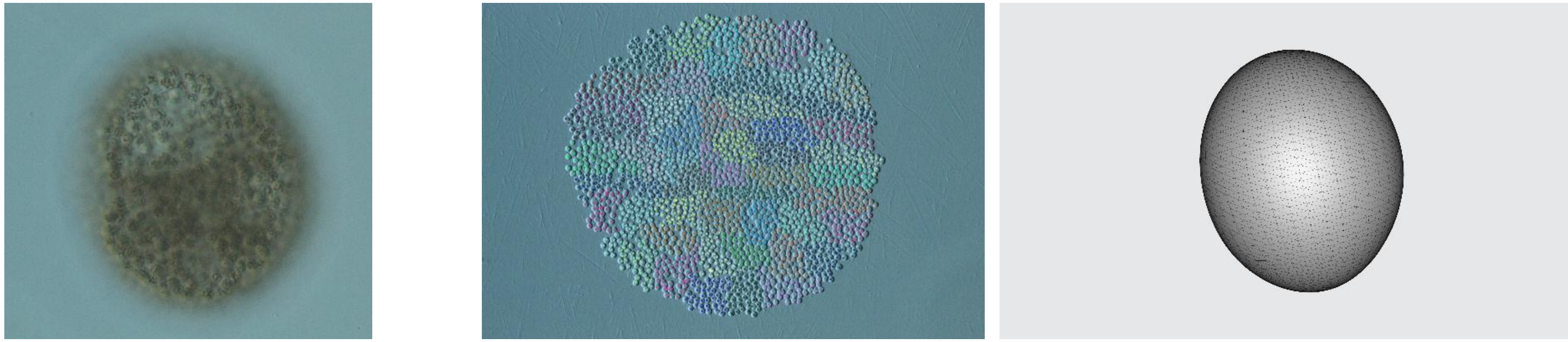

2448

18.

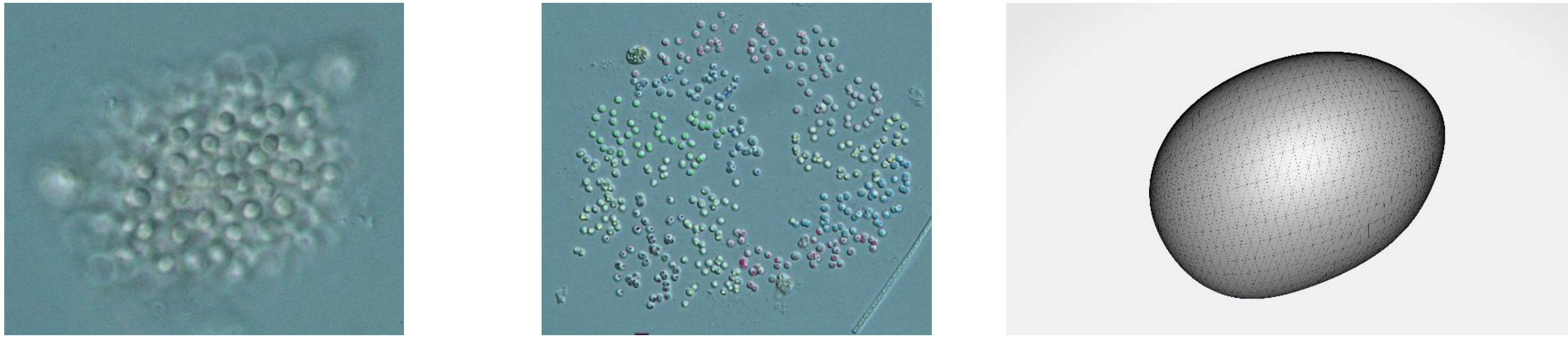

488

19.

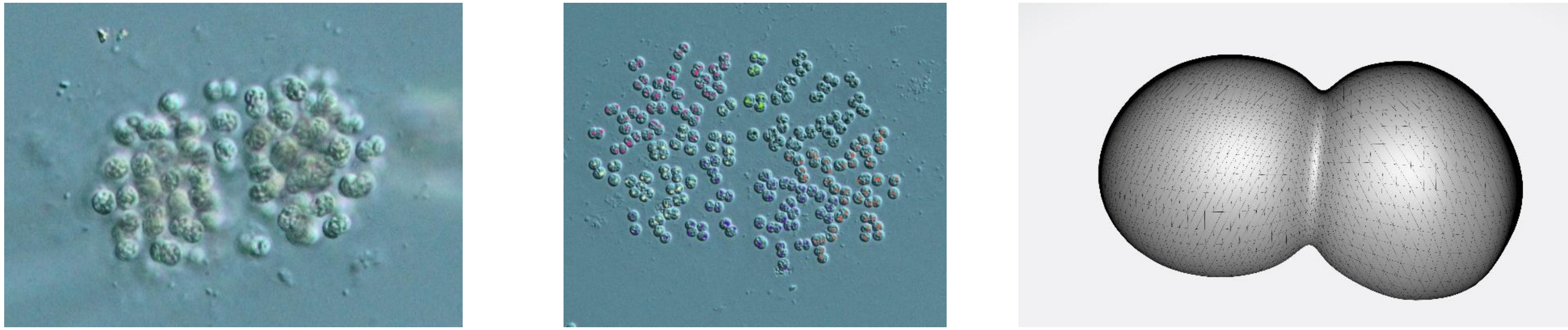

257

20.

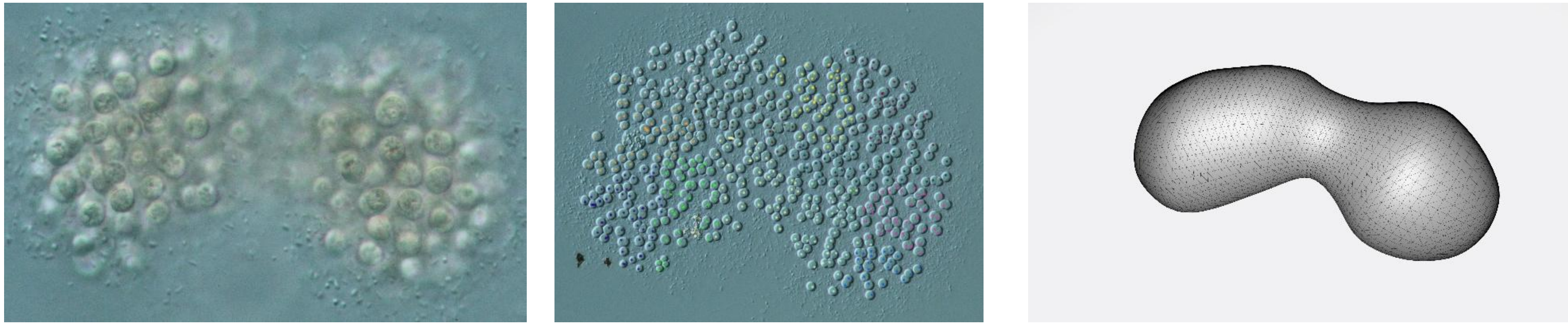

575

21.

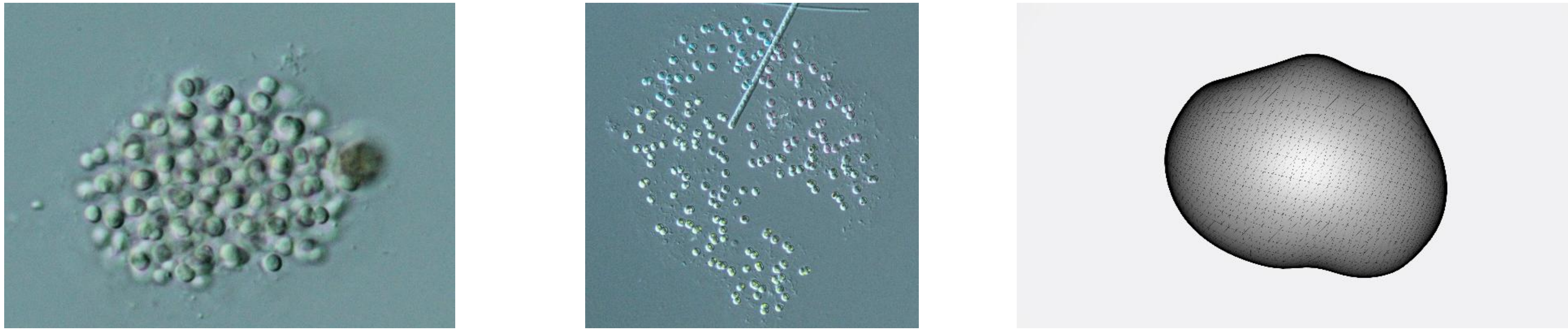

254

22.

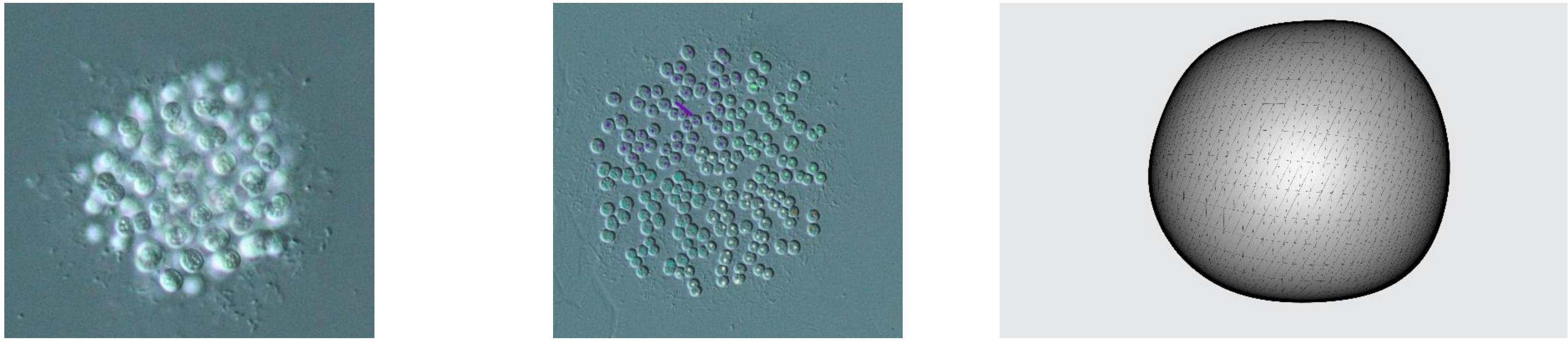

203

23.

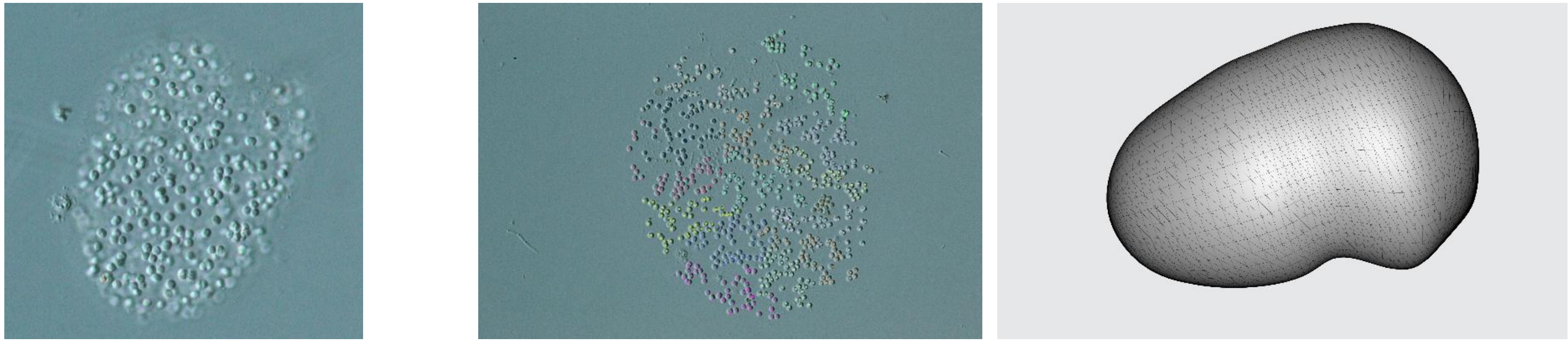

664

24.

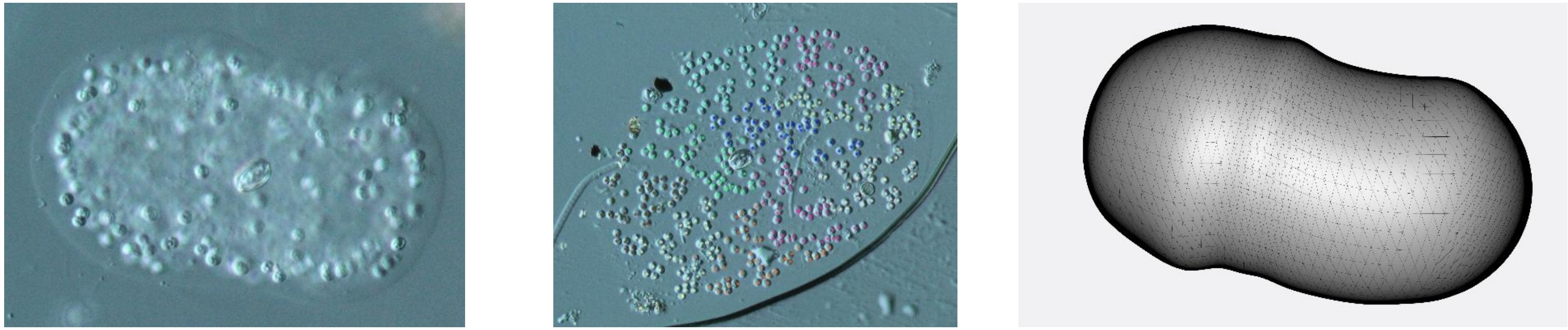

503

25.

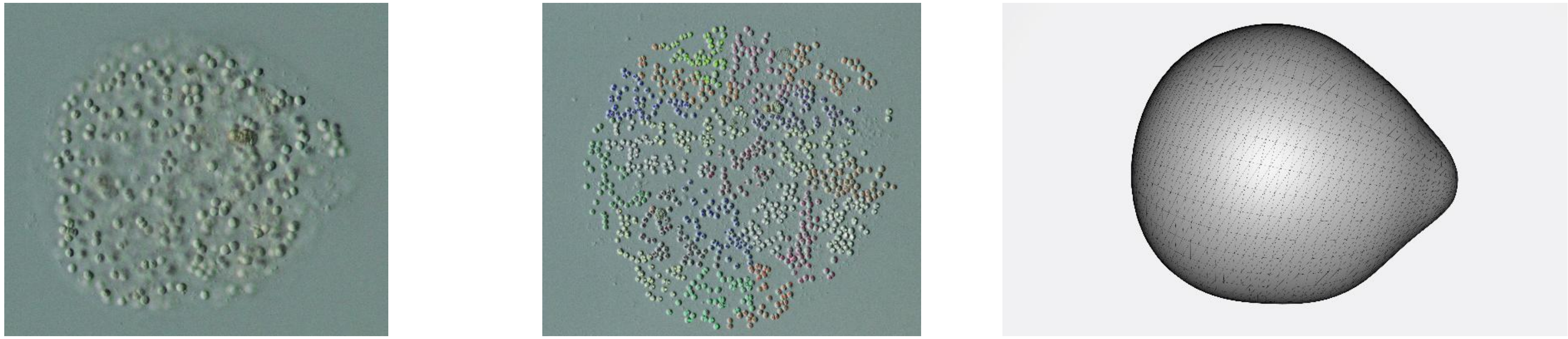

990

26.

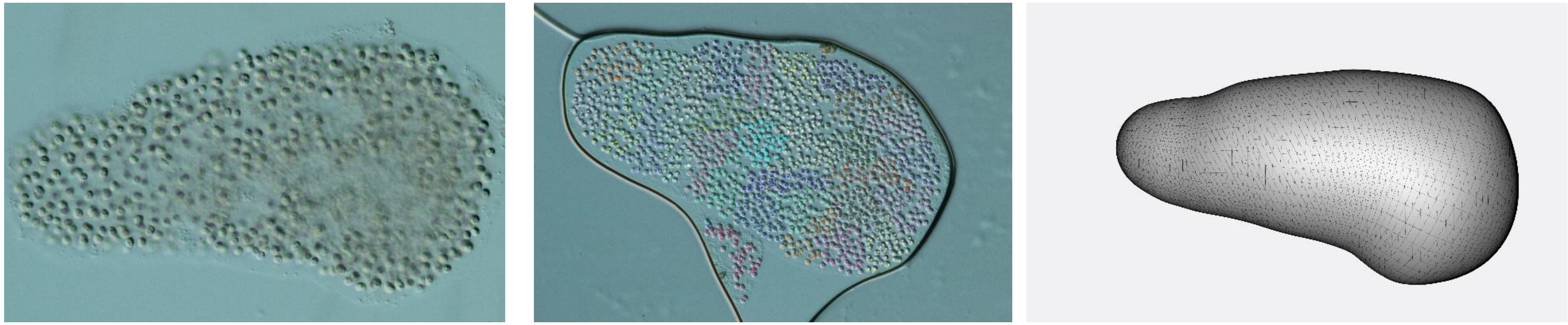

1558

27.

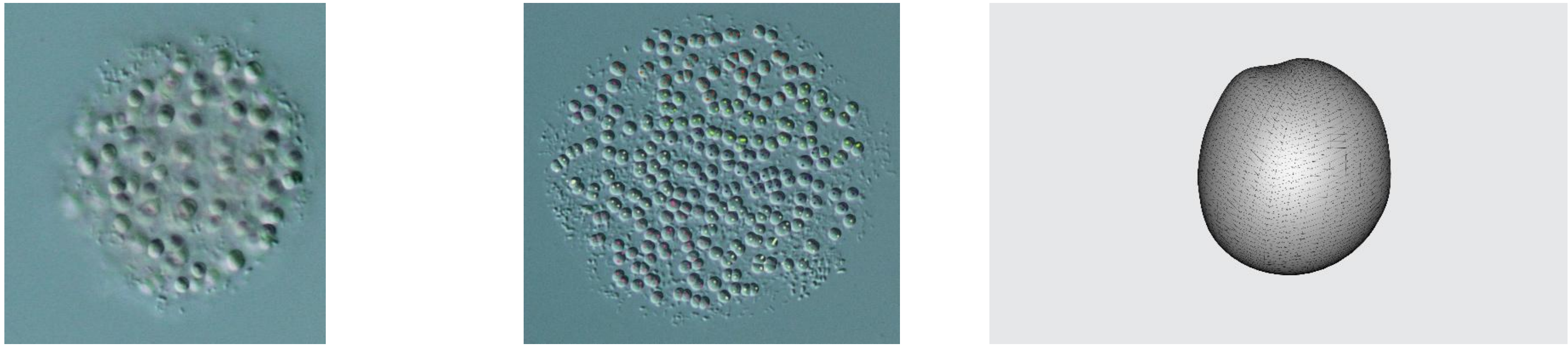

307

28.

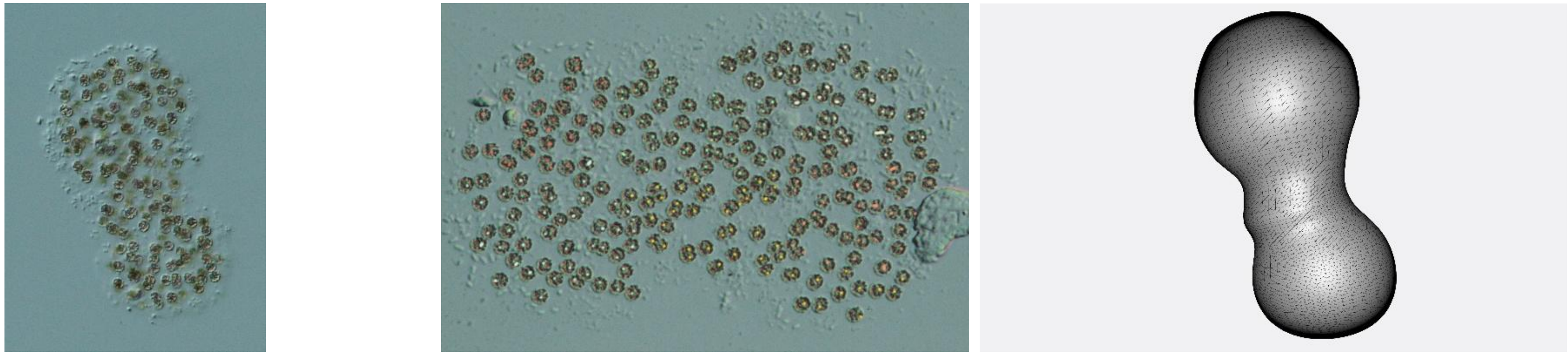

280

29.

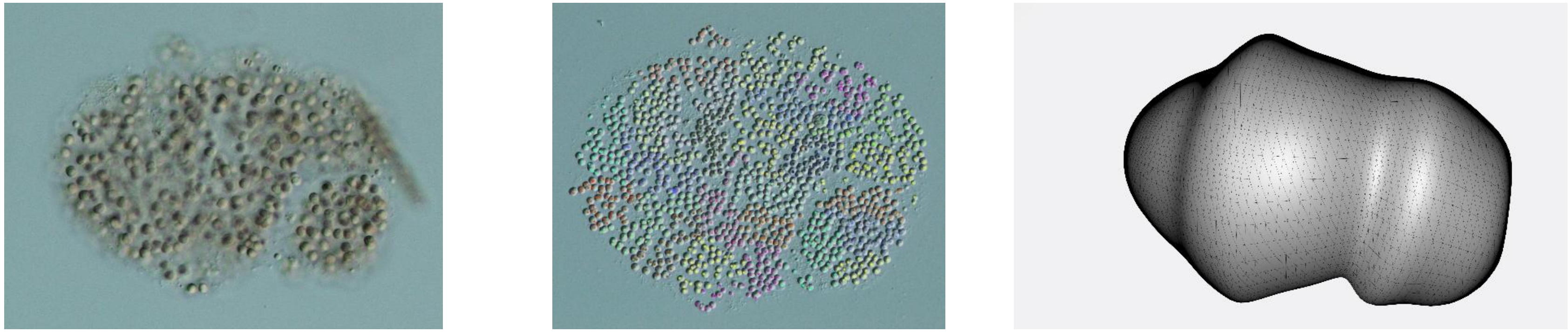

1278

30.

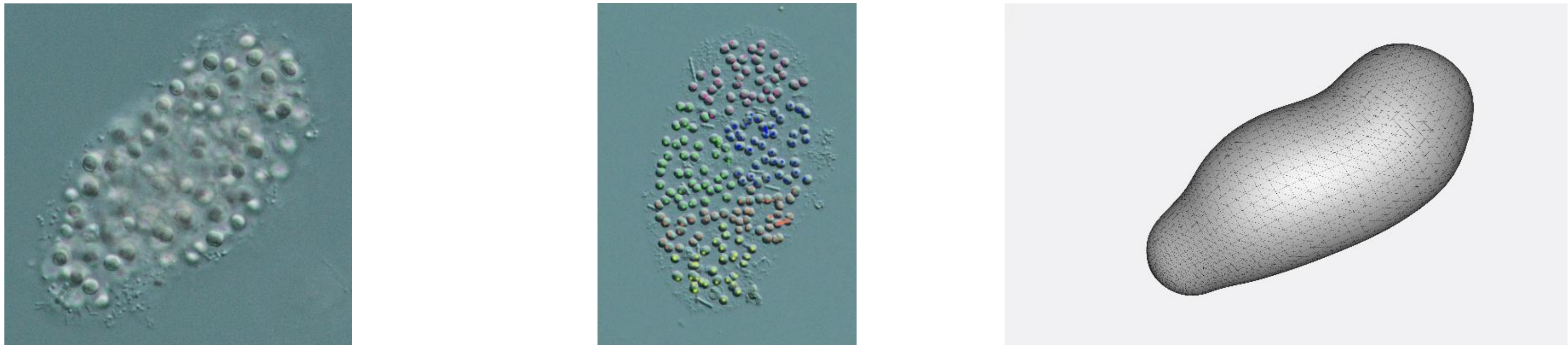

233

31.

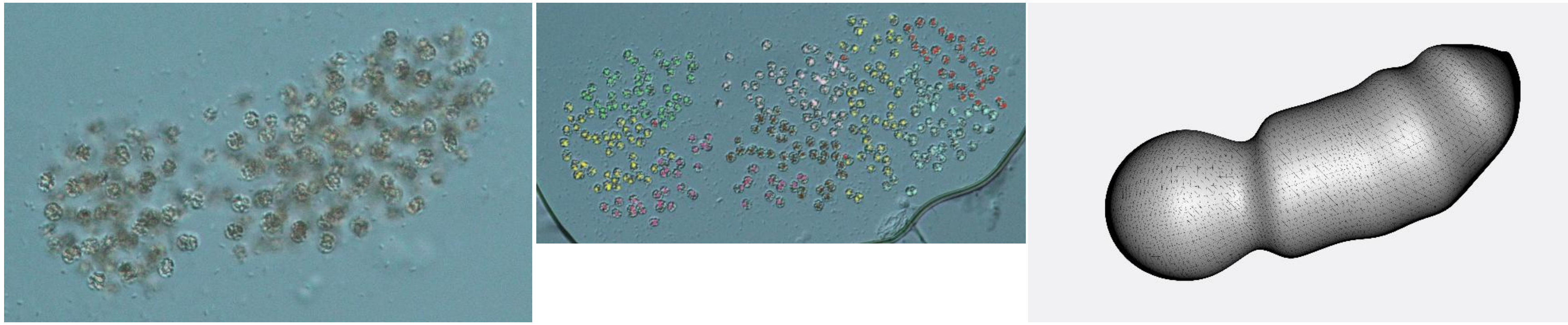

388

32.

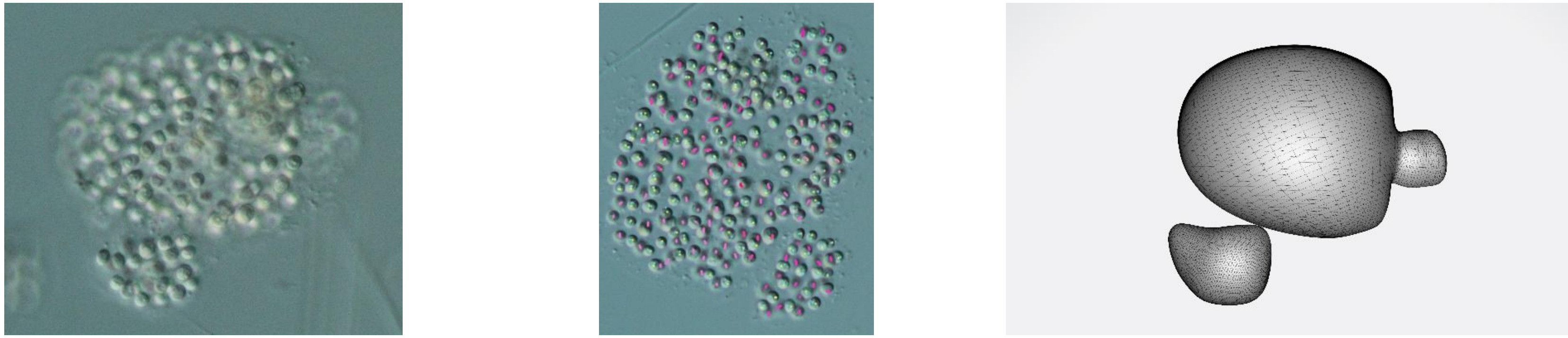

235

33.

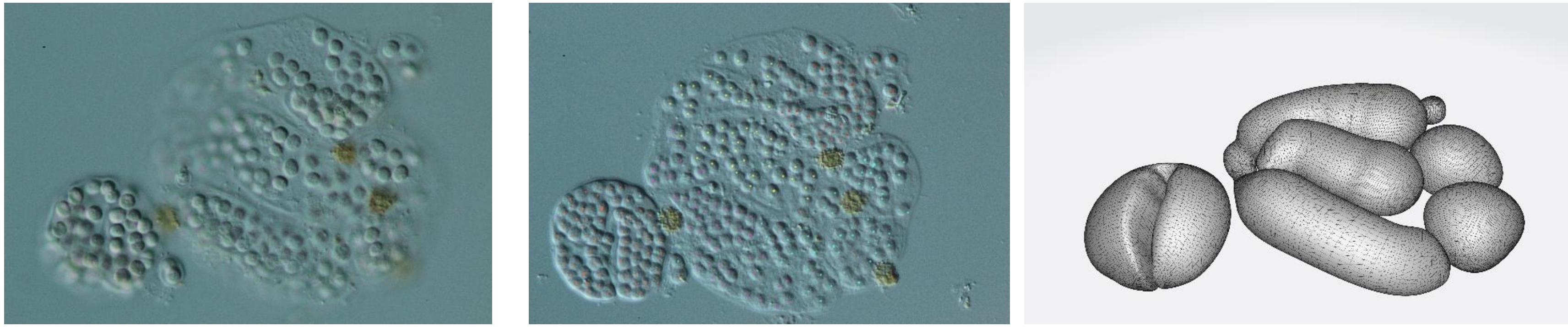

269

34.

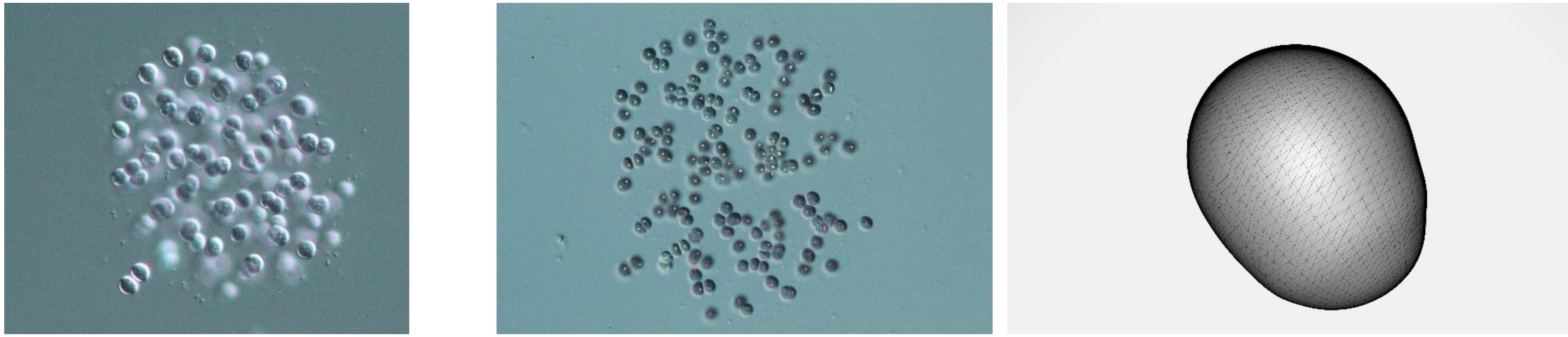

166

35.

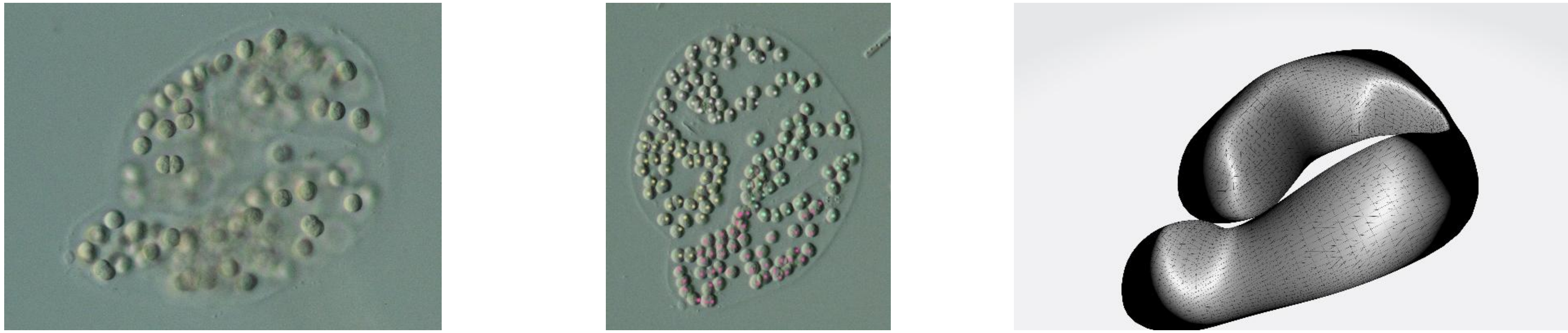

193

36.

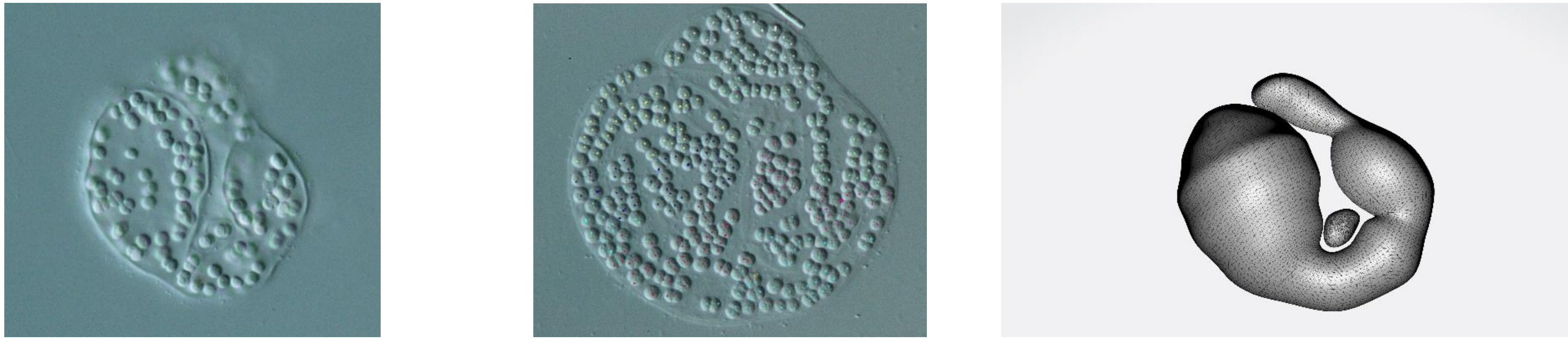

361

37.

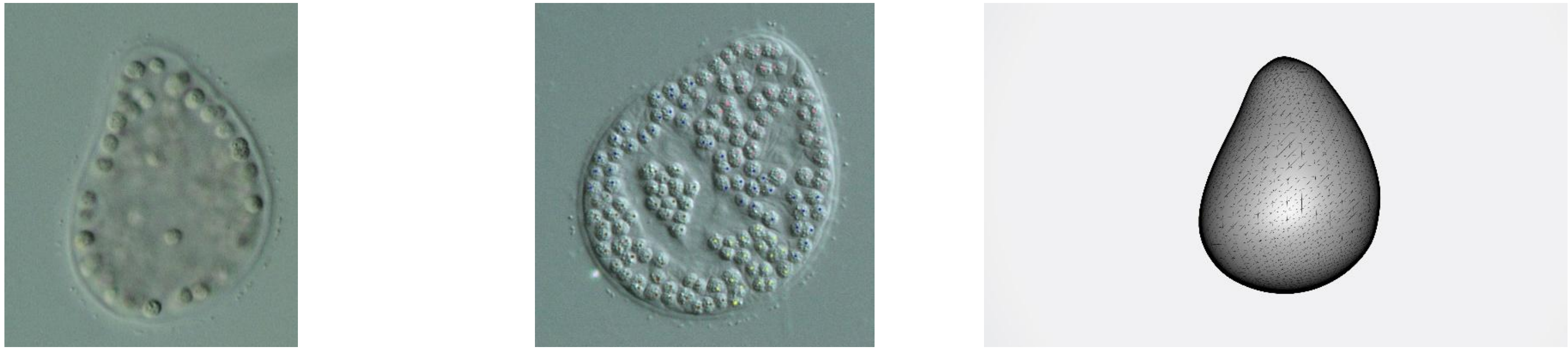

170

38.

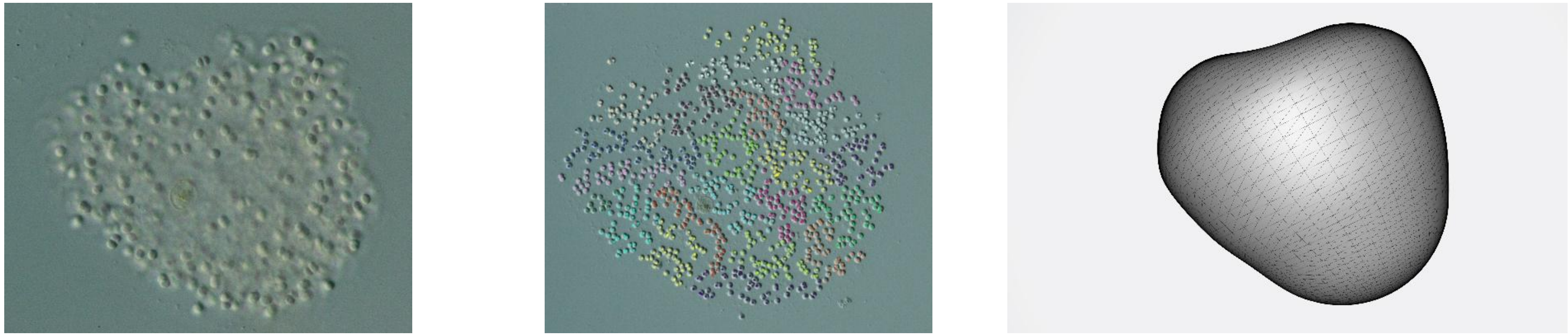

1062

39.

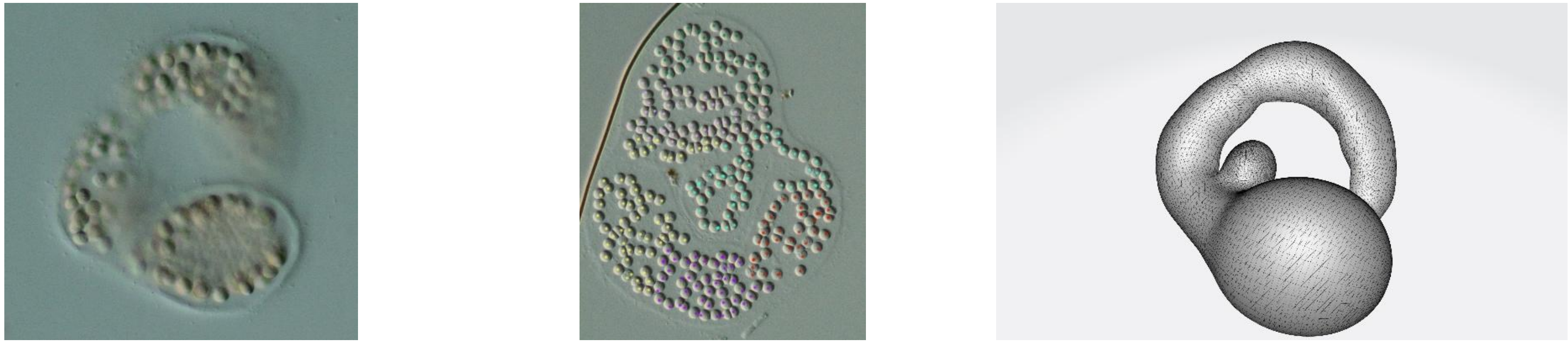

287

40.

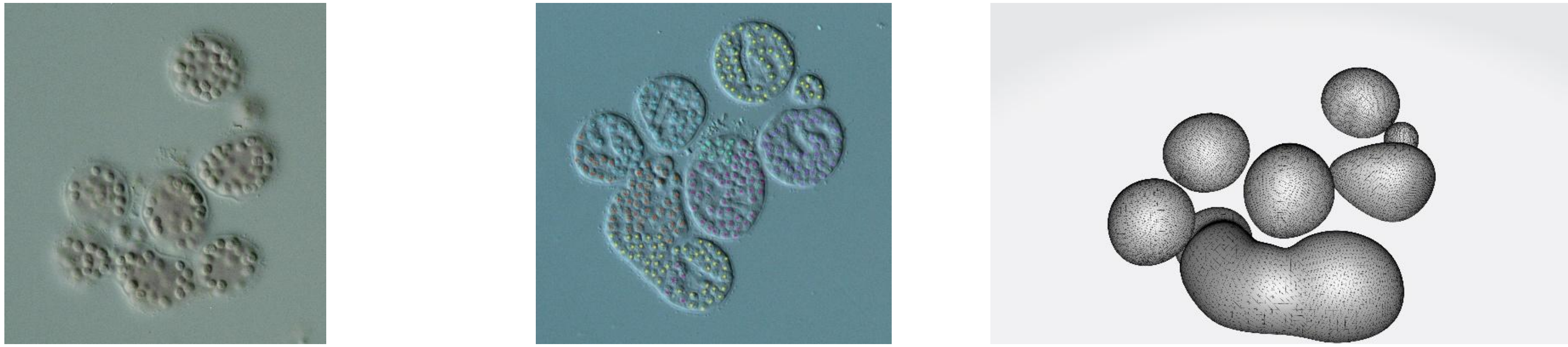

320

41.

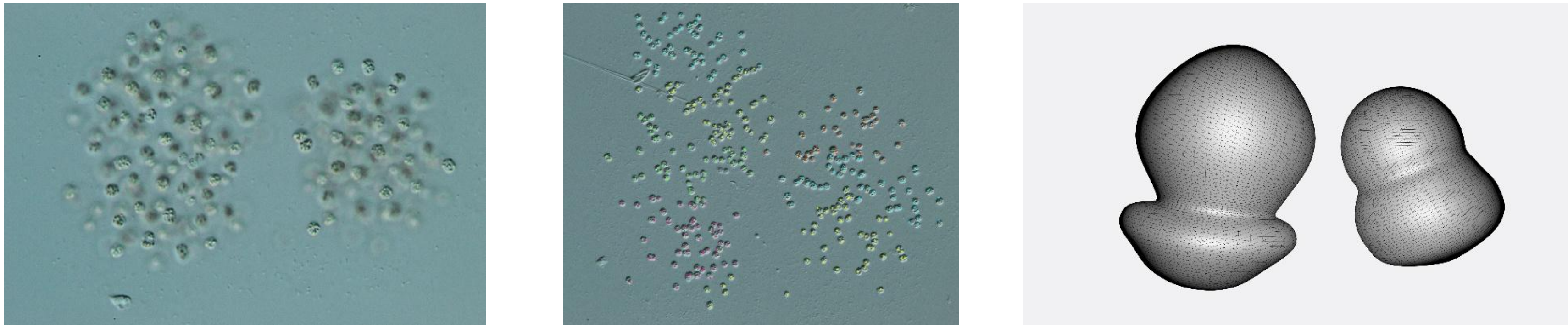

335

42.

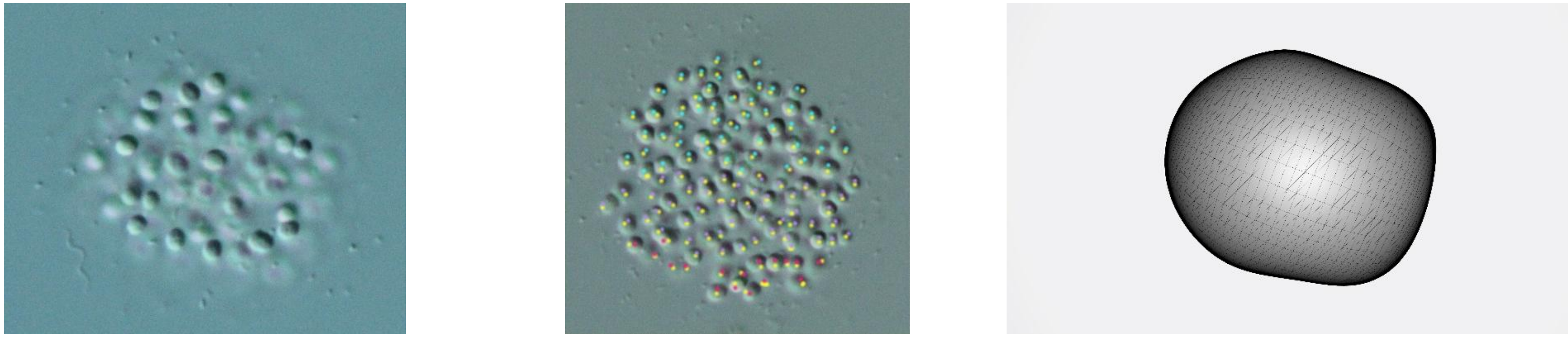

117

43.

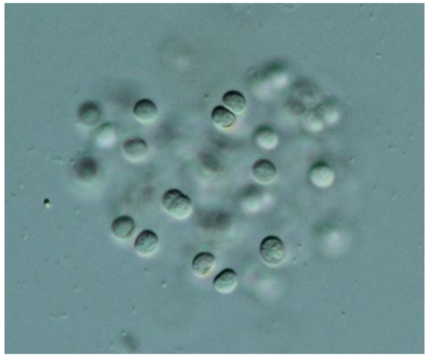

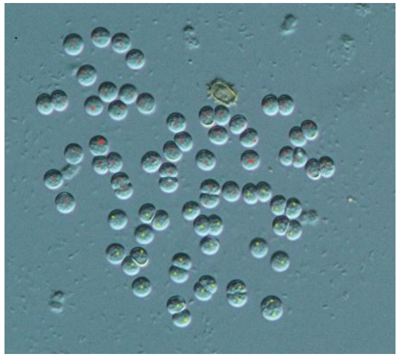

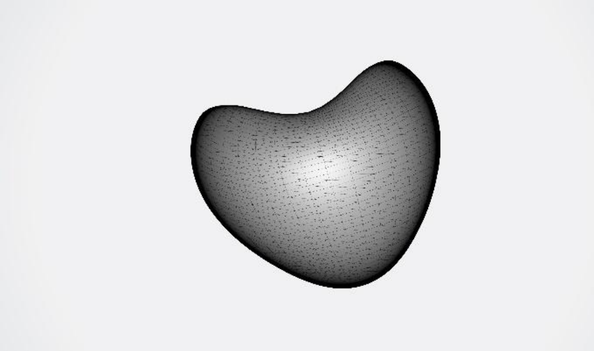

80

44.

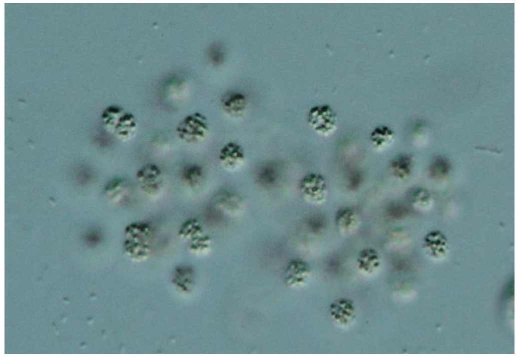

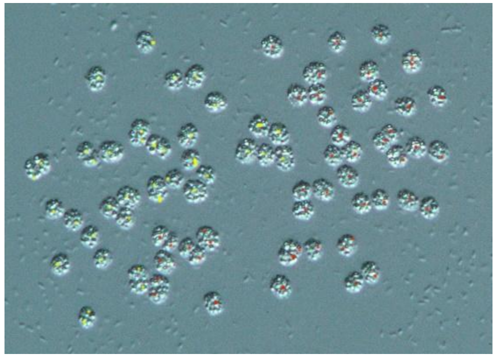

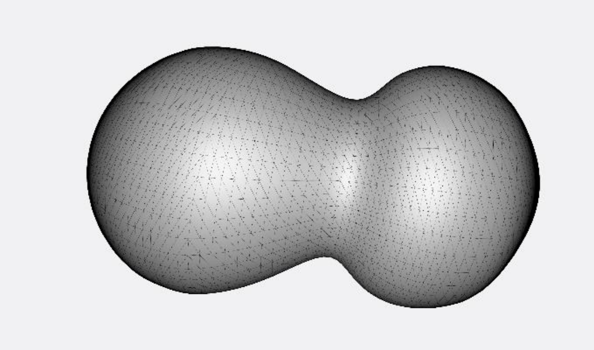

90

45.

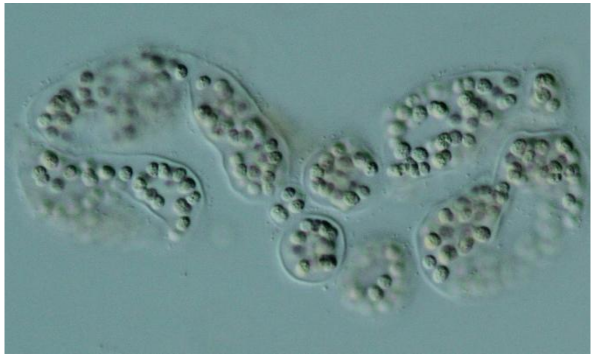

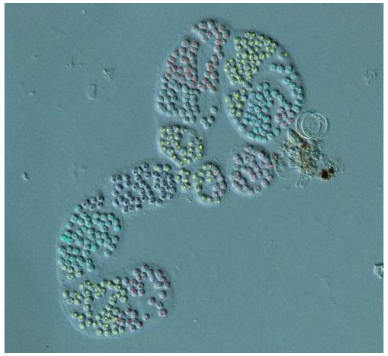

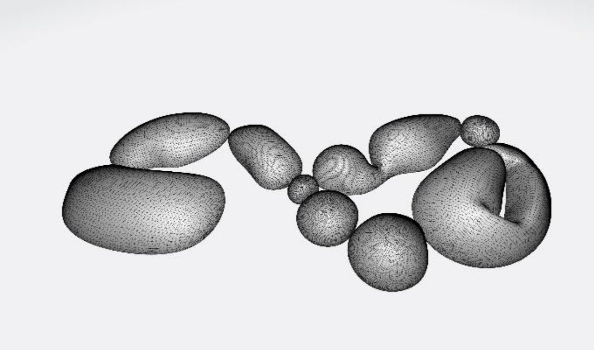

432

46.

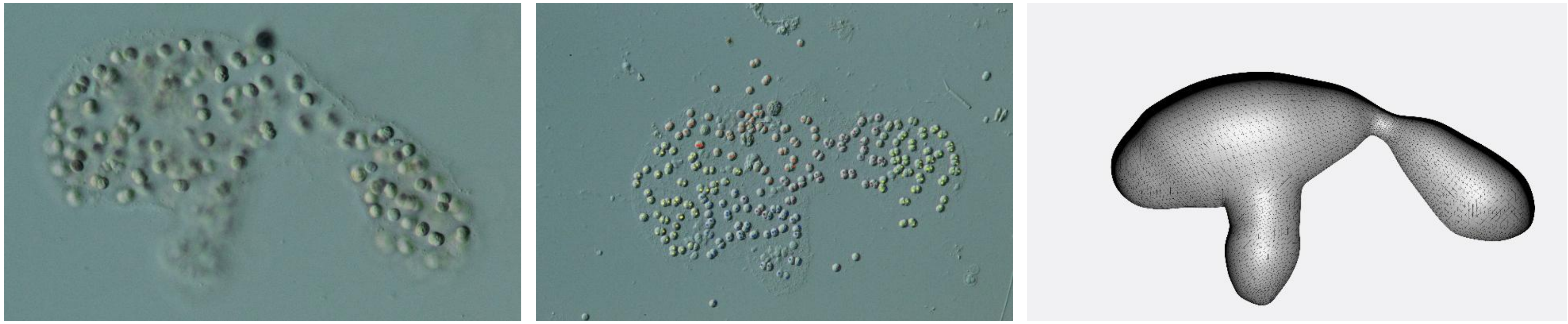

264

47.

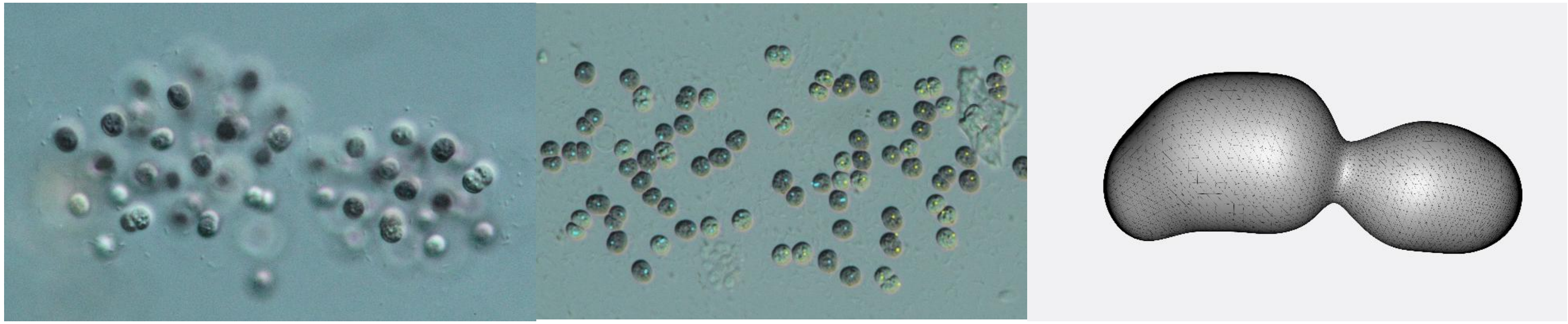

90

48.

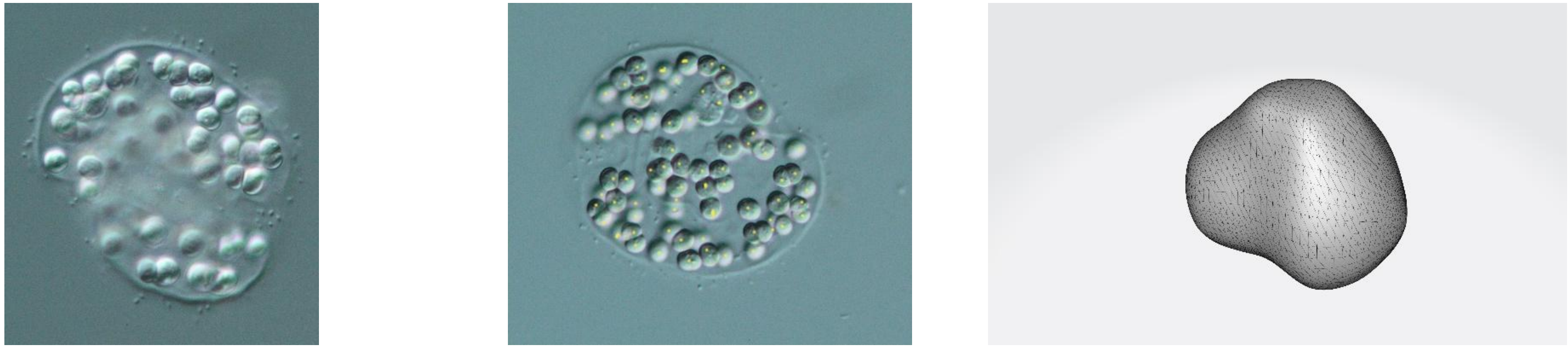

71

49.

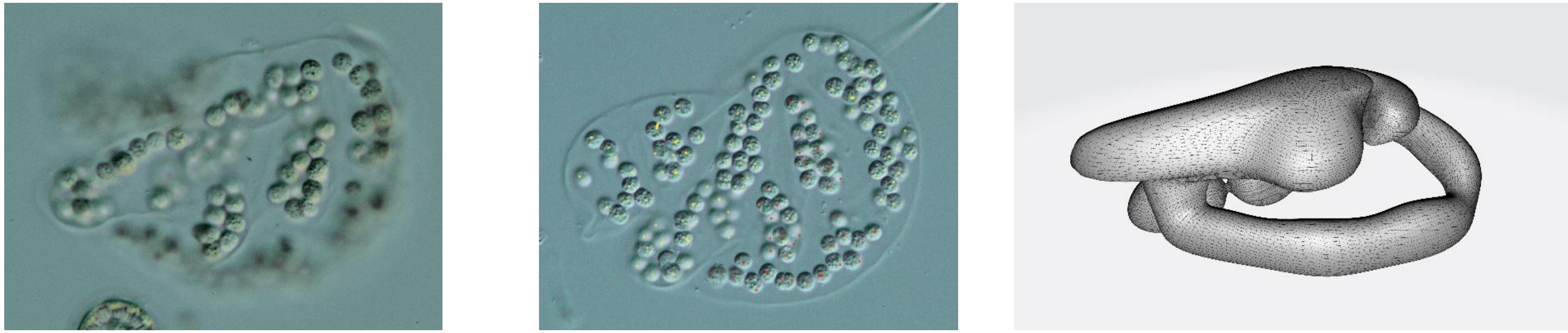

119

50.

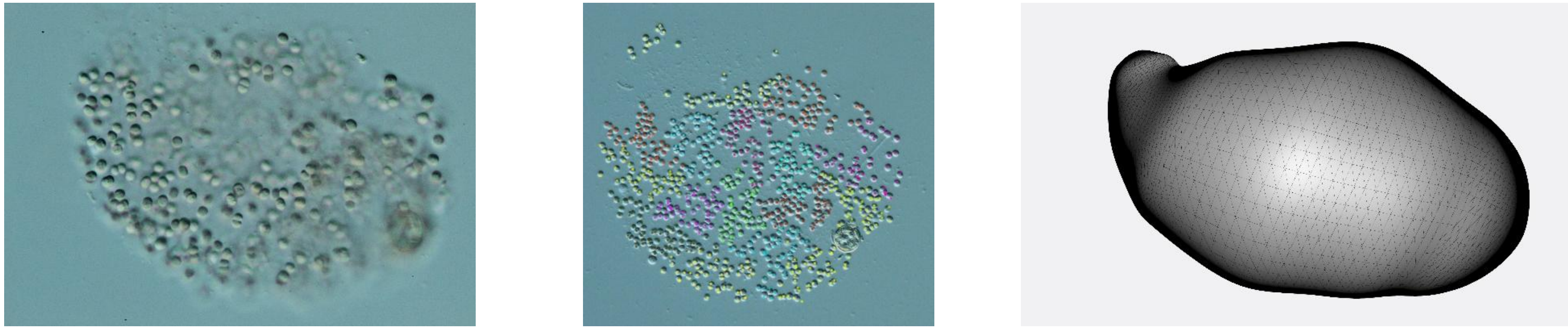

800

51.

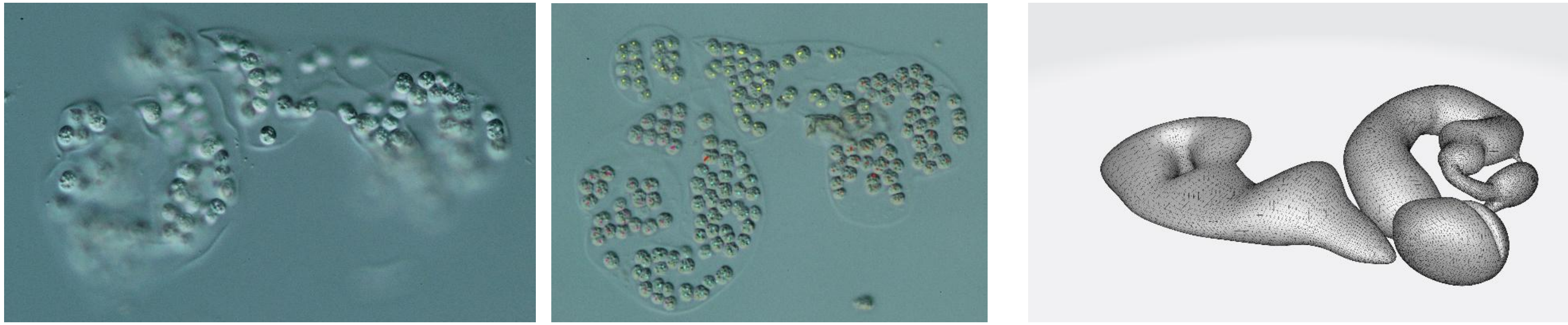

183

52.

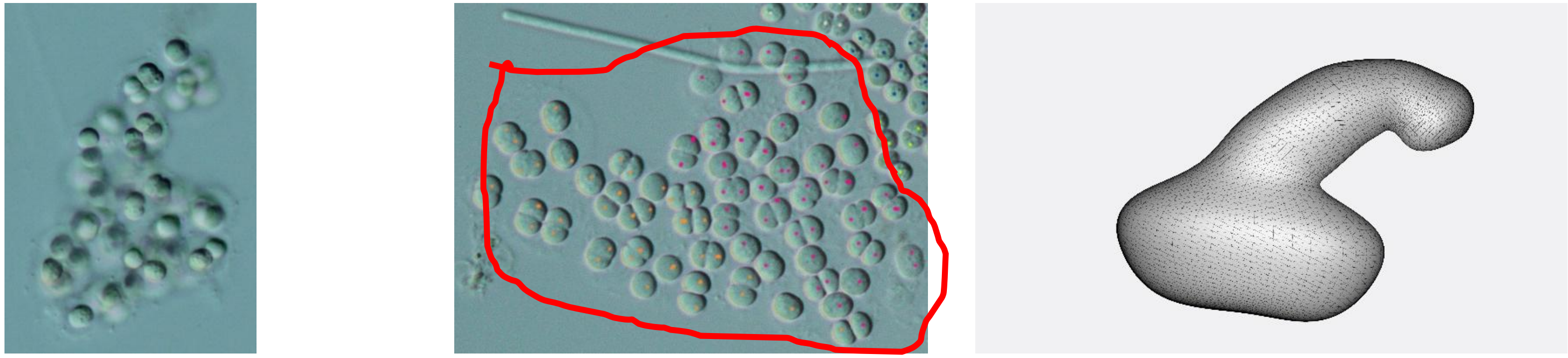

89

53.

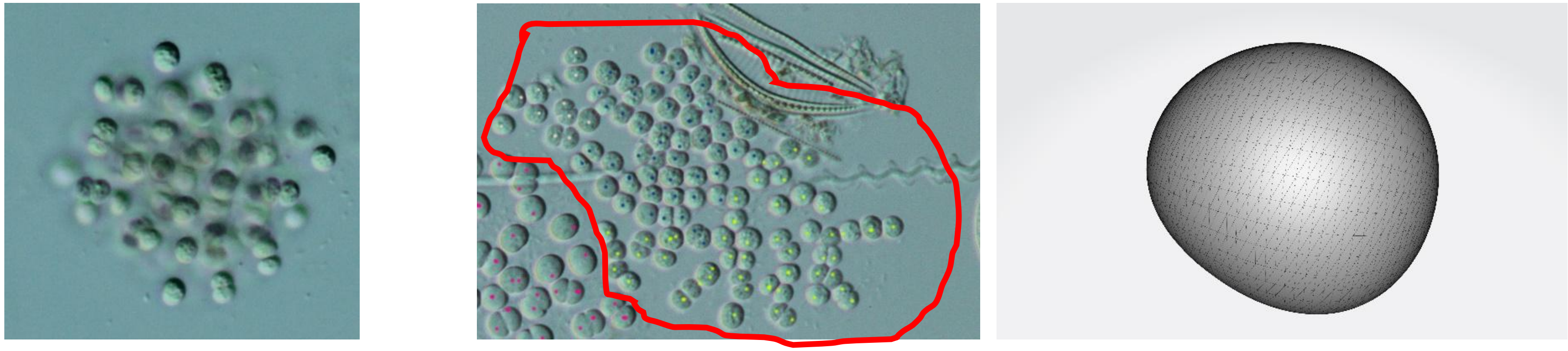

112

54.

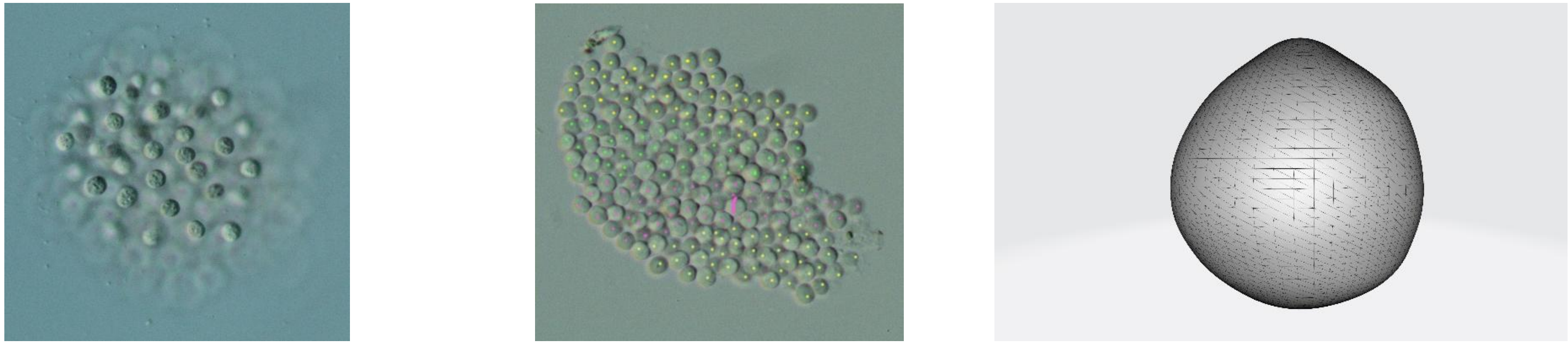

193

55.

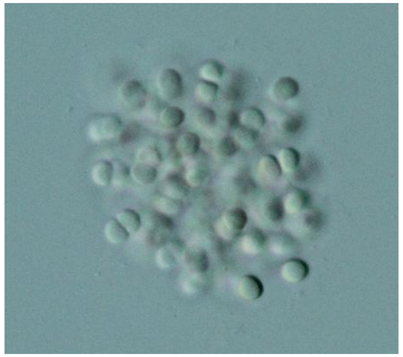

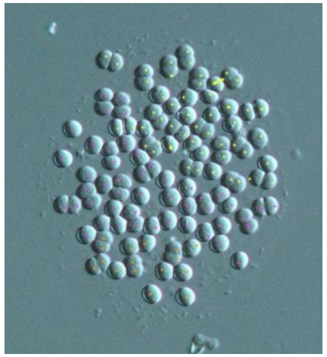

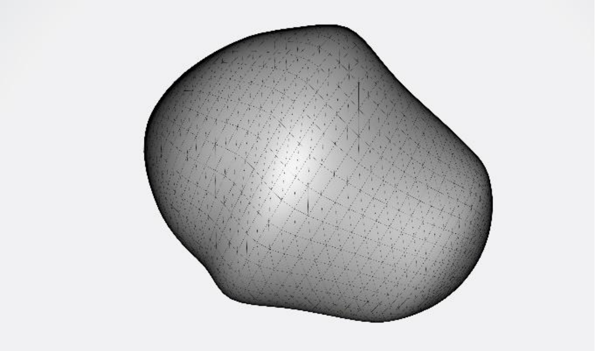

116

56.

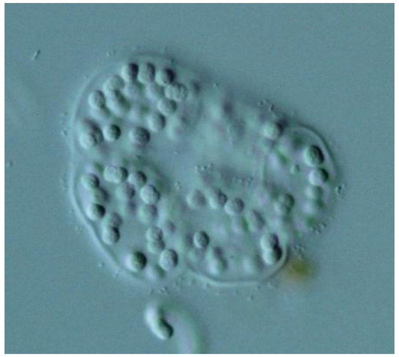

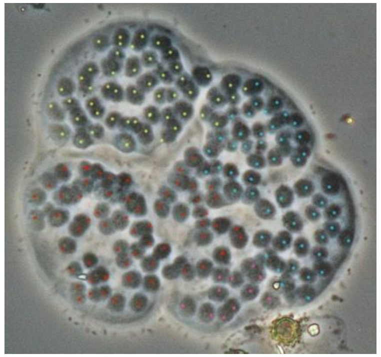

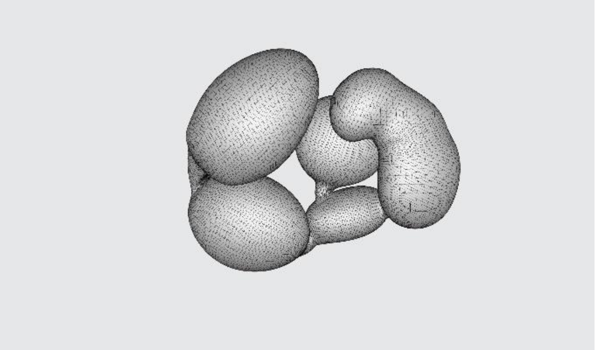

167

57.

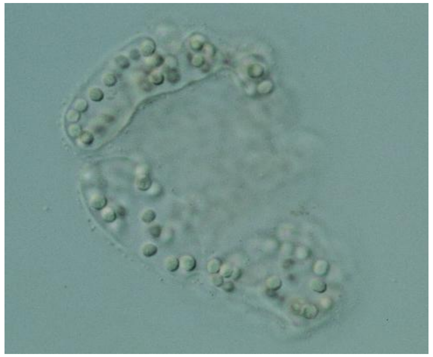

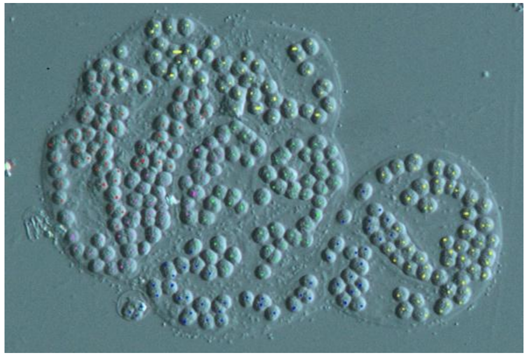

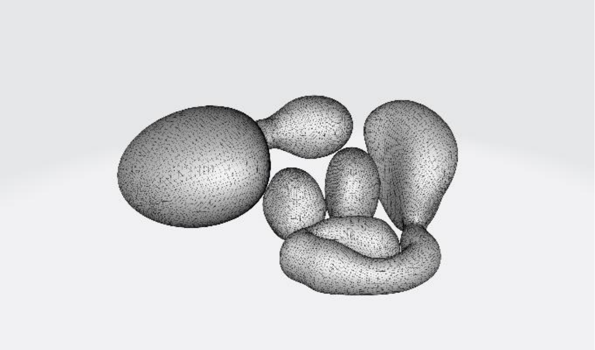

307

58.

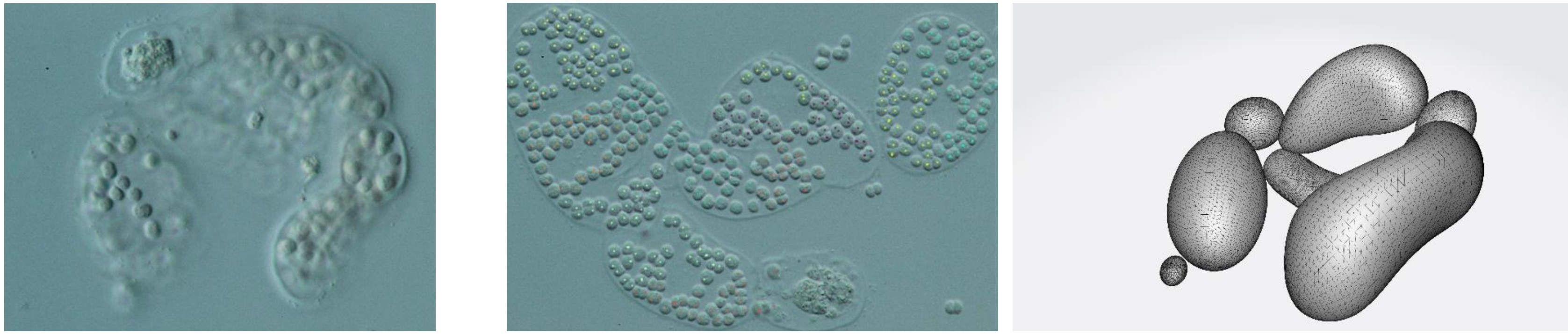

395

59.

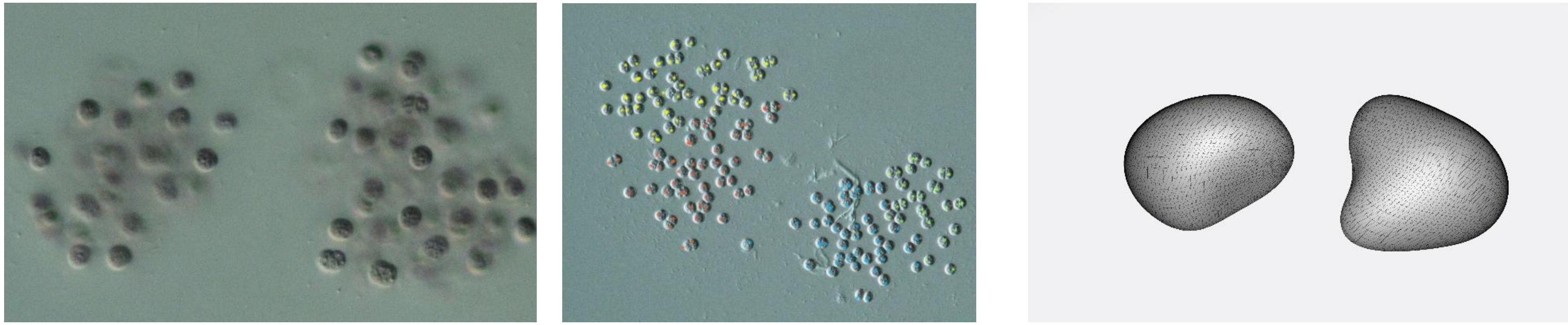

117

60.

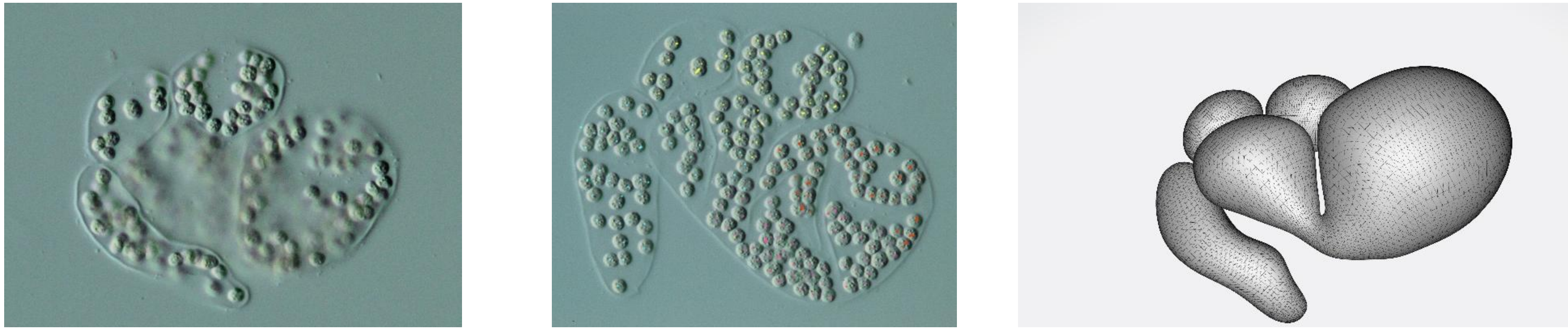

191

61.

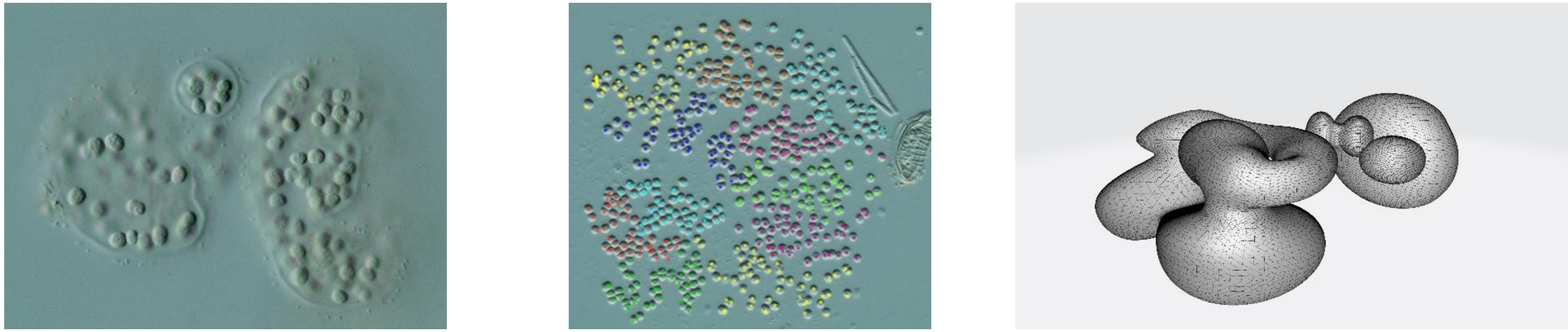

541

62.

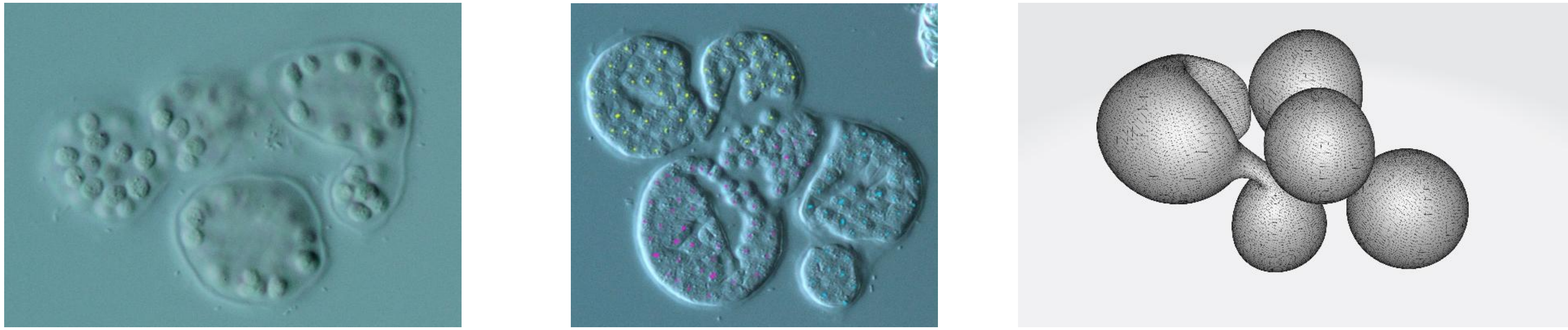

132

63.

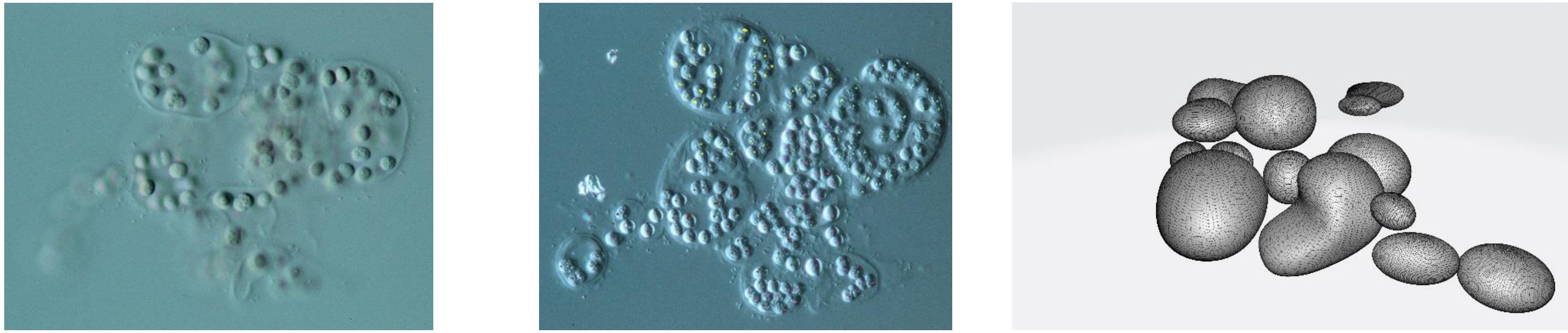

197

64.

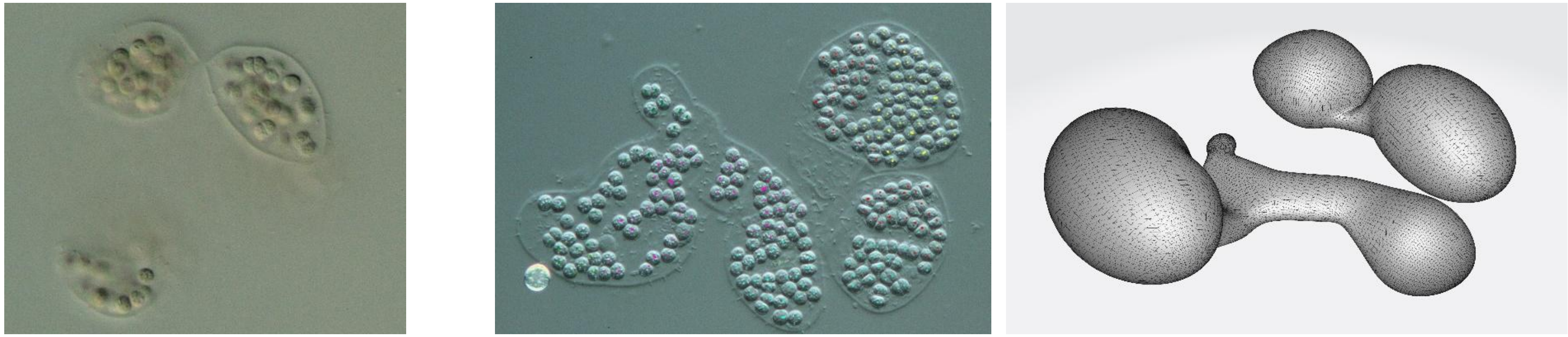

231

65.

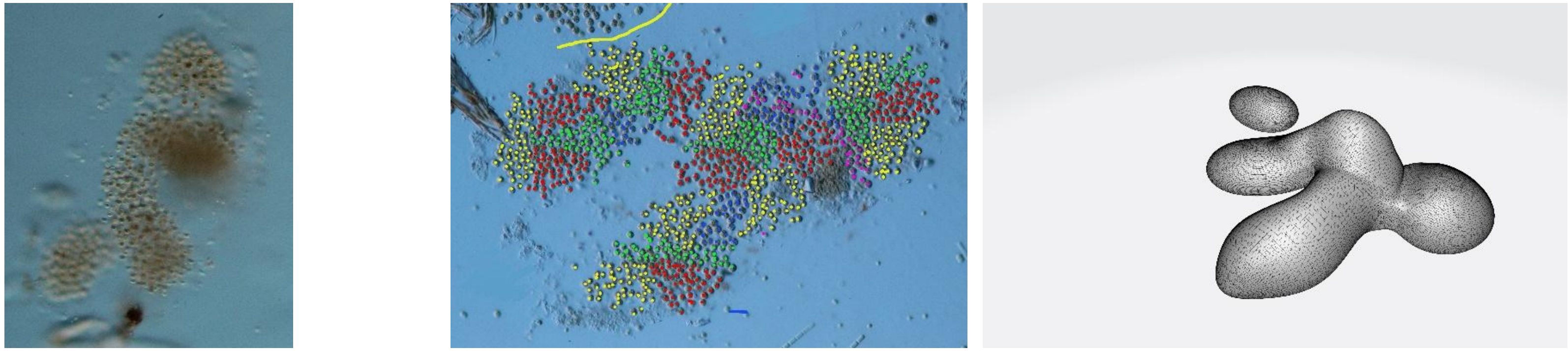

1355

66.

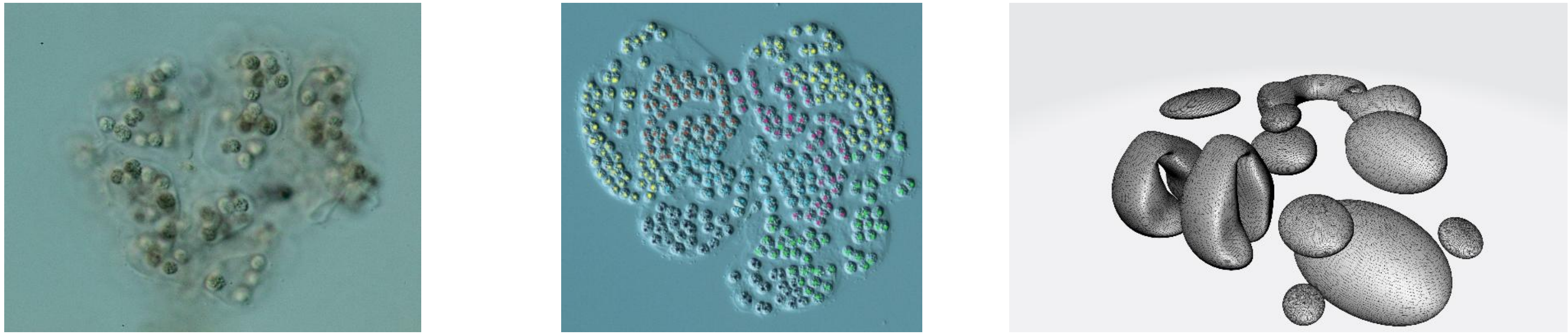

342

67.

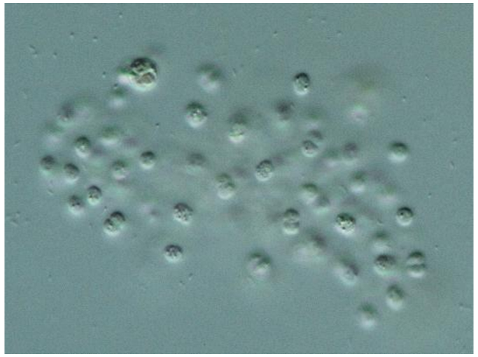

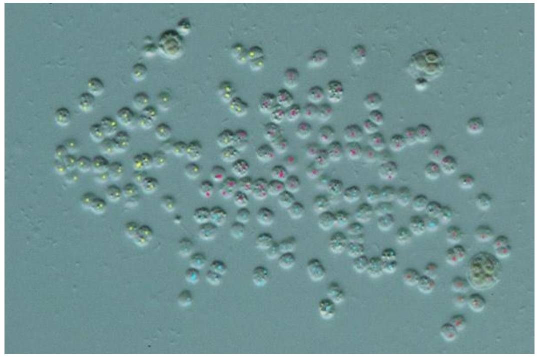

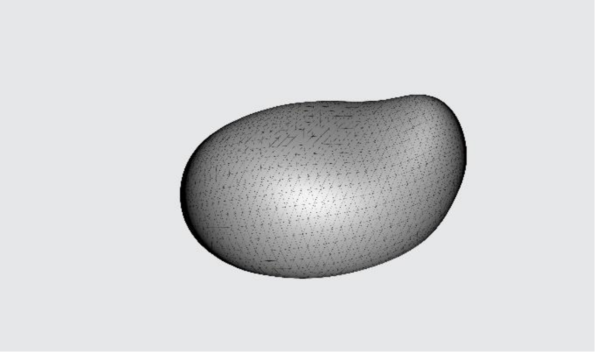

164

68.

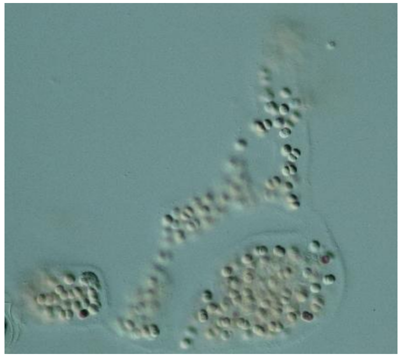

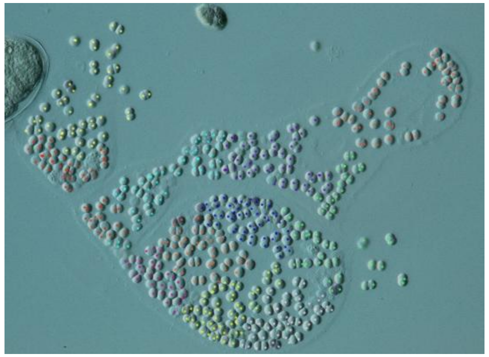

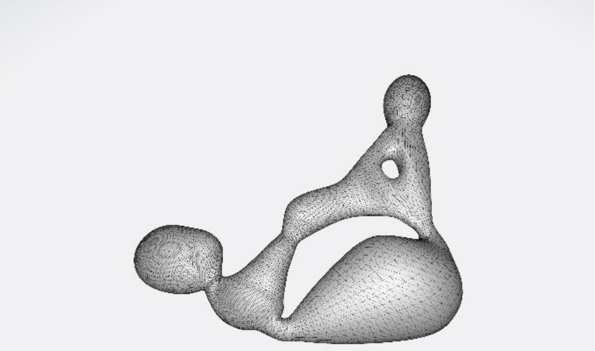

532

69.

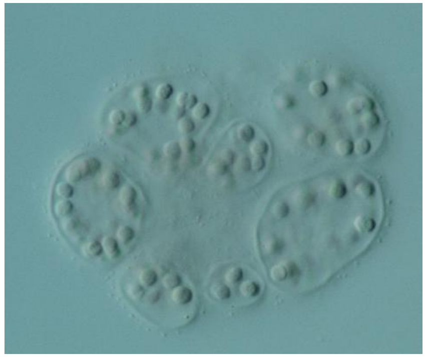

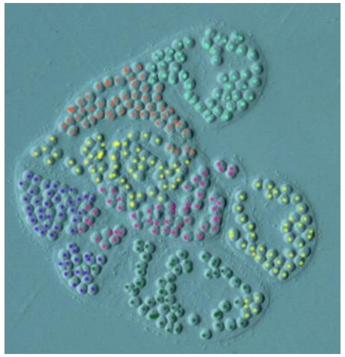

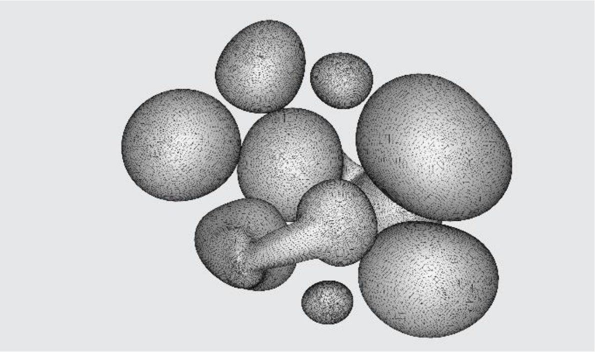

343

70. 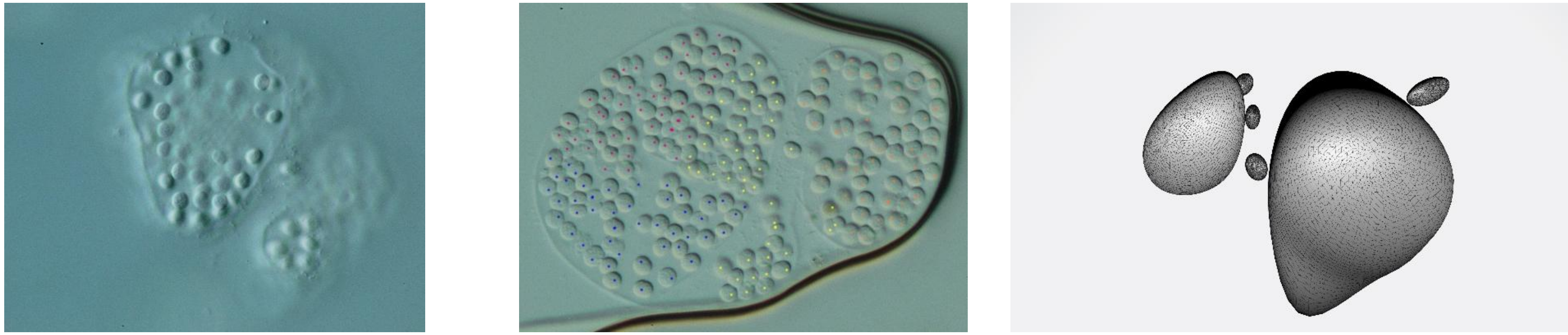 196

71. 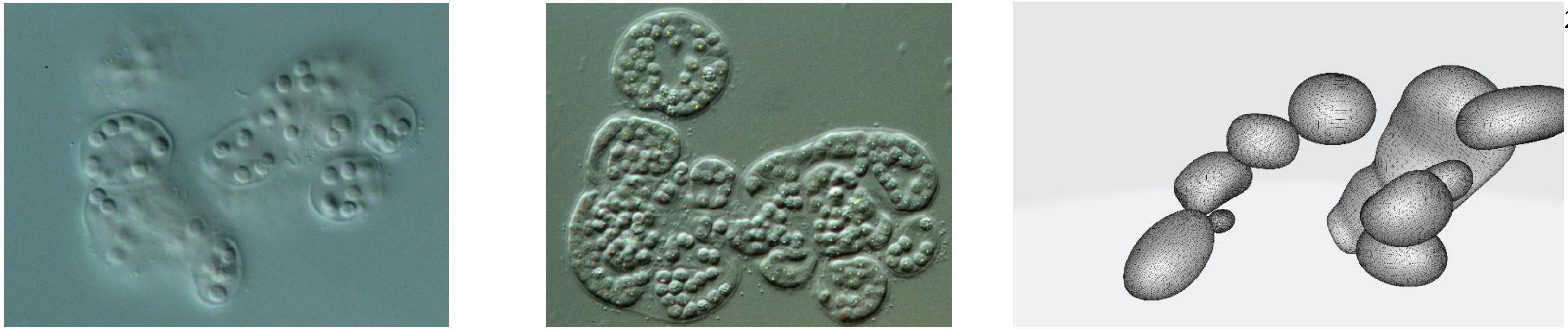 260

72. 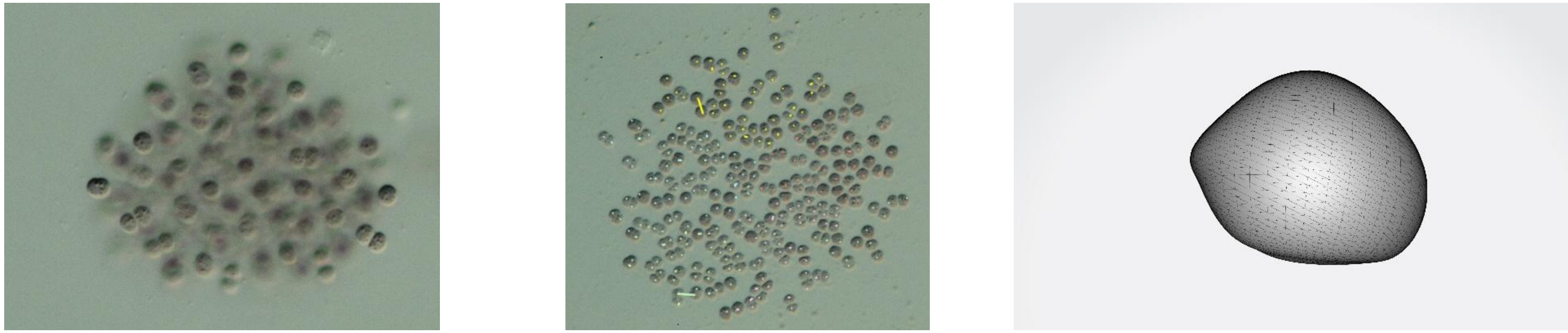 267

73.

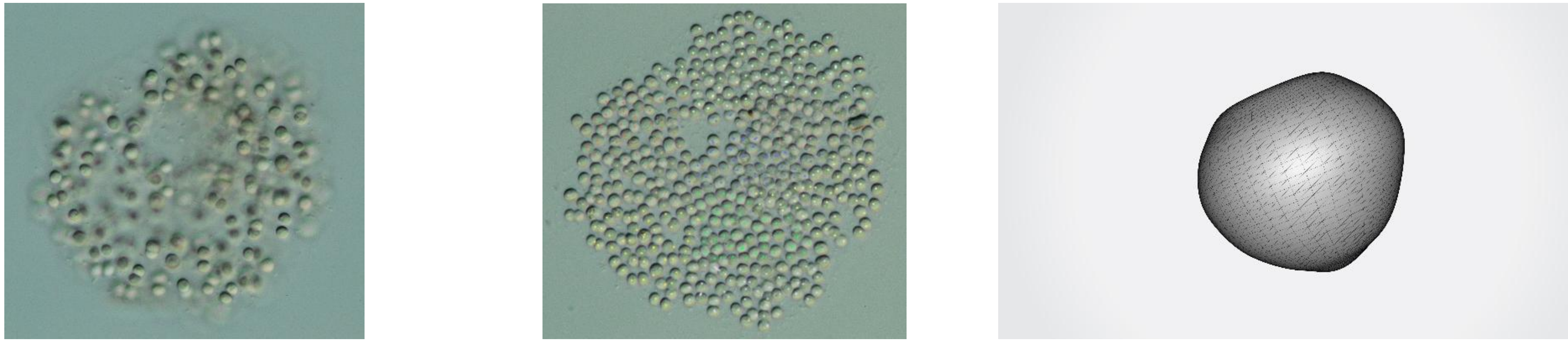

495

74.

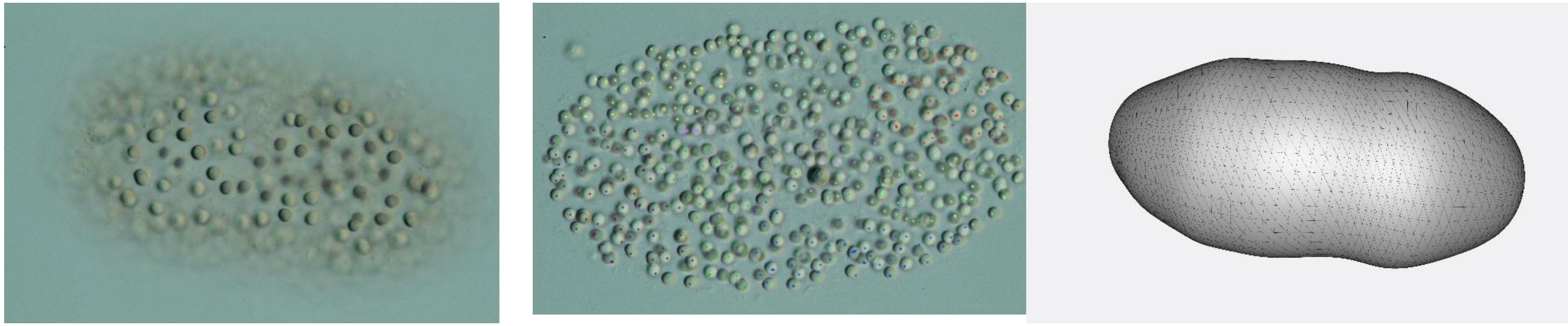

414

75.

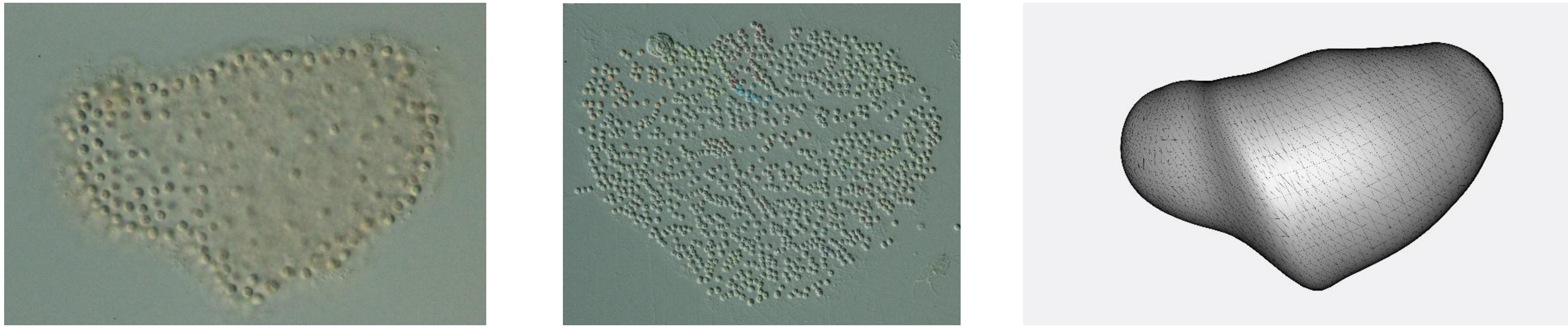

1530

76.

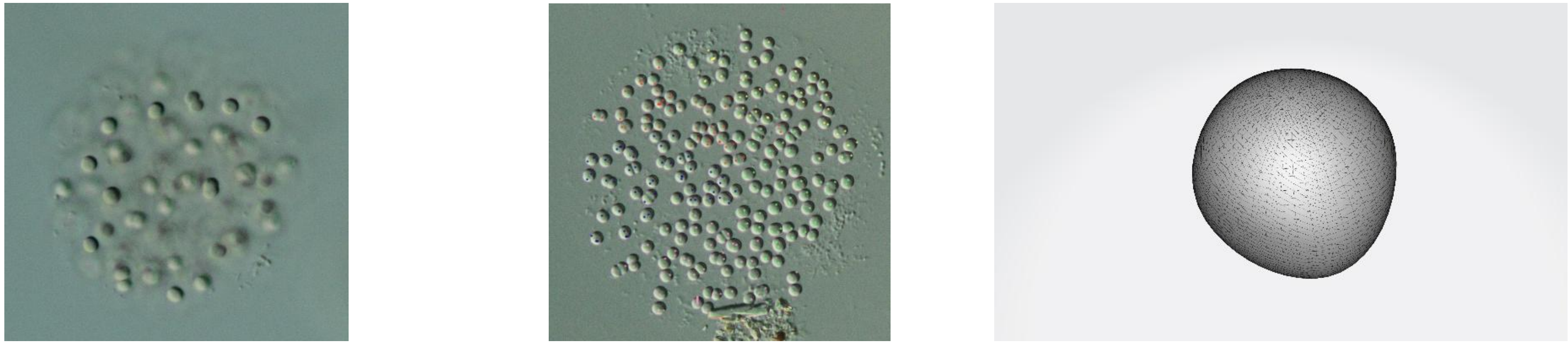

245

77.

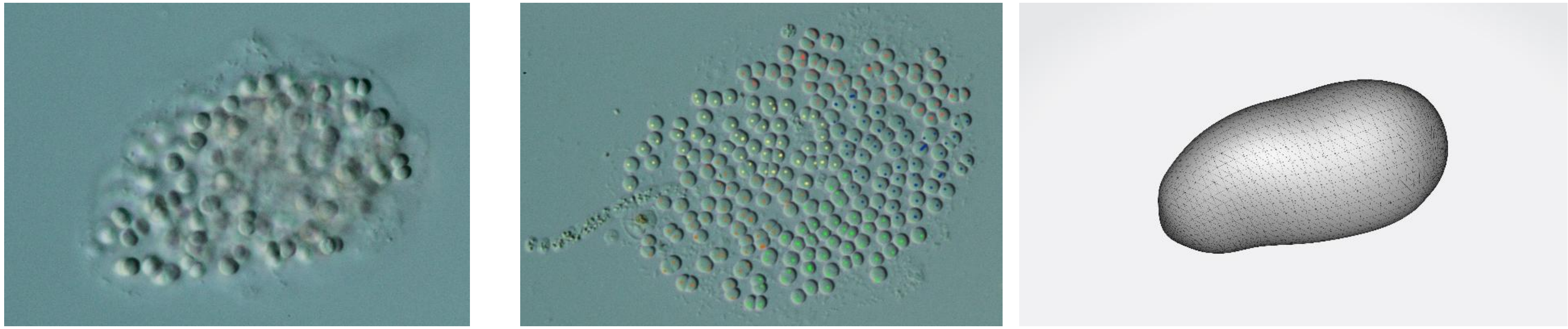

246

78.

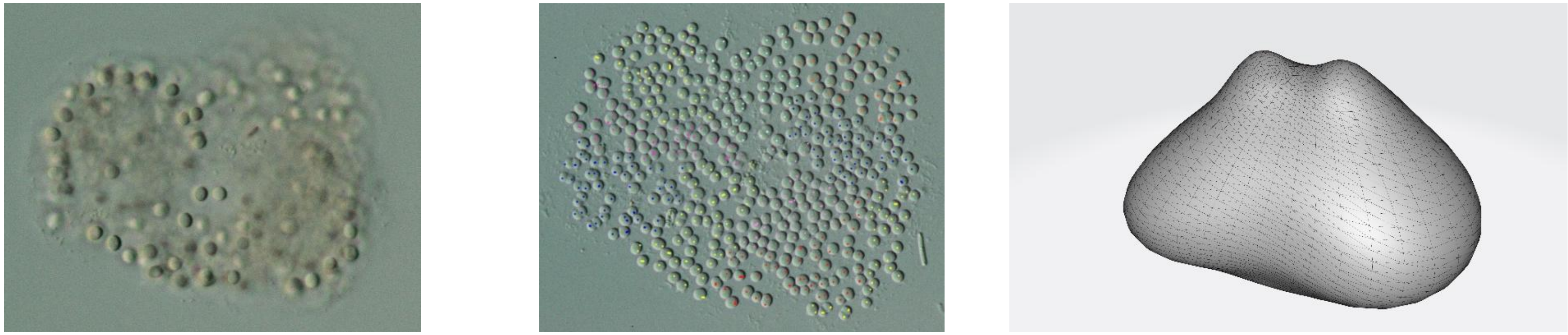

482

79.

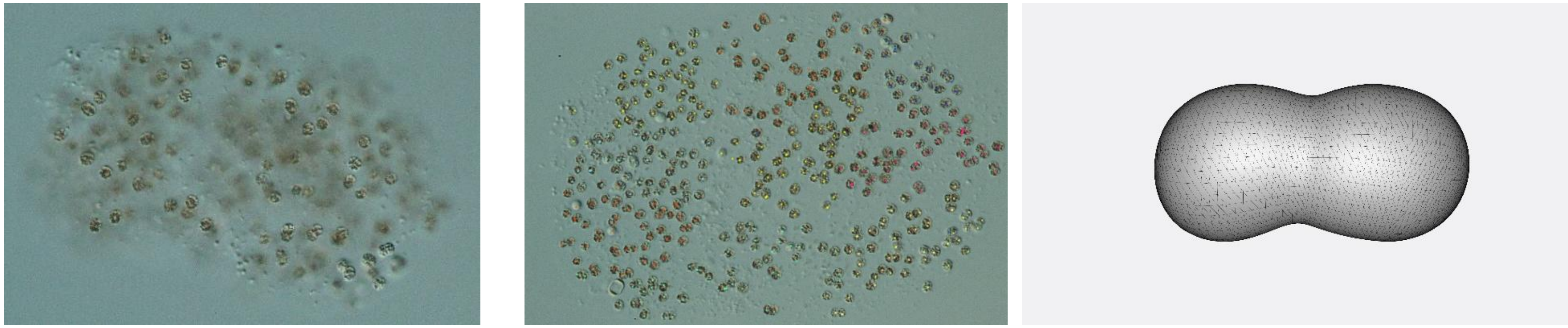

422

80.

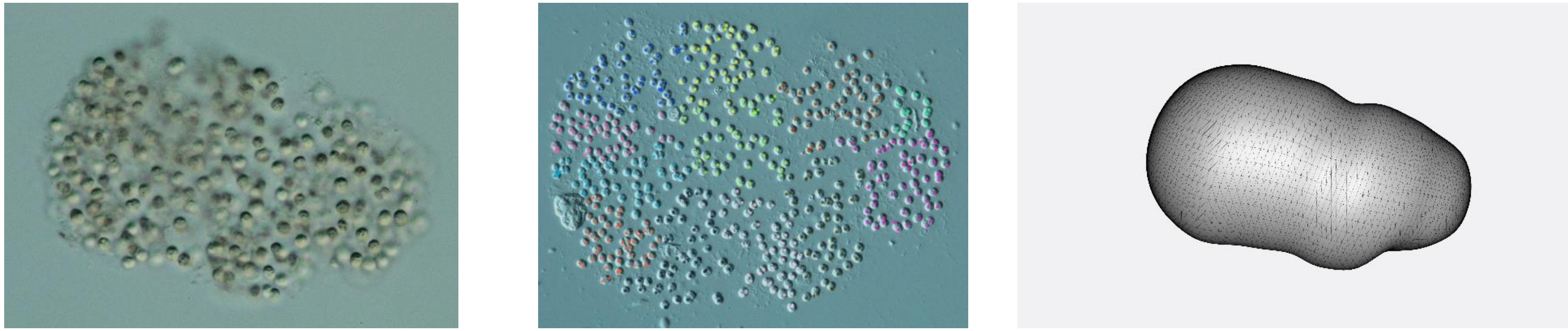

574

81.

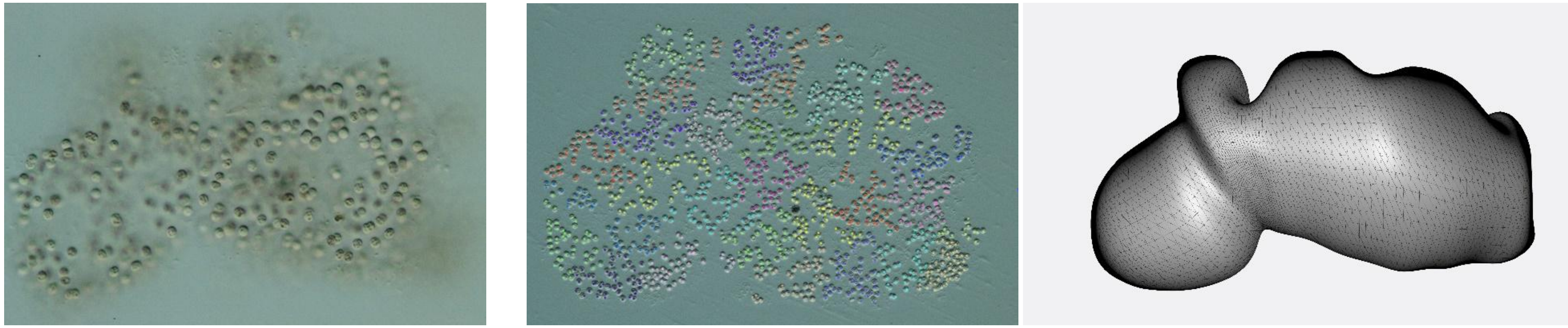

1312

82.

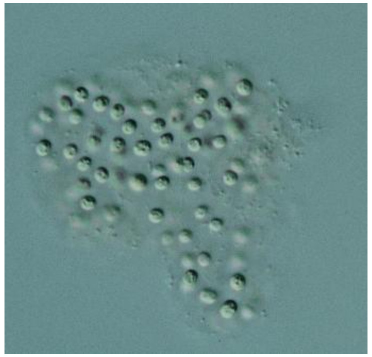

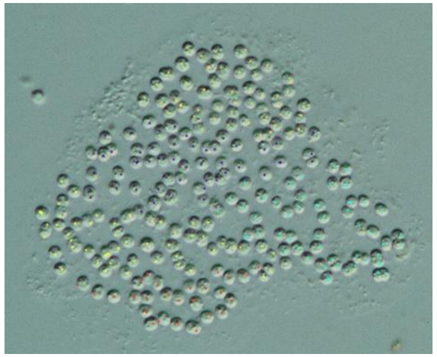

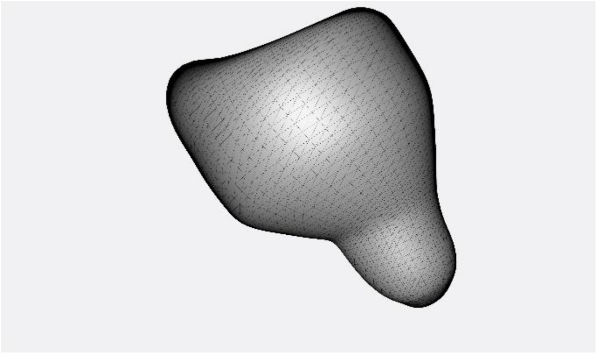

228

83.

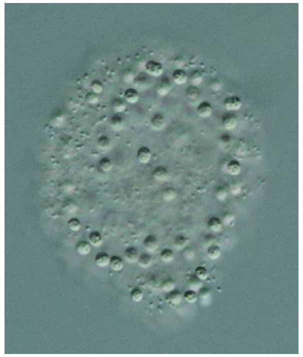

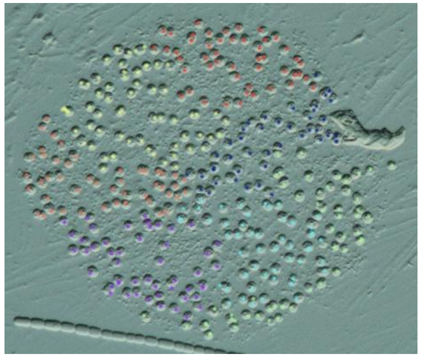

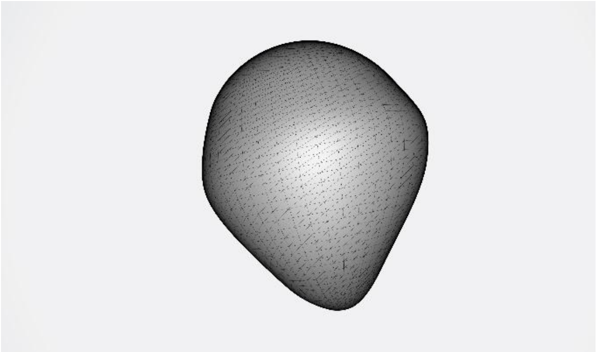

351

84.

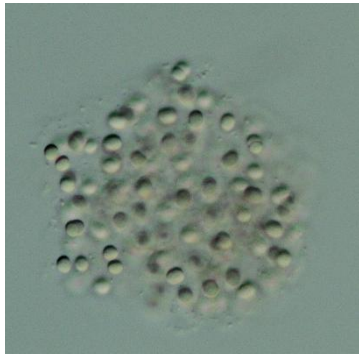

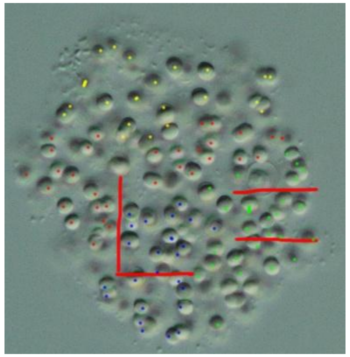

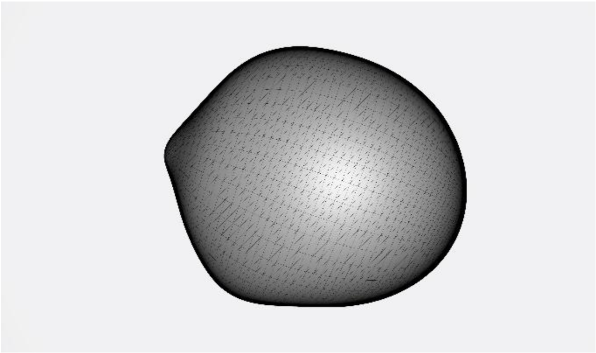

189

85.

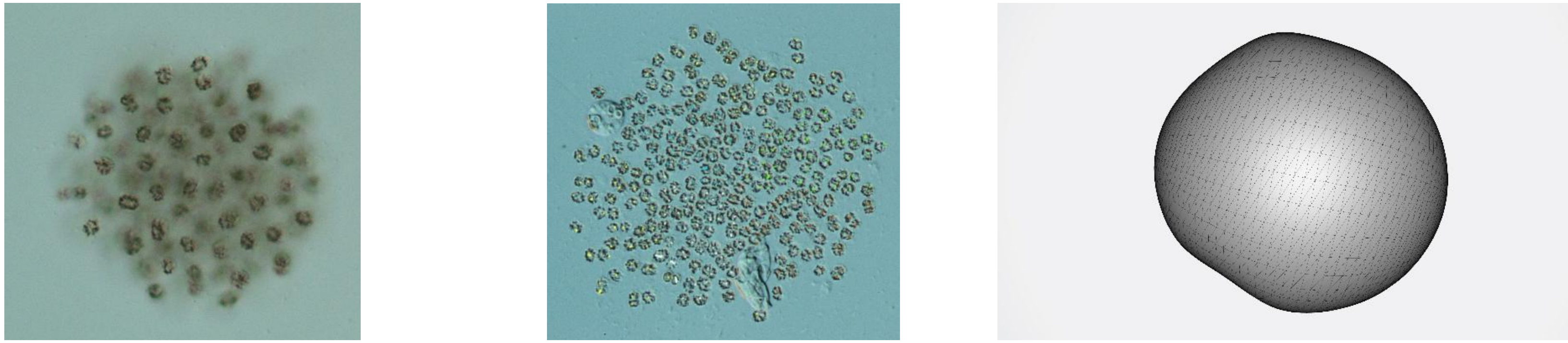

292

86.

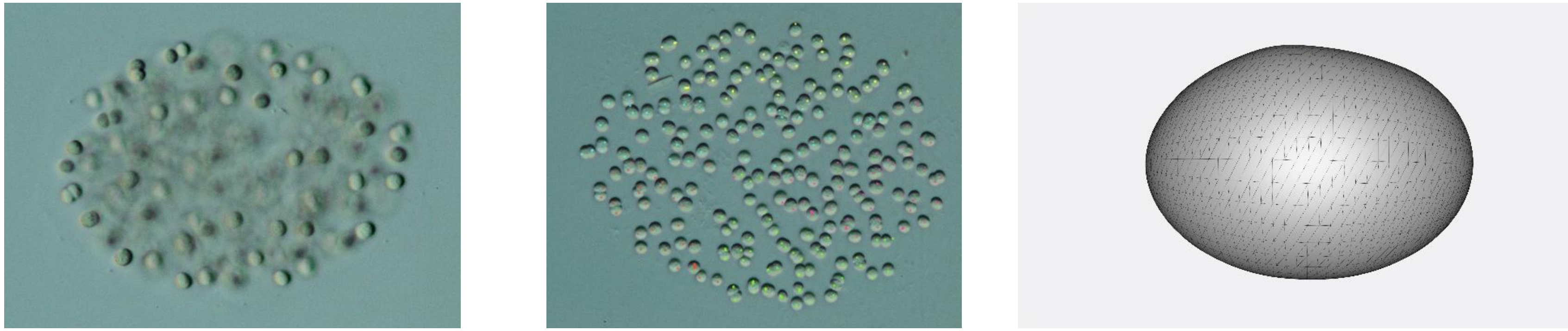

232

87.

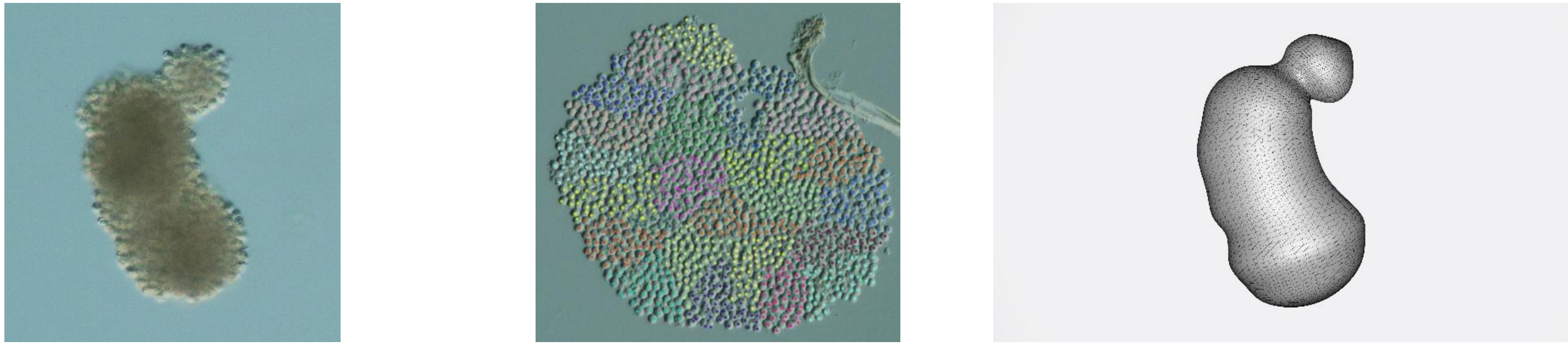

1259

88.

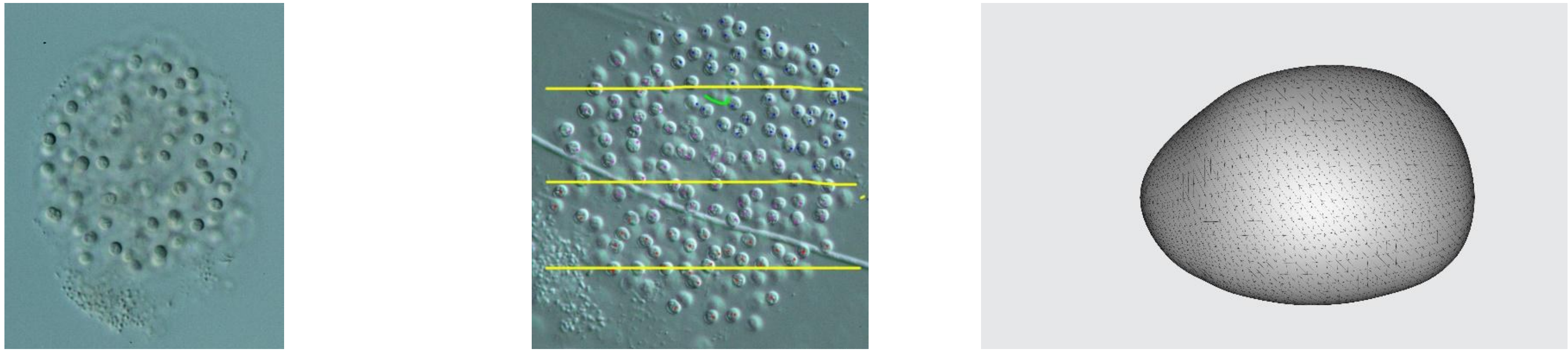

228

89.

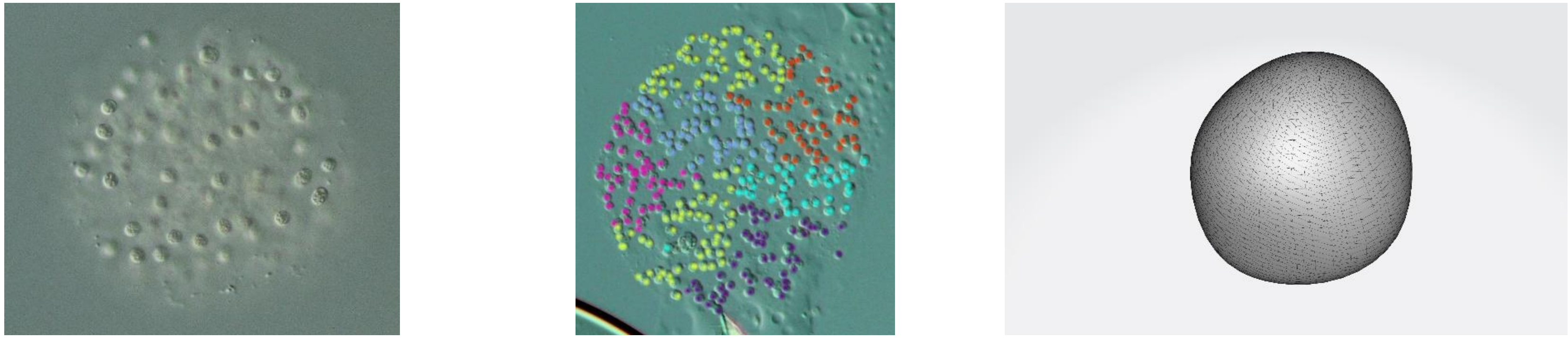

339

90.

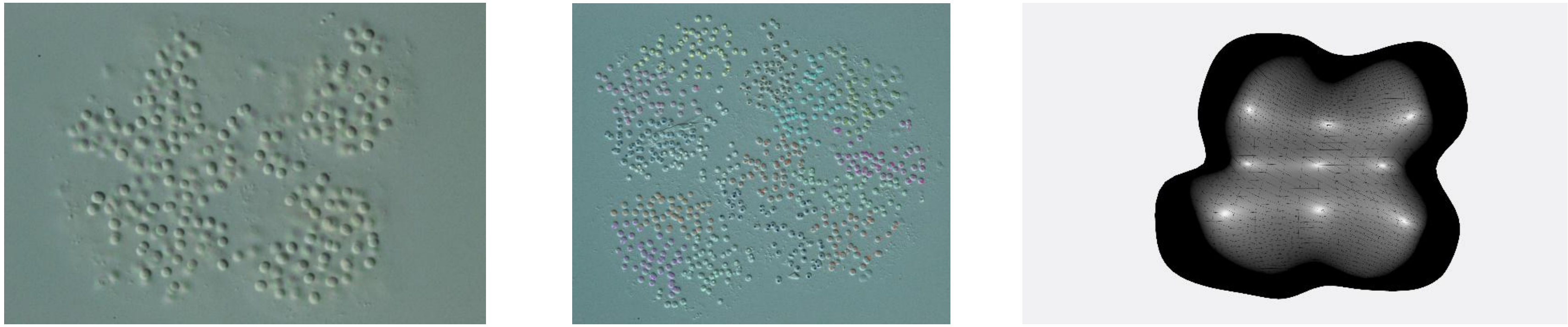

746

91.

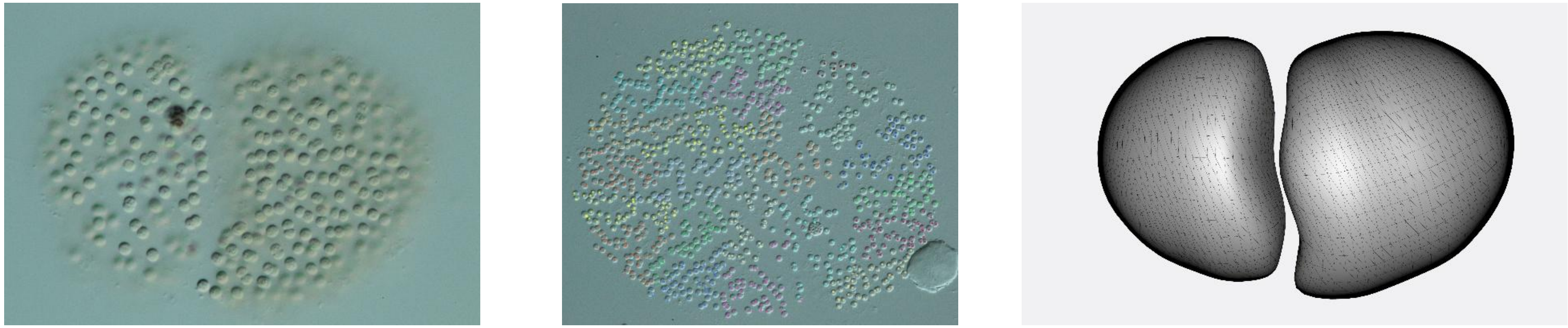

1122

92.

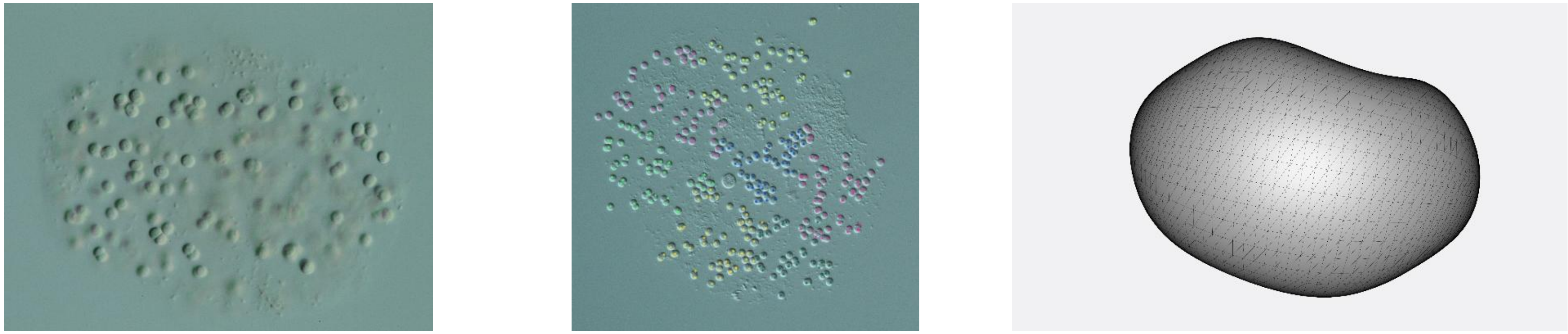

333

93.

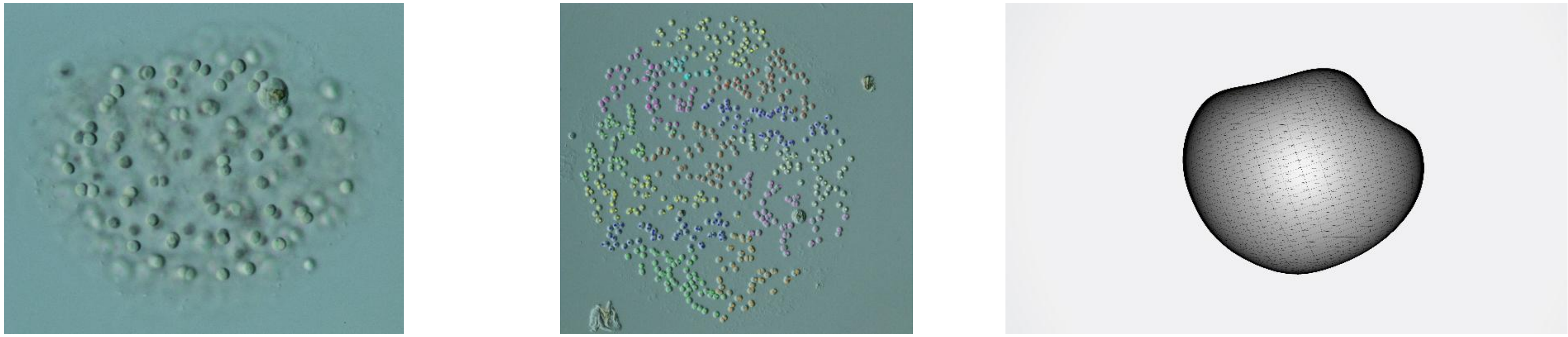

612

94.

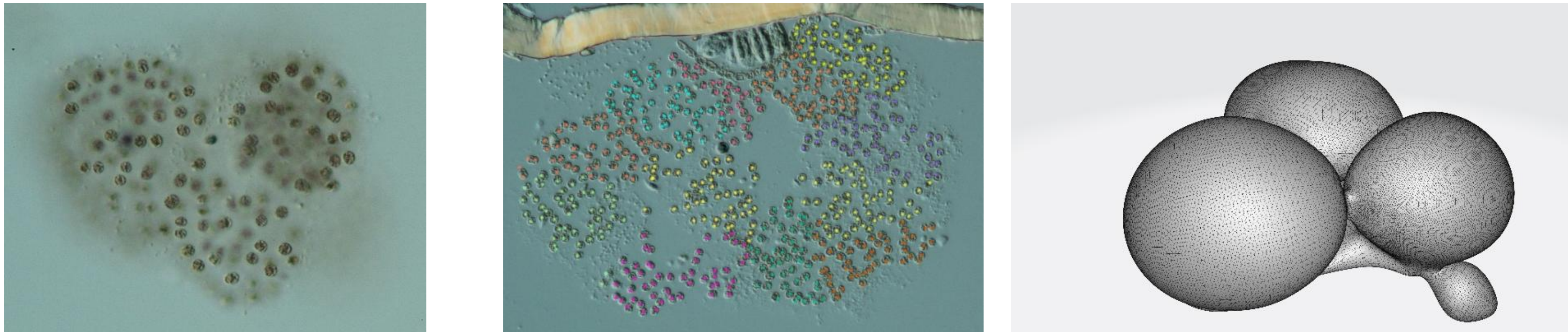

578

95.

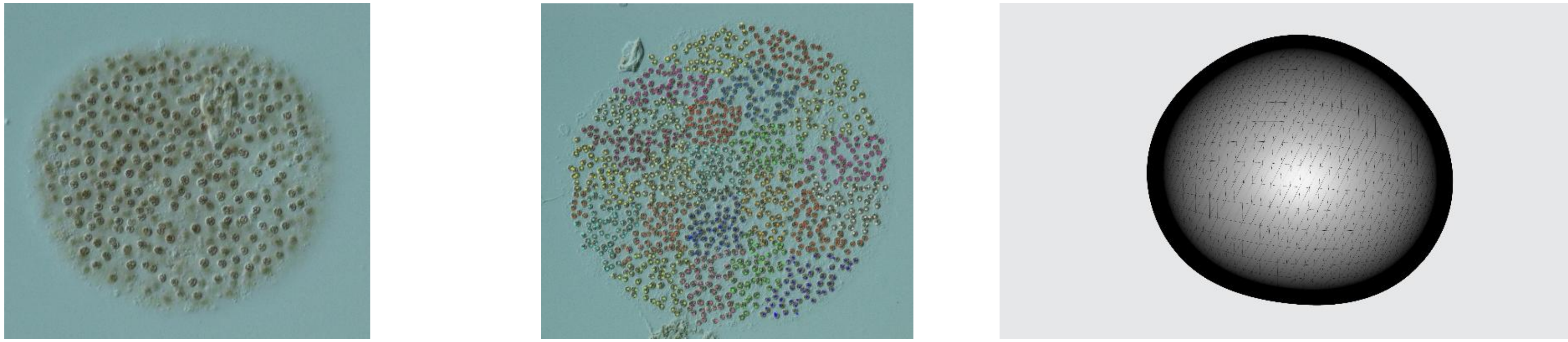

1158

96.

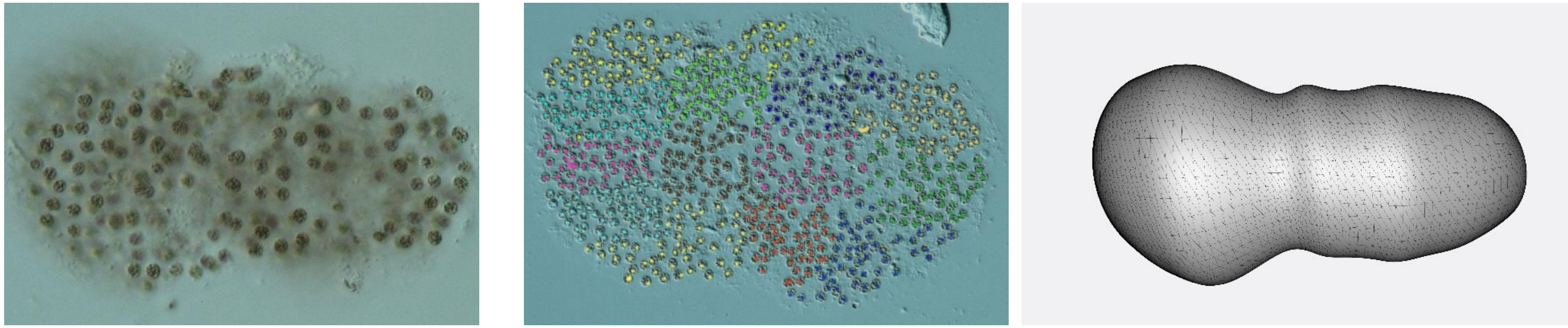

674

97.

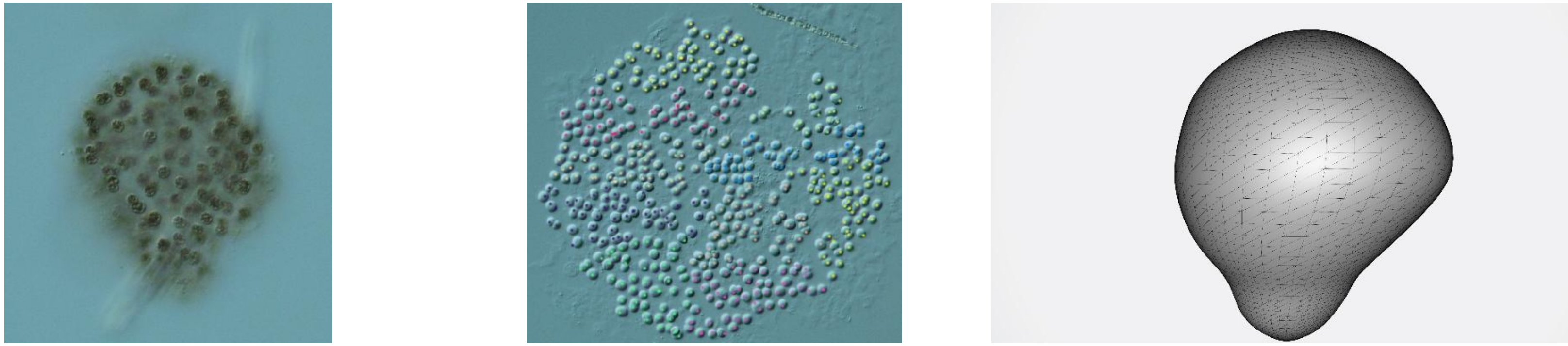

472

98.

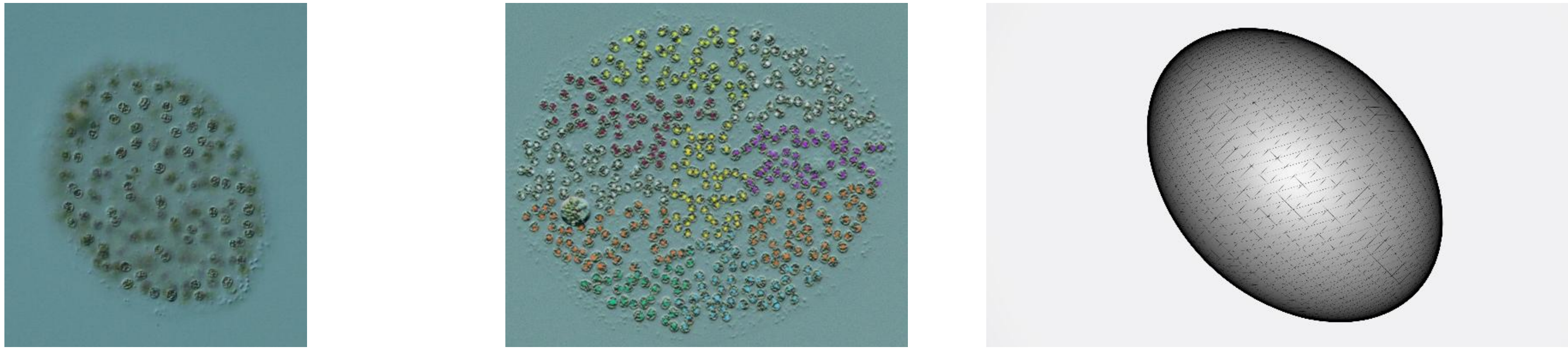

481

99.

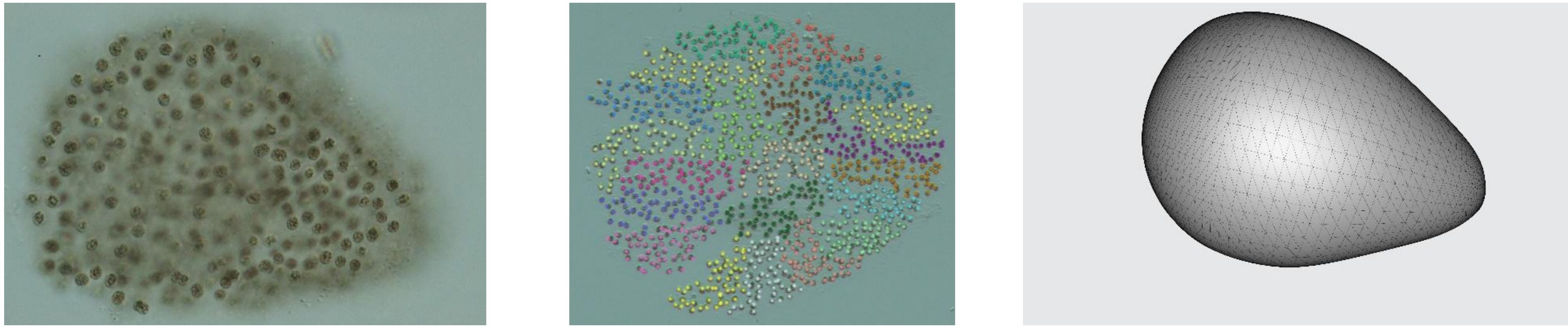

1041

100.

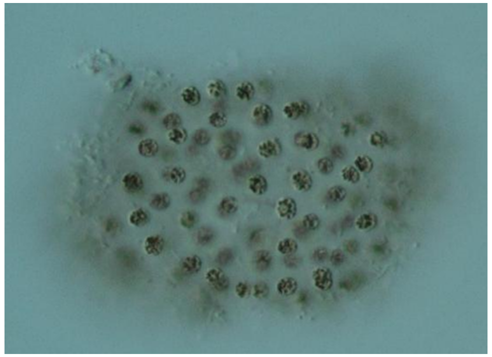

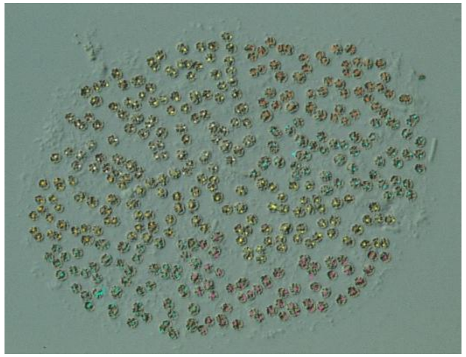

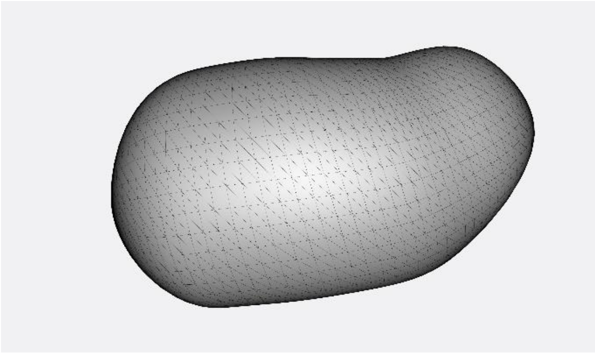

398

105.

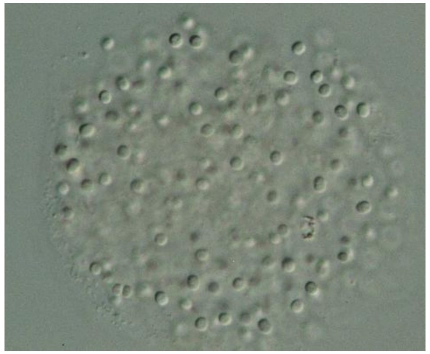

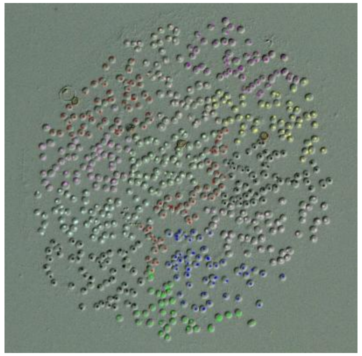

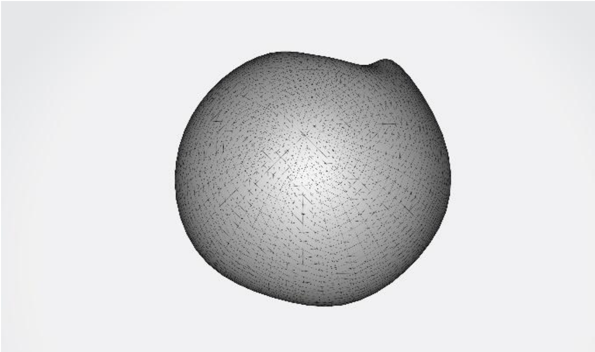

633

114.

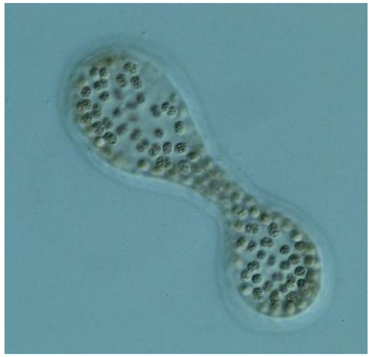

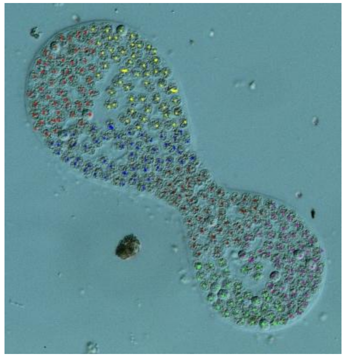

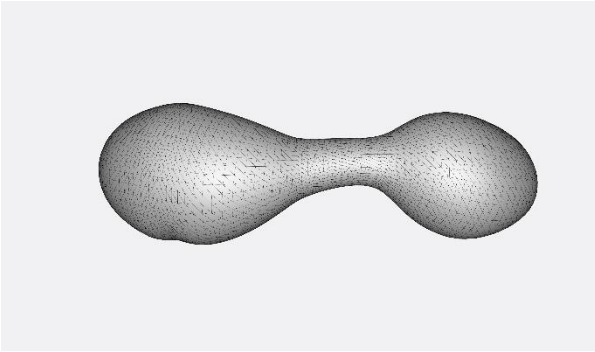

286

117. 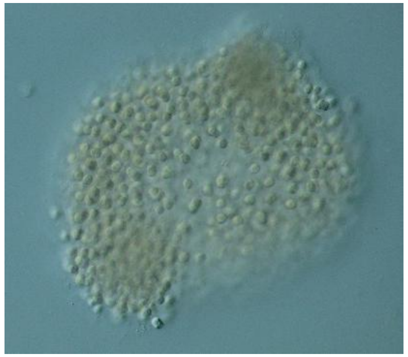 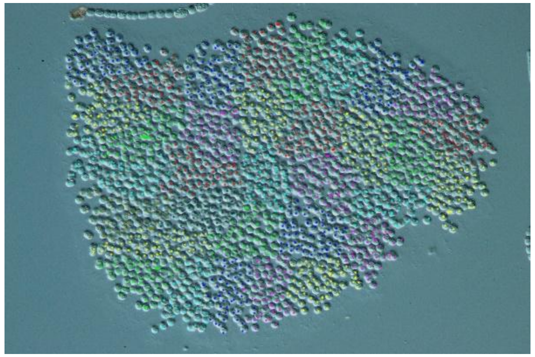 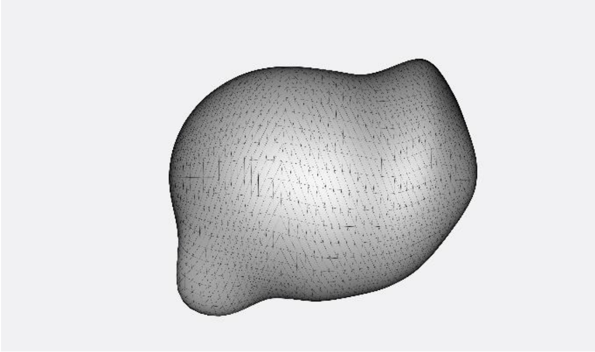 1725

119. 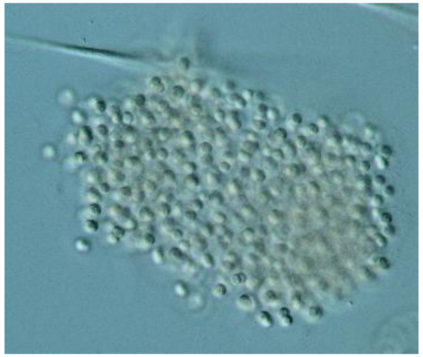 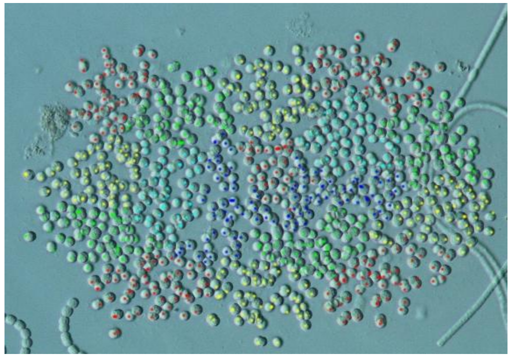 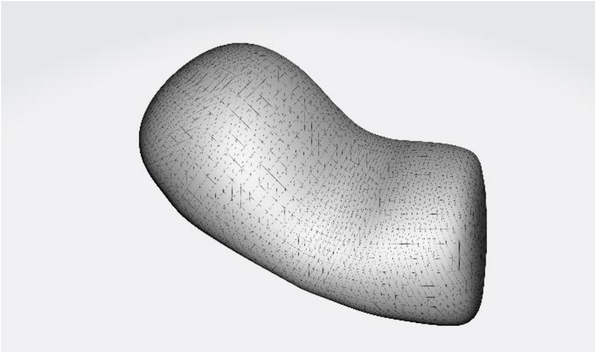 826

133. 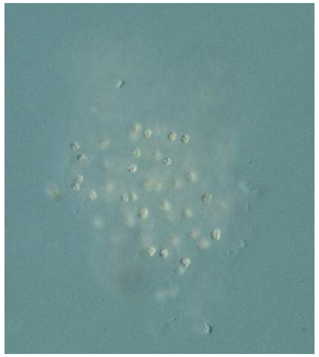 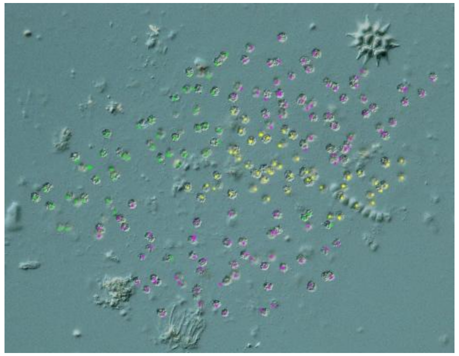 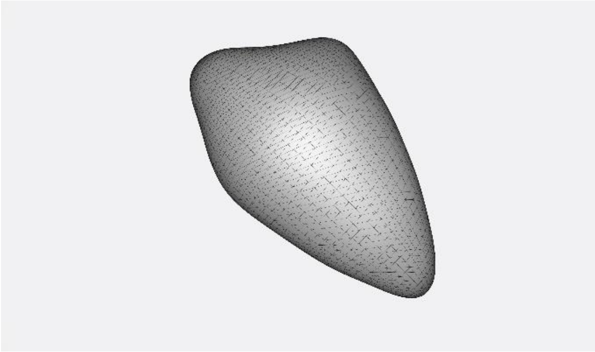 201

143.

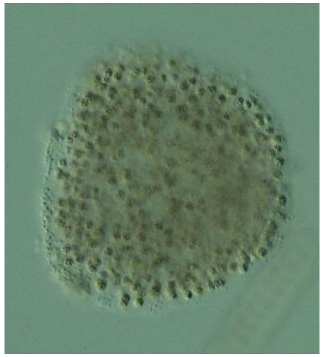

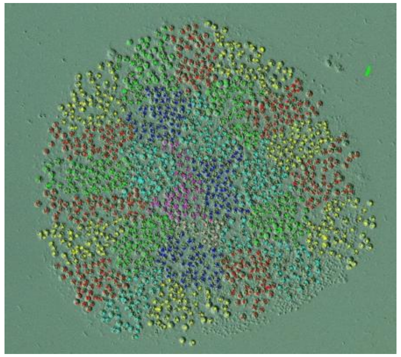

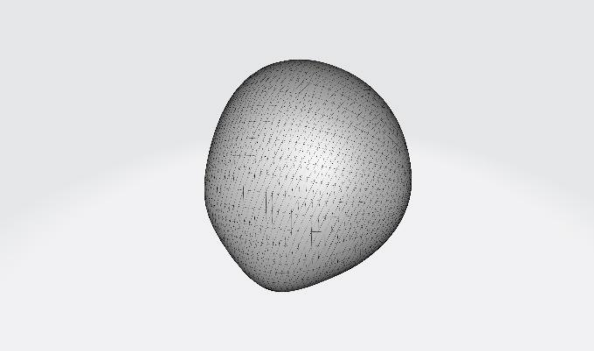

1316

147.

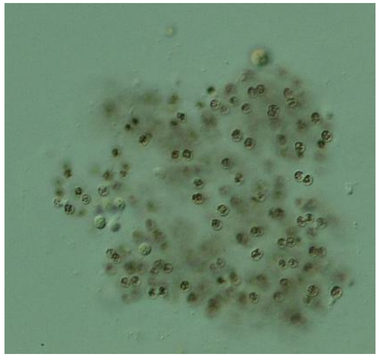

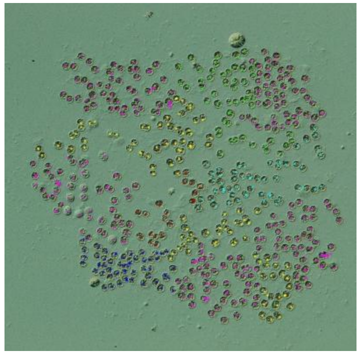

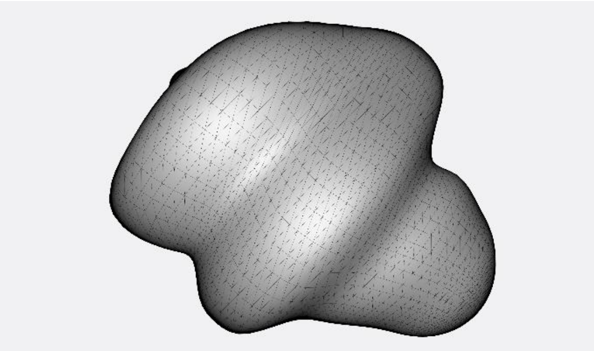

518

155.

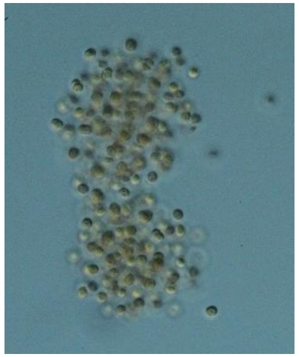

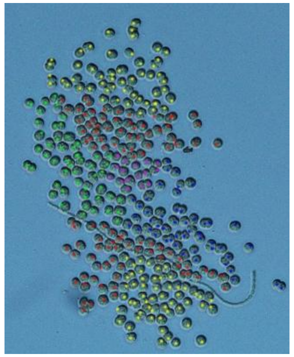

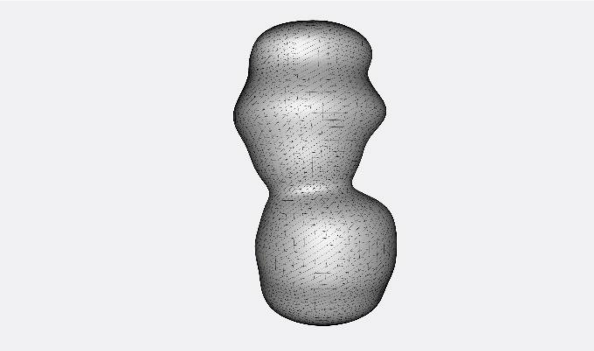

310

161.

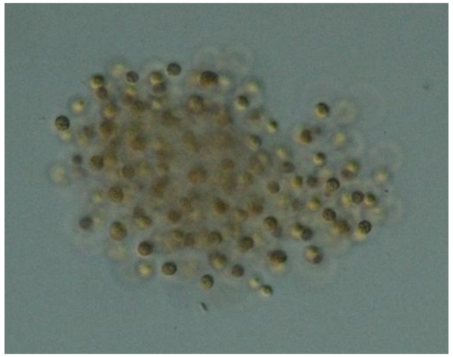

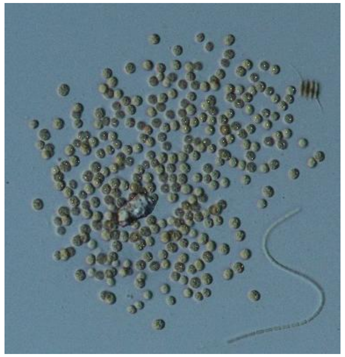

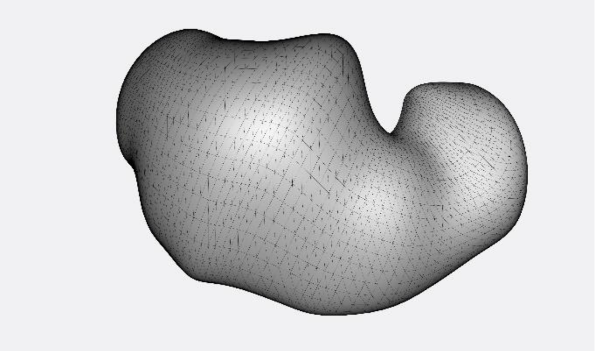

318

171.

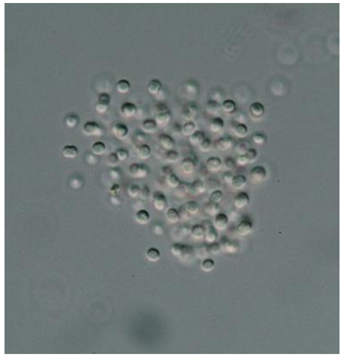

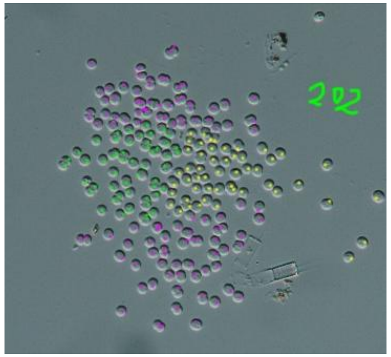


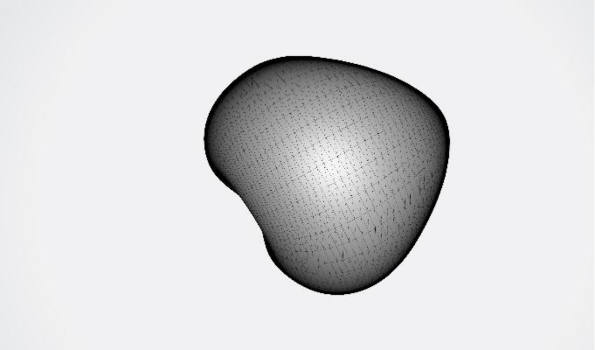

202

172.

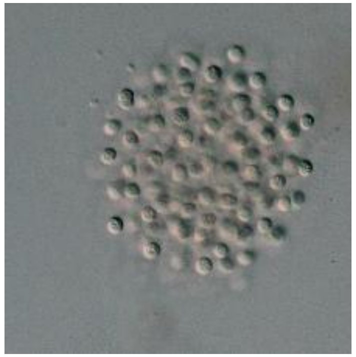

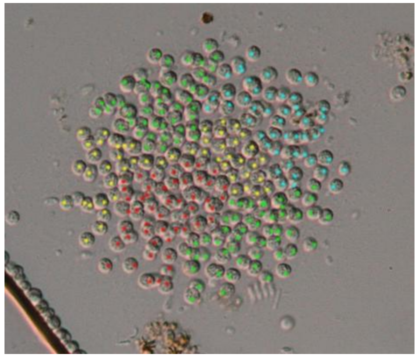

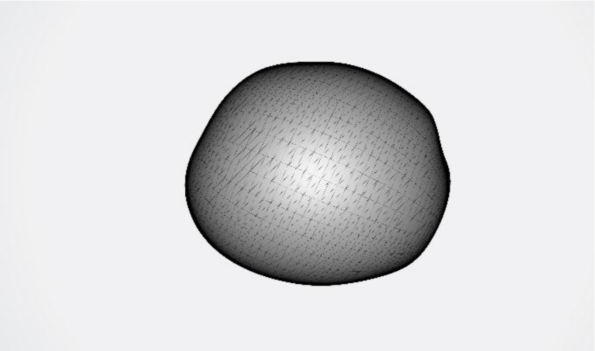

248

173.

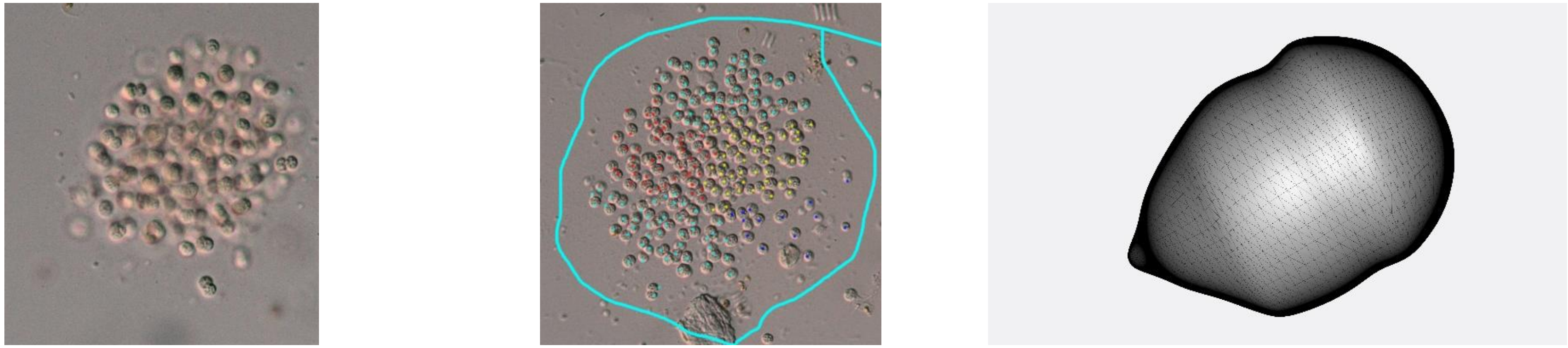

214

176.

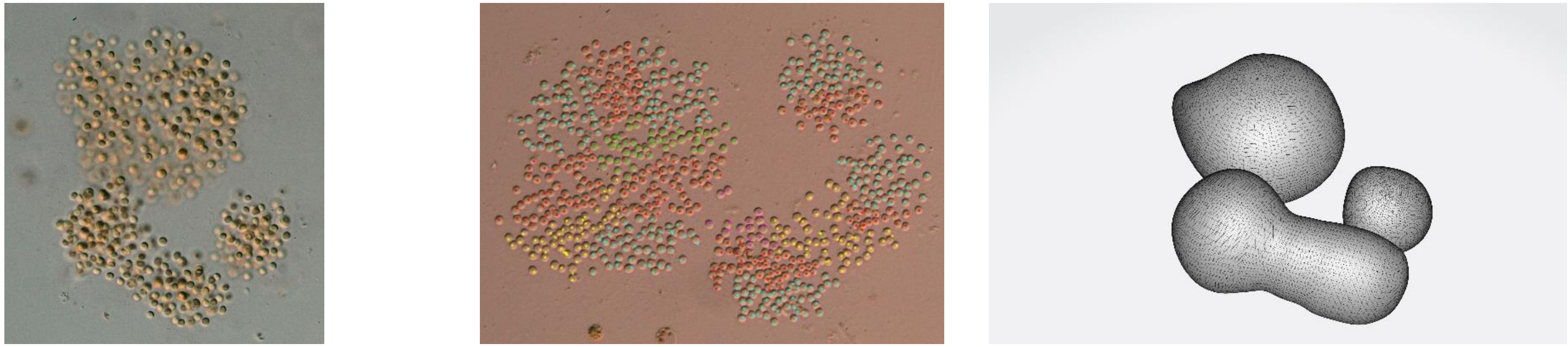

716

193.

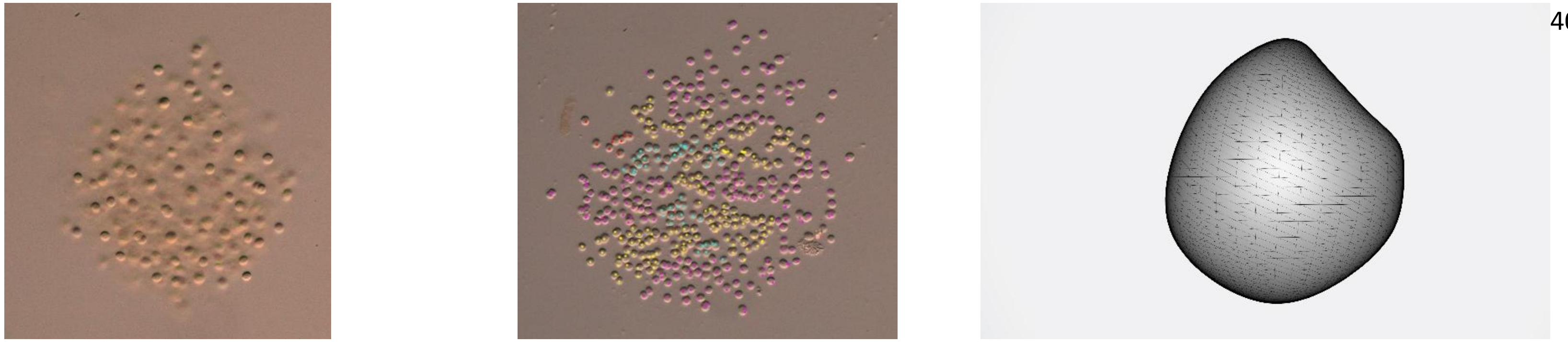

409

198.

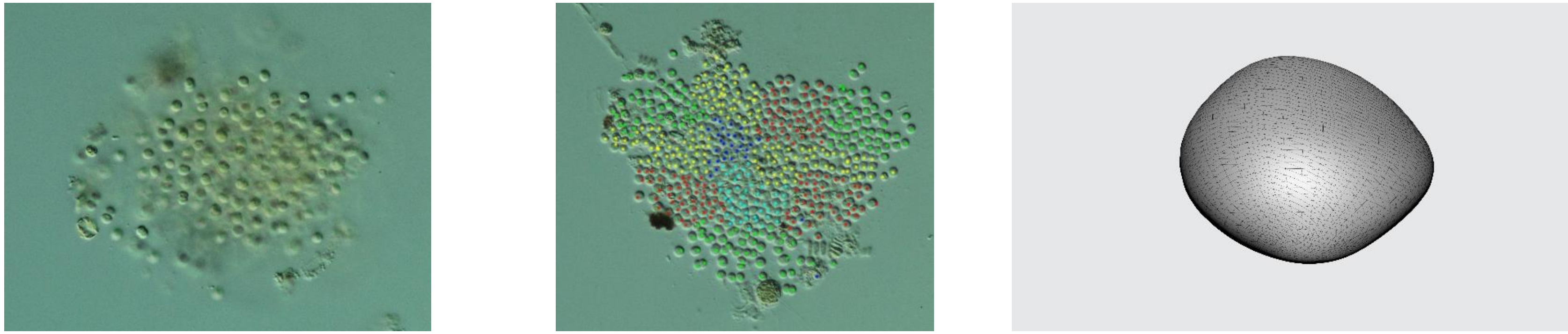

531

199.

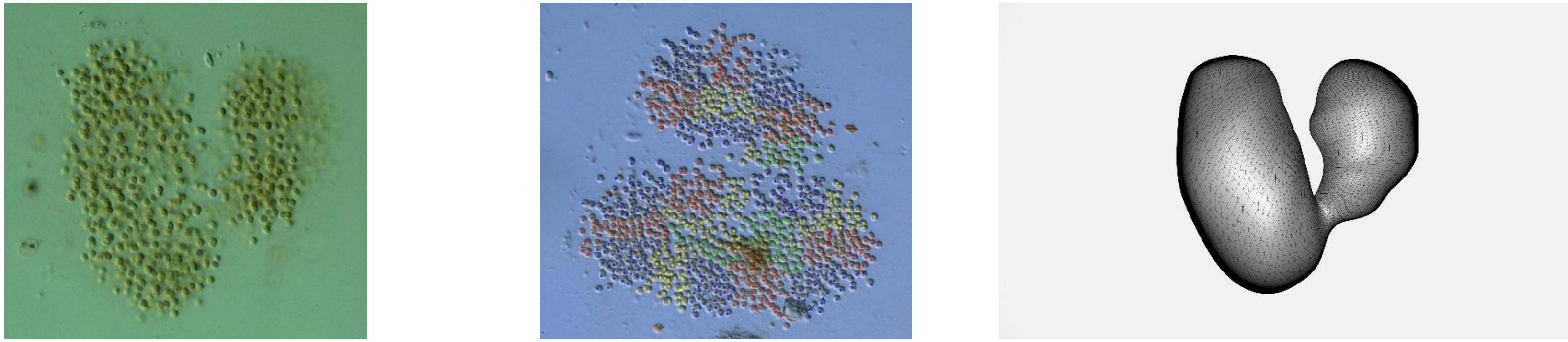

1052
